# Identification of epigenetic regulators of fibrotic transformation in cardiac fibroblasts through bulk and single-cell CRISPR screens

**DOI:** 10.1101/2025.03.28.645873

**Authors:** Laura Pilar Aguado-Alvaro, Nerea Garitano, Wolfgang Esser-Skala, Judy Sayers, Cynthia del Valle, Daniel Alameda-Serrano, Julen Mendieta-Esteban, Maria Erendira Calleja-Cervantes, Ainhoa Goñi-Salaverri, Jon Zazpe, Anna Rosaria de Vito, Francesco Marchese, Diego Alignani, Juliana Cudini, Torsten Gross, Gregorio Rábago, Nisha Narayan, Laura Martinez, Sonia Martinez, Brian Huntly, Paul Riley, Arantxa Gonzalez-Miqueo, Jake P. Taylor-King, Nikolaus Fortelny, Beatriz Pelacho, David Lara-Astiaso

## Abstract

Cardiac fibrosis is mediated by the persistent activity of myofibroblasts, which differentiate from resident cardiac fibroblasts in response to tissue damage and stress signals. The signaling pathways and transcription factors regulating fibrotic transformation have been thoroughly studied. In contrast, the roles of chromatin factors in myofibroblast differentiation and their contribution to pathogenic cardiac fibrosis remain poorly understood. Here, we combined bulk and single-cell CRISPR screens to characterize the roles of chromatin factors in the fibrotic transformation of primary cardiac fibroblasts. We uncover strong regulators of fibrotic states including Srcap and Kat5 chromatin remodelers. We confirm that these factors are required for functional processes underlying fibrosis including collagen synthesis and cell contractility. Using chromatin profiling in perturbed cardiac fibroblasts, we demonstrate that pro-fibrotic chromatin complexes facilitate the activity of well-characterized pro-fibrotic transcription factors. Finally, we show that KAT5 inhibition alleviates fibrotic responses in patient-derived human fibroblasts.

## Main

Cardiac fibrosis is a frequent outcome of many cardiac pathologies and provokes severe clinical complications including arrhythmias and heart failure^1^. The cytokine milieu released upon acute tissue damage activates resident cardiac fibroblasts and induces their differentiation into myofibroblasts, initiating a fibrotic process that enables tissue repair^1,2^. While fibrosis is initially cardioprotective after acute injury, its persistence and expansion leads to adverse tissue remodeling and rigid scar deposition that compromises cardiac function^1^. In addition, reactive interstitial fibrosis in conditions of non-ischemic heart disease also contributes to cardiac function impairment and heart failure progression^3^. TGF-β is the master signaling pathway that orchestrates fibrosis and activates myofibroblast differentiation across different tissues. TGF-β activates Smad transcription factors (TFs)^2^ to induce the expression of collagens, contractile proteins and structural and enzymatic extracellular matrix (ECM) components that mediate fibrotic transformation and adverse tissue remodeling^4^. Single-cell profiling studies have characterized the dynamics of fibroblast states during cardiac repair^5–14^ identifying different molecular mechanisms that regulate fibrotic transformation in different disease conditions. For instance, besides canonical TGF-β signaling, other pathways including non-canonical TGF-β (via Akt/PI3K) and Hippo pathways have been identified as, respectively, positive and negative regulators of fibrotic processes by modulating the assembly of Smad and Tead-YAP1/TAZ TF dimers^15–17^. In addition, a plethora of additional TFs including MRTFs^18^, Egr1/2^19^, AP-1^20^, Meox1^21^ and Runx1^22^ have been shown to cooperate with Smad TFs to drive fibrotic expression programs in the heart and other organs^23^. Identification of the molecular mechanisms that orchestrate fibrosis have enabled the development of experimental anti-fibrotic approaches, namely inhibitors of TGF-β signaling like pirfenidone and renin-angiotensin-aldosterone system modulators^3,24,25^.

Chromatin factors (ChrFs) act in coordination with TFs to regulate transcriptional responses to extracellular stimuli and therefore, are strong determinants of cellular fates^26^. Notably, ChrFs represent good targets for drug discovery since they regulate transcription through catalytic activities that can be easily modulated with chemical approaches^27^. In cardiac fibrosis, small molecule modulation of both chromatin repressors (HDACs and G9a) and activators (Brd4 and Dot1L) has been found to be beneficial^28^. However, a comprehensive understanding of ChrF functions in fibrotic transformation is still missing. Unlike TFs, ChrFs are pervasively expressed and lack specific DNA binding motifs; thus, it is difficult to deconvolve their contribution to fibrosis from single-cell transcriptomic or epigenomic atlases.

To circumvent this limitation, we developed a functional genomics platform to interrogate the contribution of 750 ChrFs to fibrotic transformation in primary murine cardiac fibroblasts. We, then, use Perturb-seq to dissect the functions of the top regulators of fibrotic transformation in primary murine cardiac fibroblasts *ex vivo* and, subsequently characterize their roles using functional assays. Our approach highlights pivotal chromatin complexes, such as SRCAP, Tip60, and NSL, as strong pro-fibrotic mediators. We show that these complexes exert their pro-fibrotic roles by facilitating the activities of fibrotic TFs and demonstrate that the H2AZ acetylation (H2AZac) axis is central in fibrotic responses. Finally, we show that TIP60 (KAT5) inhibition is a potential therapeutic avenue to alleviate fibrosis in patient-derived human cardiac fibroblasts.

## Results

### A FACS-based screen platform to study fibrotic transformation in cardiac fibroblasts ex vivo

FACS-based functional genomic approaches require a clear separation between the interrogated phenotypes, which in our case are naïve and fibrotic cardiac fibroblast states. To develop a CRISPR screen approach to interrogate cardiac fibrosis, we explored culture conditions and FACS readouts in primary cardiac fibroblasts that maximized the separation between resting (unstimulated) and fibrotic (TGF-β-stimulated) fibroblasts (Extended Data Fig. 1a-d). Gene expression analysis of cardiac fibroblasts expanded for 5 days in two different media formulations (serum and serum-free) showed that serum-free conditions not only preserve better the naïve state of resting fibroblasts but also enable a more pronounced fibrotic response after TGF-β stimulation (Extended Data Fig. 1e-f). Therefore, we chose serum-free conditions to expand primary cardiac fibroblasts. Then, using flow cytometry, we examined the dynamics of three potential fibrotic readouts: the fibrotic marker, α-SMA, and two markers of resting states: Sca-1 and PDGFRα^10^. All three markers behaved as predicted with α-SMA levels increasing and Sca-1 and PDGFRα decreasing in response to TGF-β stimulation, however Sca-1 and PDGFRα enabled a better separation of fibrotic (Sca-1/PDGFRα-low) and naïve (Sca-1/PDGFRα-high) fibroblast states (Extended Data Fig. 1g). Downregulation of PDGFRα has been linked to fibrotic progression in different cardiac pathologies^6,8,10,12,29^, thus, we chose PDGFRα as a FACS-readout of fibrotic transformation in our CRISPR screen. To validate this approach, we depleted two central components of TGF-β signaling (*Tgfbr1* and *Smad4*) and measured PDGFRα surface levels after stimulation with the pro-fibrotic signal TGF-β1. In contrast with unperturbed cells, depletion of *Tgfbr1*-KO, *Smad4*-KO in cardiac fibroblasts resulted in high PDGFRα levels after TGF-β stimulation. This indicates that surface PDGFRα is a robust readout to dissect mediators of TGF-β-induced fibrotic transformation (Extended Data Fig. 1h-j).

Next, we generated a CRISPR library targeting 750 epigenetic regulators, selected based on their expression in cardiac fibroblasts across healthy and pathological conditions^10^ (Supplementary Table 1). This library included sgRNAs targeting TGF-β signaling components (*Tgfbr1, Smad3* and *Smad4*) and essential genes (*Eif4a3, Top2a*) as controls. We expanded cardiac fibroblasts isolated from Cas9 mice for 2 days, delivered our CRISPR library and, after 5 days stimulated the cardiac fibroblast cultures with TGF-β for 24h. Then, we FACS-sorted naïve (PDGFRα-high) and fibrotic (PDGFRα-low) cardiac fibroblast states and analyzed their CRISPR library representation using deep sequencing (Fig. 1a). In parallel, we ran a similar experiment using non-Cas9 (wildtype) fibroblasts to correct for library biases (see Methods). Then, we calculated a fibrotic score for each epigenetic regulator that represents the relative enrichment of sgRNAs between naïve (PDGFRα-high) and fibrotic (PDGRFRα-low) fibroblasts. Validating our approach, sgRNAs targeting TGF-β signaling *(Tgfbr1*, *Smad3* and *Smad4)* were enriched in the naïve fibroblast subpopulation but sgRNAs against essential genes (*Eif4a3, Top2a)* were depleted across both naïve and fibrotic populations (Fig. 1b-c). Our analysis identified 35 pro-fibrotic regulators (sgRNAs enriched in the naïve population) and 17 anti-fibrotic factors (sgRNAs enriched in the fibrotic population) (Fig. 1c). Importantly, we found similar patterns for factors belonging to the same chromatin complexes. For instance, several subunits of the SRCAP remodeler (*Srcap, Acrt6, Yeats4* and *Znhit1*), Tip60 H2AZ acetylase (*Kat5* and *Dmap1*) and NSL acetylase^30^ (*Kansl1*, *Kansl*3 and *Kat8)* complexes were nominated as pro-fibrotic regulators (Fig. 1d, Extended Data Fig. 2a). Additional pro-fibrotic regulators were the cofactors *Wdr82* and *Hcfc1*, the NuRD subunit *Chd4;* and the ubiquitin ligases *Rnf20* and *Rnf40*. Conversely, factors belonging to Polycomb-PRC2 (*Ezh2*), non-canonical BAF (*Brd9, Smarcd1, Smarca4*), the H2AZ eraser INO80 (*Ino80*) and MLL-COMPASS (*Paxip1*, *Rbbp5* and *Kmt2a*) complexes scored as anti-fibrotic regulators (Fig. 1d, Extended Data Fig. 2a). Finally, we confirmed 5 pro-fibrotic hits (*Wdr82*, *Srcap*, *Kat8*, *Kat5* and *Hcfc1*) and 4 anti-fibrotic hits (*Ino80*, *Smarca4*, *Brd9* and *Paxip1*) using flow cytometry analysis of PDGFRα surface expression after individual factor depletion in Cas9 primary cardiac fibroblasts (Fig. 1e).

**Figure 1.**
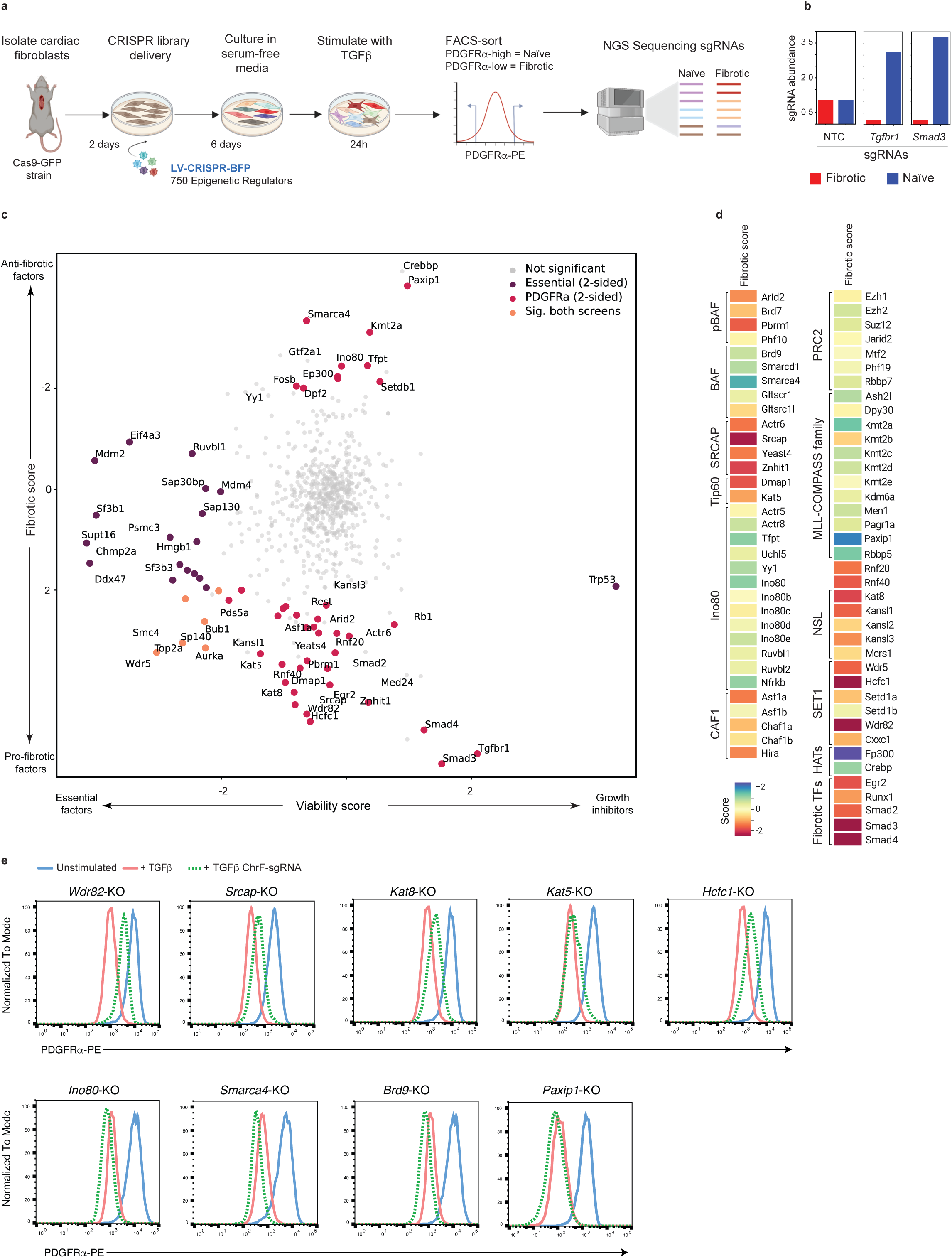
FACS-based CRISPR screen to identify epigenetic regulators of fibrosis in primary cardiac fibroblasts. **(a)** Schematic depiction of the screen system. **(b)** Plot showing abundance of sgRNAs targeting regulators of TGF-β signaling in fibrotic (PDGFRα-low) and naïve (PDGFRα-high) populations. **(c)** 2D plot showing fibrotic scores (y-axis) and viability scores (x-axis) of screen targets. Fibrotic scores derived from sgRNA abundance ratios between PDGFRα-high and PDGRFRα-low populations. Viability scores are derived by comparing non-Cas9 and Cas9 populations (including both PDGFRα-low and high populations). The scores were calculated from 2 screen replicates. Statistical significance for sgRNAs is calculated using 2-sided *t*-tests across two experimental replicates. Gene-level log fold change estimates are established through taking the median across sgRNAs targeting the same gene, and the Fisher method is used to combine *p*-values. The Benjamini-Hochberg false discovery rate is used to calculate corrected *p-*values for the 726 genes tested. **(d)** Heatmap showing fibrotic score of Chromatin Factors grouped by Chromatin Complex membership. **(e)** Representative validations of single candidates using PDGFRα FACS readout of Cas9 fibroblasts depleted for top screen hits grown under TGF-β conditions (n=2-4).

### Perturb-seq dissects regulators of fibrotic and inflammatory states in cardiac fibroblasts

Cardiac fibroblasts are a heterogeneous cell population, whose composition changes dramatically upon myocardial injury^5–14^. We therefore chose to characterize the roles of our top regulators at single-cell resolution. We generated a smaller CRISPR library targeting the top 30 hits from our bulk screen, transduced Cas9 fibroblasts expanded *ex vivo* and used Perturb-seq to characterize single-cell expression patterns of cardiac fibroblasts depleted for these ChrFs under 3 stimulation conditions: resting, IL-1β and TGF-β (Fig. 2a, Extended Data Fig. 2b). These stimuli were chosen as they are central mediators of the early/inflammatory phase of cardiac healing (IL-1β) and the subsequent fibrotic phase (TGF-β)^31,32^.

**Figure 2.**
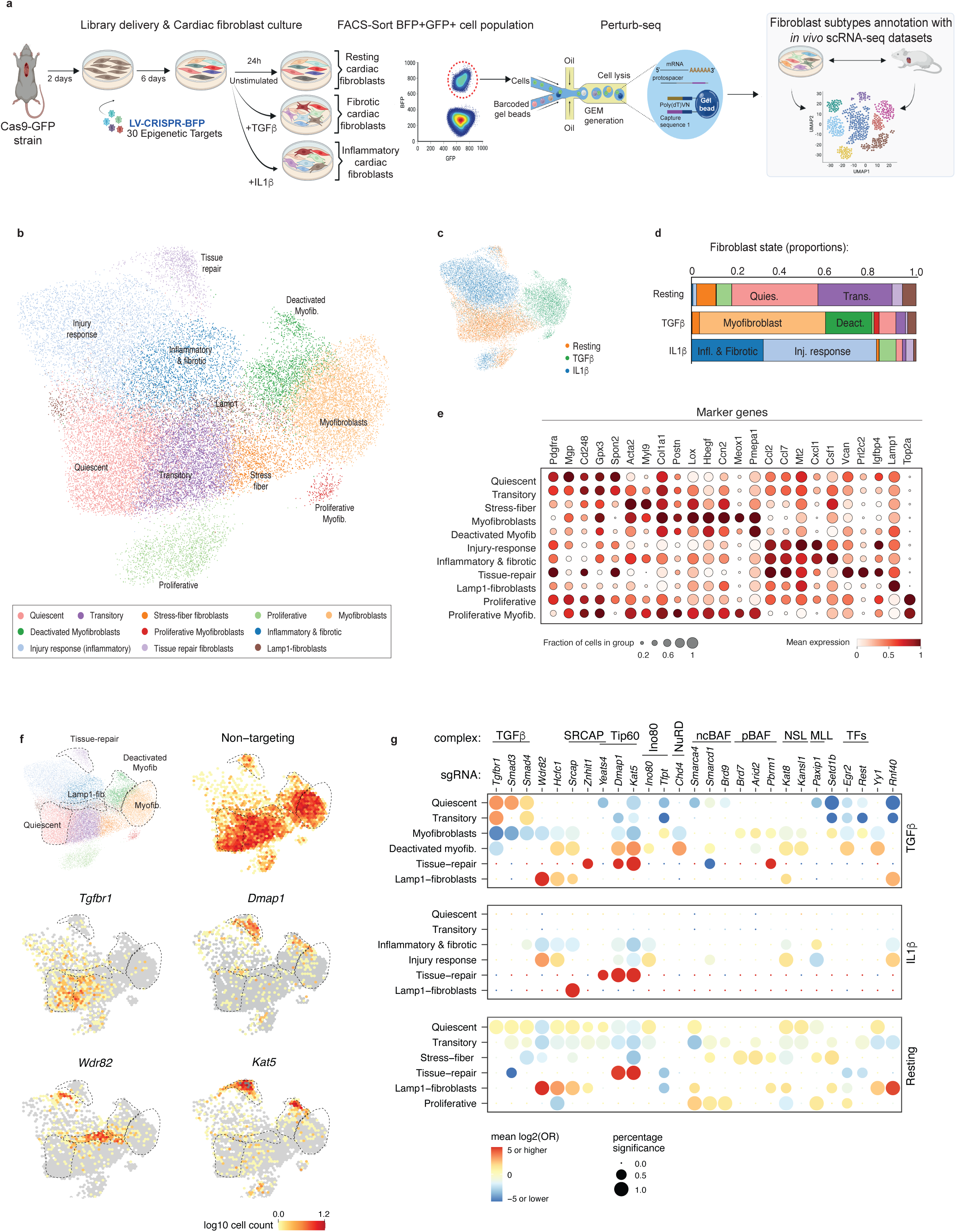
Single-cell perturbation characterization of *ex vivo* fibroblast cultures and analysis of epigenetic regulators in primary fibroblasts. **(a)** Schematic depiction of the *ex vivo* cardiac fibroblast single-cell perturbation analysis approach. **(b)** Integrated UMAP of single-cell transcriptomes derived from primary fibroblasts grown under unstimulated, IL-1β and TGF-β conditions. Fibroblast states are labelled by marker genes and similarity to the fibroblast subpopulations found in murine hearts from available studies^11,13^. **(c)** UMAP projection showing cultured conditions: unstimulated (orange), stimulated with TGF-β (green) and IL-1β (blue). **(d)** Bar plots showing fibroblast subpopulation proportions in the different culture conditions. **(e)** Plot showing expression of fibroblast markers across fibroblast subpopulations. Resting markers: *Pdgfra*, *Mgp*, *Cd248*, *Gpx3, Spon2*. Fibrotic markers: *Acta2*, *Myl9*, *Col1a1*, *Postn*, *Lox*, *Hbegf*, *Ccn2, Meox*1. Inflammatory markers: *Ccl2, Ccl7, Mt2, Cxcl1, Csf1*. Pro-repair genes: *Vcan*, *Prl2c2, Igfbp4*. Proliferating: *Top2a*. Dot size represents percentage of cells in the cluster expressing the marker gene. Color represents scaled expression values. **(f)** UMAPs showing the distribution of unperturbed fibroblasts (non-targeting) and *Tgfbr1*, *Dmap1*, *Wdr82* and *Kat5* perturbed fibroblasts. **(g)** Enrichment analysis of cells with specific epigenetic perturbations across the different fibroblast states identified in TGF-β, IL-1β and resting conditions. Dot color and size relate to the log2 odds ratio and the percent of significant enrichments (one test was performed per NTC), respectively. The analysis is based on measurements of two merged sgRNAs per target.

Resting expansion conditions produced two major fibroblast states: a naïve state (Quiescent and Transitory) expressing steady-state fibroblast markers including *Pdgfra*, *Cxcl14*, *Gpx3*, *Junb* and *Cd248* and, a fibroblast population with high expression of contractile components (*Acta2, Tpm2, Tagl2, Myl9*), which we called “Stress-fiber fibroblasts” (Fig. 2b-e). Stimulation with TGF-β induced myofibroblast differentiation, which expressed high levels of typical myofibroblast markers (*Acta2*, *Hbegf*^33^, *Ccn2*^34^), collagen biosynthesis factors (*Col1a*, *Lox*, *Loxl2*) and the pro-fibrotic TF, *Meox1* (Fig. 2b-e, Extended Data Fig. 2c). IL-1β stimulation induced two fibroblast phenotypes: “Injury response” with high levels of inflammatory mediators (*Ccl2*, *Ccl7, Cxcl1*) and “Inflammatory & fibrotic” with joint expression of inflammatory and fibrotic genes. Notably, both IL-1β subpopulations expressed molecules mediating the initial stages of cardiac healing such as osteopontin (*Spp1*), decorin (*Dcn*)^35^ and methalothioneins (*Mt1* and *Mt2*)^36^. To better characterize our *ex vivo* system, we cross-referenced the *ex vivo* fibroblast expression signatures with the signatures of cardiac fibroblast subpopulations found in *in vivo* single-cell maps of murine hearts in healthy and fibrotic conditions^11,13^ (Extended Data Fig. 2d-e). This approach demonstrated similarities between our *ex vivo* states and their *in vivo* counterparts, especially between *in vivo* pathological fibrotic states and *ex vivo* myofibroblasts. Finally, we used SCENIC^37^ to identify the transcriptional regulators of each of the different *ex vivo* subpopulations. Bhlhe41, Foxc2 and Egr2 were nominated as myofibroblast drivers and, Fosl1 and Nf-kb (Relb, Nfkb1, Nfkb2) were identified as inflammatory drivers^38^ (Extended Data Fig. 3).

Besides the subpopulations described above (naïve, fibrotic and inflammatory), we found three smaller fibroblast subpopulations, whose abundance increased after specific ChrF perturbations (Fig. 2f-g). TGF-β conditions contained a population with attenuated expression of fibrotic markers (*Acta2*, *Col1a1*, *Lox*) and high levels of the TGF-β repressor *Pmepa1*^39^, which we called “Deactivated myofibroblasts”. Additionally, all three conditions contained a population of “Tissue-repair fibroblasts” with high levels of anti-fibrotic (*Spon2, Sfrp1, Tgfbr3*)^40–42^, pro-repair and angiogenic (*Prl2c2-3*, *Fgf7, Nid1*) factors^43,44,45^ and, a subpopulation characterized by the expression of the autophagic marker, *Lamp1* (Fig. 2g, Extended Data Fig. 2c).

To quantitatively measure the effect of ChrF perturbations on the dynamics of fibroblast populations in response to stimuli, we analyzed the distribution of each perturbation across fibroblast subpopulations and stimulation conditions using non-targeting control (NTC) cells as a background distribution (Fig. 2f, Extended Data Figs. 4-5). As expected, depletion of TGF-β signaling components (*Tgfbr1*-, *Smad3*- and *Smad4*-KO) prevented myofibroblast transformation in response to TGF-β, with perturbed cells retained in the naïve fibroblast state (Fig. 2g). Perturbations of Tip60, SRCAP, NuRD, NSL, *Wdr82* and *Hcfc1* also reduced the myofibroblast subpopulation. However, instead of preserving the naïve state, these ChrF perturbations pushed fibroblasts towards the deactivated myofibroblast state mentioned above. This suggests that these epigenetic regulators are required to enable a full fibrotic response after TGF-β stimulation (Fig. 2g, Extended Data Fig. 6a). Finally, analysis of IL-1β conditions identified *Paxip1* and Tip60 components (*Kat5*-KO and *Dmap1*-KO) as mediators of IL-1β-driven inflammatory response in cardiac fibroblasts (Fig. 2g, Extended Data Fig. 6a). Interestingly, depletion of Tip60 members (*Kat5*-KO and *Dmap1*-KO) abrogated both fibrotic and inflammatory responses and markedly enriched the “Tissue-repair fibroblast” state, which was practically absent in unperturbed fibroblasts (Fig. 2f-g). Similarly, *Wdr82* and *Rnf40* depletion induced the “Lamp1-fibroblast” state. Collectively, these results highlight how different chromatin complexes regulate specific fibroblast states in response to stimuli.

### Chromatin factor complexes regulate fibrotic functions

We then decided to characterize the functions of the top fibrotic regulators identified above in detail: the Tip60 components *Kat5* and *Dmap1*, the SCRAP complex subunit *Srcap*, the NSL catalytic component *Kat8*, and the co-factors *Wdr82* and *Hcfc1*. To this end, we examined the effect of these perturbations on the expression of genes involved in collagen biosynthesis, stress-fiber formation and ECM deposition, which are three pivotal fibrotic processes (Fig. 3a). All ChrF perturbations had a clear impact on the expression of fibrotic mediators but each selectively. For instance, whereas *Wdr82* was highlighted as a key mediator of collagen biosynthesis, Tip60 and *Srcap* were nominated as top regulators of stress-fiber and ECM deposition. Additionally, in line with the subpopulation enrichment analysis, depletion of *Kat5* and *Dmap1* increased the levels of anti-fibrotic factors (*Prg4)* and pro-repair molecules including versican (*Vcan*)^46^, proliferins (*Prl2c2-3*)^43^ and *Igfbp4*^47^ (Fig. 3a). Finally, gene set enrichment analysis confirmed the pro-inflammatory role of *Paxip1* and highlighted additional cellular functions regulated by these chromatin complexes such as the regulation of mitochondrial activity by *Kat8*, or retinol metabolism by the pBAF subunits *Pbrm1* and *Brd7* (Extended Data Fig. 6b).

**Figure 3.**
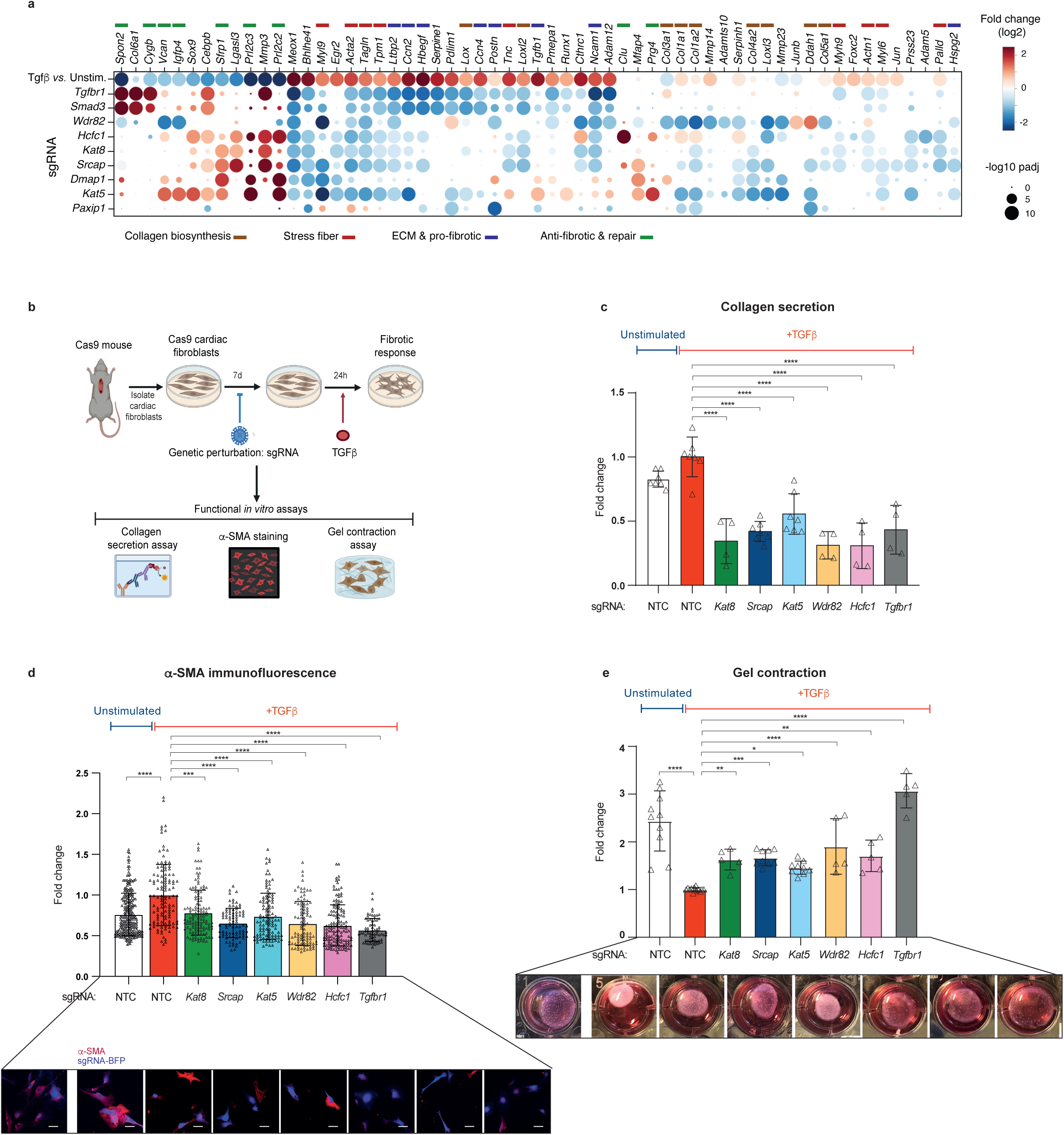
Kat8, Srcap, Kat5, Wdr82 and Hcfc1 are required for pivotal fibrotic cellular functions. **(a)** Effect of epigenetic perturbations on the expression of genes mediating fibrotic functions. The color of each dot represents the log2 fold change (compared to NTCs), the size represents the –log10 adjusted p-value, as per key to the bottom right. **(b)** Schematic depiction of fibrotic assays. **(c)** Collagen secreted levels in control fibroblasts (NTC) and in fibroblasts depleted for *Kat8*, *Srcap*, *Kat5, Wdr82, Hcfc1* and *Tgfbr1.* Collagen levels were determined using an ELISA assay. The measurements were performed in four-seven replicate experiments. **(d)** Quantification of the immunofluorescence levels of α-SMA in control fibroblasts (NTC) and in fibroblasts depleted for *Kat8*, *Srcap*, *Kat5, Wdr82, Hcfc1* and *Tgfbr1.* The measurements were derived from two replicate experiments; a total number of ∼100 cells were quantified per condition. Representative images of α-SMA immunostaining assays are shown at the bottom. Scale bars: 7 μm. **(e)** Quantification of the contractile capability of control fibroblasts (NTC) and fibroblasts depleted for *Kat8*, *Srcap*, *Kat5, Wdr82, Hcfc1* and *Tgfbr1*. The measurements were performed in five-nine replicate experiments. Representative images of collagen gel contraction are shown at the bottom. All data are shown as fold change *vs.* NTC+TGF-β values and are mean ± SD. Statistical significance was analyzed by one-way ANOVA or Kruskal-Wallis tests and is indicated as follows: *P <0.05, **P<0.01, ***P<0.001, ****P <0.0001.

Next, we decided to link the transcriptional effects of ChrF perturbation with three functional processes mediated by fibrotic fibroblasts: collagen deposition, stress-fiber formation and contractility^4,48^ (Fig. 3b, Extended Data Fig. 7a-b). Depletion of *Kat5*, *Srcap*, *Kat8*, *Wdr82* and *Hcfc1* provoked a marked decrease in secreted collagen levels in cardiac fibroblasts stimulated with TGF-β. In line with the transcriptional signatures described above, *Wdr82*-KO fibroblasts showed the strongest reduction in collagen secretion (3-fold) (Fig. 3c). Similarly, the levels of the contractile protein α-SMA and the contractile ability of cardiac fibroblasts (TGF-β stimulated) were reduced upon ChrFs perturbation, with *Srcap*-KO fibroblasts showing the strongest effects for this functional readout (Fig. 3d-e). Of note, none of the analyzed perturbations compromised the proliferation capacity of fibroblasts or induced apoptosis. Thus, the observed decrease in fibrotic functions is likely not a consequence of a decrease in cellular fitness (Extended Data Fig. 7 c-d). Taken together, our results demonstrate that SRCAP, Tip60, NSL chromatin complexes and *Hcfc1* and *Wdr82* cofactors mediate fibrotic functions in cardiac fibroblasts with their perturbation leading to attenuated fibrotic responses.

### SRCAP, Tip60 and NSL complexes and Wdr82 regulate chromatin accessibility of pro-fibrotic transcription factors

To explore the molecular mechanisms underlying the pro-fibrotic roles of our top epigenetic regulators, we analyzed the chromatin accessibility patterns of cardiac fibroblasts depleted for *Srcap*, *Kat5*, *Kat8* and *Wdr82* after TGF-β stimulation. As shown in previous studies^49^, TGF-β stimulation led to marked changes in chromatin accessibility in unperturbed cells (3,201 and 1,290 peaks with increased and decreased accessibility, respectively) with a strong induction of chromatin accessibility at fibrotic loci (Fig. 4a, Extended Data Fig. 8a-b). In line with their pro-fibrotic roles, depletion of *Srcap, Kat5* and *Wdr82* led to a prominent decrease in chromatin accessibility at TGF-β-responsive loci located in the vicinity of key pro-fibrotic regulators including *Acta2*, *Postn*, *Col1a1*, *Loxl1, Mmp13* and *Meox1* (Fig. 4a, Extended Data Fig. 8a). Overall, *Srcap* and *Wdr82* depletion had the largest effects on global chromatin accessibility with respectively 3,336 upregulated and 3,316 downregulated peaks for *Srcap*-KO and 4,985 upregulated and 4,200 downregulated peaks for *Wdr82*-KO (Extended Data Fig. 8b).

**Figure 4.**
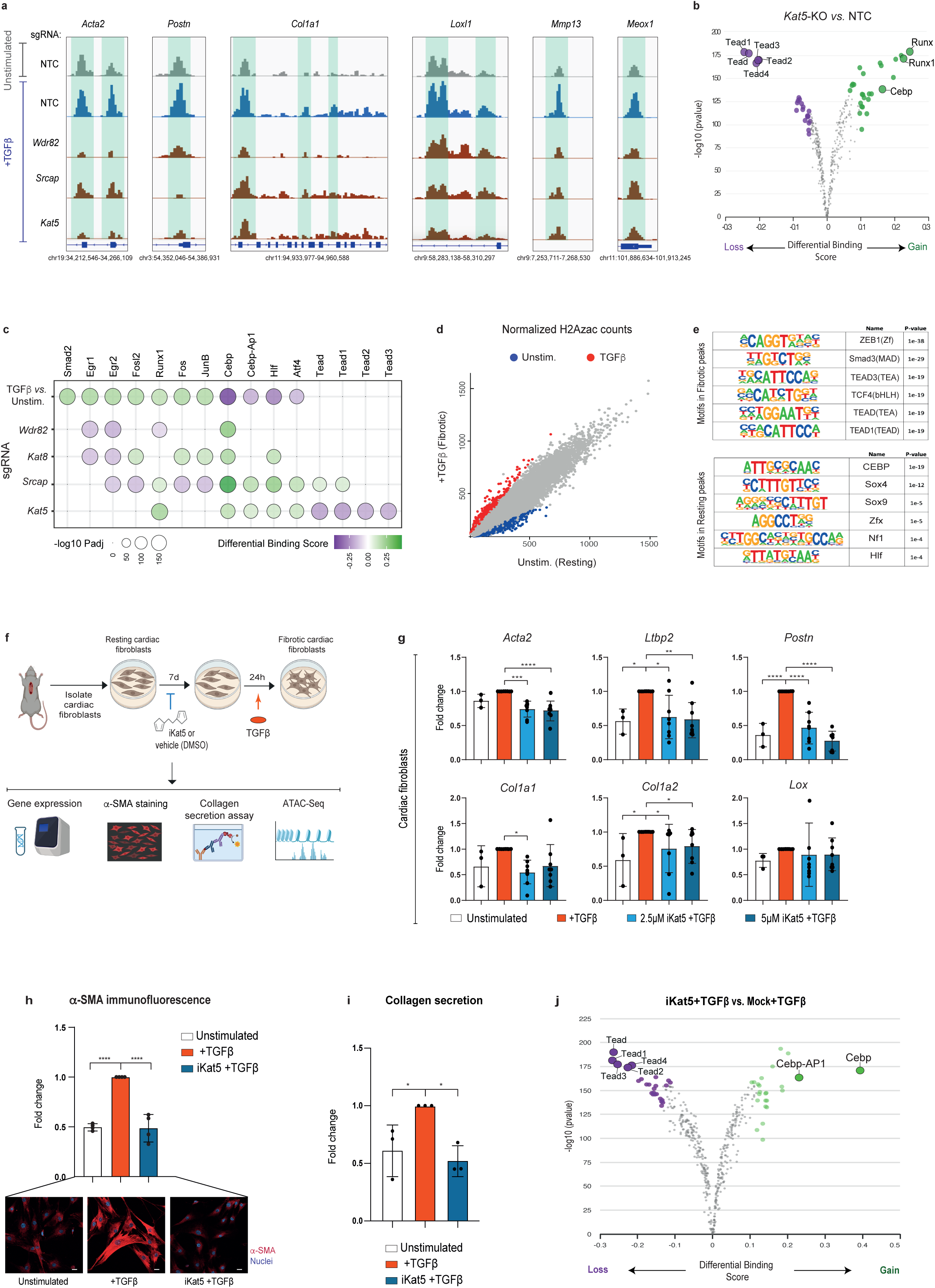
Epigenetic mechanisms underlying the pro-fibrotic roles of Kat8, Srcap, Kat5 and Wdr82. **(a)** Genome browser snapshots of ATAC-seq signal for fibrotic loci in cardiac fibroblasts: unstimulated (grey), 24h TGF-β-stimulated (blue) and *Wdr82*, *Srcap* and *Kat5* knockouts in TGF-β stimulated conditions (red). **(b)** Volcano plot showing differential TF motif footprints between conditions in TGF-β-stimulated (24h) conditions: *Kat5* knockout *vs.* control fibroblasts (NTC). Differential footprinting analysis was performed with TOBIAS. **(c)** Heatmap summarizing the effect of *Wdr82*, *Srcap, Kat8* and *Kat5* on TF motif accessibility in TGF-β-stimulated (24h) conditions. Dot color and size relate to log2 fold change and the -log10 adjusted p-value, respectively. **(d)** Scatter plot comparing H2AZac signal between unstimulated and TGF-β-stimulated (24h) fibroblasts. Red dots mark loci with increased H2AZac signal (log2FC > 0.75, padj <0.01) after TGF-β-stimulation. Blue dots label loci with decreased H2AZac signal (log2FC < -0.75, padj <0.01) after TGF-β stimulation. Axis values are normalized ChIP-seq counts. **(e)** Top TF motifs enriched in the TGF-β-stimulated (fibrotic) and unstimulated H2AZac differential peaks. Motif enrichment was computed with Homer using as background the entire H2AZac peak repertoire. **(f)** Schematic depiction of the experimental approach in murine cardiac fibroblasts. **(g)** Expression of fibrotic marker genes following TGF-β stimulation in control and NU-9056 pre-treated murine cardiac fibroblasts. The measurements were performed in three-eight replicate experiments. **(h**) Quantification of the average immunofluorescence intensity of α-SMA, following TGF-β stimulation in control and NU-9056 pre-treated murine cardiac fibroblasts. The measurements were performed in four replicate experiments. Representative images of α-SMA immunostaining assays are shown at the bottom. Scale bars: 20 μm. **(i)** Quantification of collagen secretion following TGF-β stimulation in control and NU-9056 pre-treated murine cardiac fibroblasts. The measurements were performed in three replicate experiments. **(j)** Differential TF-footprinting analysis between control and NU-9056 pre-treated murine fibroblasts after TGF-β stimulation. Dots in purple represent TF motifs with decreased accessibility in NU-9056 pre-treated murine fibroblasts. All data are shown as fold change *vs.* TGF-β values and are mean ± SD. Statistical significance was analyzed by one-way ANOVA or Kruskal-Wallis tests and is indicated as follows: *P <0.05, **P<0.01, ***P<0.001, ****P <0.0001.

Chromatin complexes cooperate with TFs to regulate cellular states and differentiation processes. Therefore, we decided to use differential TF footprinting analysis to identify the TF partners of the studied ChrFs that jointly mediate the fibrotic response to TGF-β stimulation. Analysis of TGF-β stimulated *vs.* resting conditions in unperturbed fibroblasts highlighted a marked increase in motif accessibility of Smad2, Egr1, Egr2, Fosl2, Runx1 and Fra2, which are TFs involved in fibrotic transformation across different cellular contexts^2,18,19,21,50^ (Extended Data Fig. 8c). Depletion of *Wdr82*, *Kat8* and *Srcap* significantly reduced the motif accessibility of several of these pro-fibrotic TFs, with reduction of Egr2 motif accessibility common to *Wdr82*, *Kat8* and *Srcap* perturbations (Fig. 4b-c, Extended Data Fig. 8c-f). Confirming this trend, genetic perturbation of *Egr2* in cardiac fibroblasts maintains high PDGFRα levels and reduces the expression of fibrotic genes after TGF-β stimulation (Extended Data Fig. 8g-h). *Kat5* depletion produced a very different pattern, with a specific downregulation of the motif accessibility of Tead TFs (Fig. 4b-c). Tead TFs mediate fibrosis via YAP signaling^51,52^ suggesting that Kat5 regulates fibrosis by using alternative molecular pathways than *Wdr82*, *Kat8* and *Srcap*. Interestingly, all four ChrF knockouts increased the accessibility of the Cebp motif, downregulated in unperturbed cells after TGF-β stimulation (Fig. 4b-c, Extended Data Fig. 8c-f). Of note, Cebpa and Cebpd TFs mediate anti-fibrotic roles on other tissues^53,54^ and Cebpd was nominated as a top regulator of quiescent fibroblasts in our SCENIC analysis (Extended Data Fig. 3). This suggests that the depletion of our four pro-fibrotic ChrFs increases the anti-fibrotic activity of Cebp TFs.

The main catalytic activity of Tip60 complex is the acetylation of the H2AZ histone variant^55^. To examine the relationship between H2AZac deposition and fibrosis, we analyzed H2AZac levels in primary cardiac fibroblasts under resting and TGF-β stimulation conditions. This analysis revealed a marked increase in H2AZac at pro-fibrotic loci (*Col1a1*, *Loxl1*, *Acta2*, *Palld*, *Postn*, *Myl9* and *Cthrc1*) and an enrichment for Smad and Tead motifs at TGF-β responsive loci (Fig. 4d-e, Extended Data Fig. 8i). This provides additional evidence for the synergistic interaction between Tead TFs and the Tip60 complex during fibrotic responses. Finally, to confirm the requirement of Kat5 H2AZac activity in fibrotic transformation, we used a Kat5 chemical inhibitor (NU-9056) to block H2AZac in cardiac fibroblasts and measured fibrotic responses after TGF-β stimulation using gene expression and functional readouts (Fig. 4f). Inhibition of Kat5 acetylase activity provoked a marked decrease in fibrotic gene expression, α-SMA protein levels and collagen deposition (Fig. 4g-i, Extended Data Fig. 8j). Importantly, we also observed the anti-fibrotic effect of *Kat5* inhibition in fibroblasts stimulated with TGF-β for 24h and subsequently treated with NU-9056, indicating that *Kat5* inhibition may lead to the reversion of fibrotic phenotypes (Extended Data Fig. 8k). Furthermore, chemical inhibition of Kat5 recapitulated the decrease in Tead motif chromatin accessibility observed in genetically perturbed (*Kat5-KO*) cardiac fibroblasts (Fig. 4j). These results confirm that the pro-fibrotic activity of Kat5 is mediated by its acetylase activity.

### Kat5 is a pro-fibrotic mediator across different tissues

Next, we decided to interrogate whether Kat5 is a universal mediator of fibrosis across different organs or if its pro-fibrotic function is restricted to cardiac fibroblasts. To this end, we isolated fibroblasts from lung, skin and kidney, treated them with Kat5 inhibitor and measured their response to TGF-β stimulation by measuring gene expression, α-SMA protein levels and collagen deposition (Fig. 5a, Extended Data Fig. 9a-c). Treatment with Kat5 inhibitor reduced fibrotic transcriptional responses in fibroblasts derived from lung, skin and kidney, with lung fibroblasts showing the strongest effects (Fig. 5b-d). Similarly, α-SMA protein levels were decreased after Kat5 inhibition across all 3 tissues (Fig. 5e-g). Finally, collagen secretion was markedly reduced in lung fibroblasts after Kat5 inhibition but only mildly decreased, albeit without statistical significance, in skin and kidney derived fibroblasts (Fig. 5h-j). These results suggest that Kat5 is a pan-tissue regulator of fibrotic responses to TGF-β stimulation, however certain organs (lung, heart) show a strongest dependence on Kat5 to mount fibrotic responses to TGF-β.

**Figure 5.**
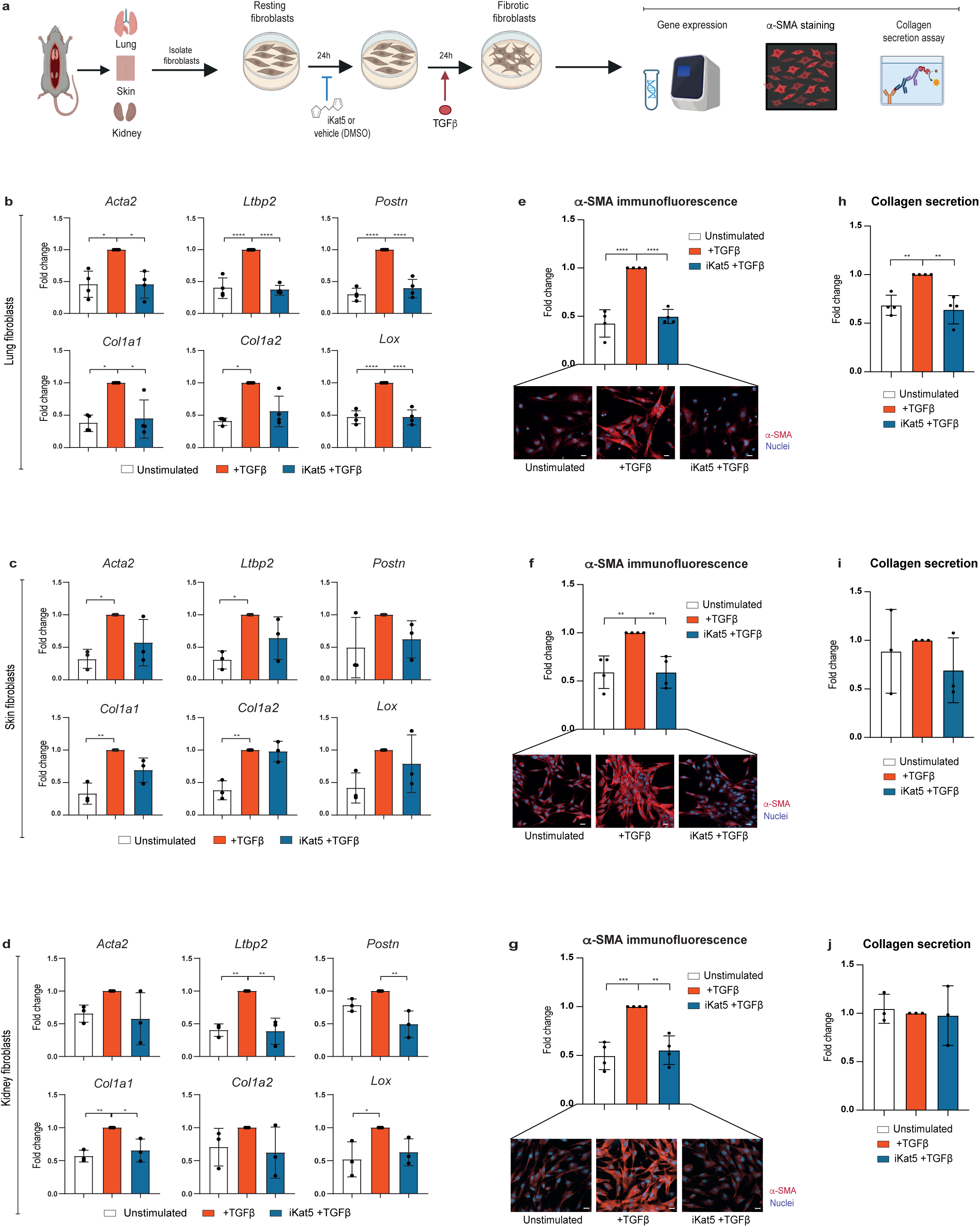
Kat5 inhibition attenuates fibrotic responses in murine primary fibroblasts. **(a)** Schematic depiction of the experimental approach in murine lung, skin and kidney fibroblasts. **(b-d)** Expression of fibrotic marker genes following TGF-β stimulation in control and NU-9056 pre-treated murine lung **(b)**, skin **(c)** and kidney **(d)** fibroblasts. The measurements were performed in three-four replicate experiments. **(e-g)** Quantification of the average immunofluorescence intensity of α-SMA, following TGF-β stimulation in control and NU-9056 pre-treated murine lung **(e)**, skin **(f)** and kidney **(g)** fibroblasts. The measurements were performed in four replicate experiments. Representative images of α-SMA immunostaining assays are shown at the bottom. Scale bars: 20 μm. **(h-j)** Quantification of collagen secretion following TGF-β stimulation in control and NU-9056 pre-treated murine lung **(h)**, skin **(i)** and kidney **(j)** fibroblasts. The measurements were performed in three replicate experiments. All data are shown as fold change *vs.* TGF-β values and are mean ± SD. Statistical significance was analyzed by one-way ANOVA or Kruskal-Wallis tests and is indicated as follows: *P <0.05, **P<0.01, ***P<0.001, ****P <0.0001.

### Inhibition of KAT5 attenuates fibrosis in patient-derived human cardiac fibroblasts

Finally, we decided to investigate the relevance of our findings to human cardiac fibrosis. To this end, we cross-referenced our *ex vivo* Perturb-seq results with the single-cell expression patterns of human cardiac fibroblasts isolated from hearts in different clinical conditions^6,8,9,12,14^ (Fig. 6a). This analysis showed a clear correlation between our *ex vivo* murine fibroblast states and clinically relevant human cardiac subpopulations. For instance, our *ex vivo* naïve fibroblasts were related to human fibroblast subpopulations characteristic of healthy myocardium and our *ex vivo* myofibroblasts were related to fibrotic subpopulations of fibroblasts that appear in failure hearts from patients with myocardial infarction (MI) or Dilated (DCM), Hypertrophic (HCM) or Arrhythmogenic (ARVC) cardiomyopathies (Fig. 6b). In addition, depletion of *Kat5*, *Smad3*, *Srcap* or *Wdr82* in our *ex vivo* murine fibroblasts downregulated gene expression signatures specific of human fibrotic fibroblasts found in pathological conditions (Extended Data Fig. 10a). These results suggest a functional conservation (human-mouse) of the transcriptional regulators that orchestrate cardiac fibrotic responses.

**Figure 6.**
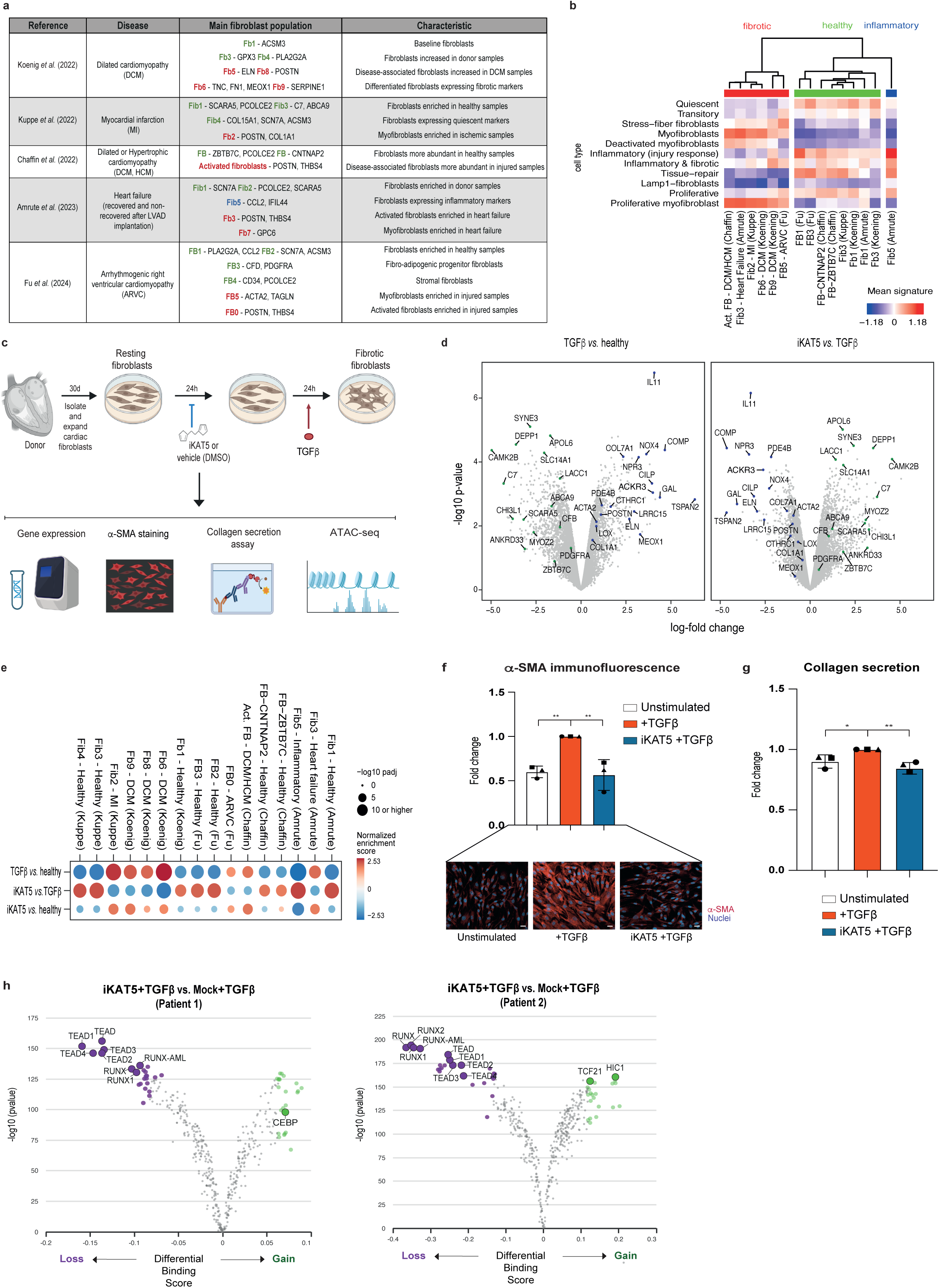
KAT5 inhibition attenuates fibrotic responses in human primary cardiac fibroblasts. **(a)** Table summarizing key papers used for human signature comparisons conducted with the *ex vivo* murine data. **(b)** Enrichment of *in vivo* fibroblast expression signatures from references in (a) over *ex vivo* fibroblast populations from this study. **(c)** Schematic depiction of the experimental approach in human cardiac fibroblasts. **(d)** Differentially expressed markers of fibrosis in TGF-β-treated patient-derived cardiac fibroblasts (left) and in fibroblasts pre-treated with KAT5 inhibitor before the TGF-β stimulus (right). **(e)** Gene set enrichment analysis of differentially expressed genes across treatments. Fibroblast expression signatures are taken from references in (a). The color of each dot represents normalized enrichment score, the size represents the –log10 adjusted p-value. **(f)** Quantification of the average immunofluorescence intensity of α-SMA, following TGF-β stimulation in control and NU-9056 pre-treated human cardiac fibroblasts. The measurements were performed in three patients. Representative images of α-SMA immunostaining assays are shown at the bottom. Scale bars: 20 μm. **(g)** Quantification of collagen secretion following TGF-β stimulation in control and NU-9056 pre-treated human cardiac fibroblasts. The measurements were performed in three patients. Each patient is represented by a different symbol. All data are shown as fold change *vs.* TGF-β values and are mean ± SD. Statistical significance was analyzed by one-way ANOVA test and is indicated as follows: *P <0.05, **P<0.01. **(h)** Differential TF-footprinting analysis between control and NU-9056 pre-treated human fibroblasts after TGF-β stimulation. Dots in purple represent TF motifs with decreased accessibility in NU-9056 pre-treated human fibroblasts. Patient 1 (left) and Patient 2 (right) are shown.

Encouraged by these results we tested the clinical potential of KAT5 inhibition to treat human cardiac fibrosis. To this end, we isolated primary human cardiac fibroblasts from discarded tissue of 2 patients that underwent cardiac surgery, pre-treated them with either KAT5 inhibitor or vehicle, and measured their fibrotic responses after TGF-β stimulation using both gene expression (RNA-seq) and functional analysis (Fig. 6c, Extended Data Fig. 10b). TGF-β treatment induced a marked fibrotic response in unperturbed (vehicle) patient-derived cardiac fibroblasts characterized by the upregulation of typical hallmark fibrosis markers including POSTN, CTHRC1, ACTA2 (α-SMA) (Fig. 6d). Similarly to the murine model, pre-treatment of patient derived-fibroblasts with KAT5 inhibitor markedly reduced the fibrotic response to TGF-β, pushing cardiac fibroblasts back to a healthy (unstimulated) state (Extended Data Fig. 10c-d). Furthermore, patient-derived fibroblasts pre-treated with KAT5 inhibitor downregulated the transcriptional signatures characteristic of pathological human fibroblast subpopulations described above (Fib2-Kuppe, Fib6,8-9-Koenig, Act. FB-Chaffin, Fib3-Amrute, FB0-Fu) (Fig. 6e, Extended Data Fig. 10e). Confirming these results, genetic perturbation (CRISPR) of *KAT5* in patient-derived fibroblasts replicated the pattern observed in using KAT5 chemical perturbation (Extended Data Fig. 10f-g). Analysis of α-SMA protein after KAT5 inhibition showed a clear reduction of this hallmark fibrotic marker, which was reduced to the levels present in unstimulated human fibroblasts (Fig. 6f). Collagen secretion in patient-derived fibroblasts treated with KAT5 inhibitor showed a similar trend but the effect size was much smaller than in murine cardiac fibroblasts (Fig. 6g). This could be due to the pre-activated state of patient-derived fibroblasts, which already present high collagen secretion and, thus, are less prone to modulation by KAT5 inhibition. Finally, chromatin accessibility analysis of patient-derived cardiac fibroblasts treated with KAT5 inhibitor recapitulated the results obtained with murine fibroblasts highlighting the same pattern observed in mice with a clear dependency of TEAD TFs on KAT5 activity, and an antagonistic relationship between KAT5 and CEBP TFs (Fig. 6h). The striking similarity between human and mouse ChrF-TF relationships suggests that the epigenetic mechanisms regulating fibrosis are conserved in evolution. Taken together, these results highlight KAT5 inhibition as a potential therapeutic avenue to alleviate cardiac fibrosis.

## Discussion

In this study, we have highlighted the functional diversity of chromatin complexes as regulators of fibroblast states. We show how specific complexes behave selectively as either mediators or repressors of fibroblast state transitions in response to inflammatory (IL1-β) or fibrotic (TGF-β) stimuli. Several of our top pro-fibrotic regulators belong to the related SRCAP and Tip60 chromatin complexes that, respectively, mediate H2AZ incorporation and acetylation^55^. The connection between H2AZac and TGF-β signaling seems to be a conserved theme as it mediates two other related processes: TGF-β-induced epithelial-mesenchymal transition^56^ and endoderm differentiation, triggered by BMP4/Activin a TGF-β-related signaling pathway^57^. Notably, a recent study has reported a massive rewiring of Tip60 genetic interactions in cells subjected to different extracellular conditions^58^. These findings highlight H2AZ as a central axis modulating cellular responses to environmental changes.

Using chromatin profiling, we provide a mechanistic explanation for the profibrotic roles of Srcap, Kat5, Kat8 and Wdr82 ChrFs. We show that well-known pro-fibrotic TFs such as Egr2 or Teads rely on these chromatin complexes to access their binding sites and induce fibrotic transcriptional responses. Since none of these ChrFs catalyzes nucleosome eviction, which is required for TF accessibility^59^, the mechanisms by which these chromatin complexes facilitate TF accessibility must be indirect. A potential explanation may lie in the low affinity and residency time reported for nucleosomes containing H2AZac histones^60^. This suggests that the deposition of H2AZ by SRCAP and the subsequent acetylation by Tip60 generate permissive chromatin at TGF-β-responsive loci that facilitates the activity of pro-fibrotic TFs upon TGF-β stimulation. In this indirect model, SRCAP and Tip60 are already deployed to profibrotic loci in resting conditions and upon TGF-β stimulation its H2AZ catalytic activities facilitate the subsequent binding of pro-fibrotic TFs. However, our results are also compatible with a direct recruitment of SRCAP and Tip60 by the same TFs they regulate, this has been reported for BAF remodelers^61^. In this scenario, upon TGF-β stimulation SRCAP and Tip60 are deployed to their target loci via interactions with their partner TFs, where they increase the affinity and activity of these TFs by changes in the local chromatin context. Sensitive interactome methods such as TurboID^62^ and single-molecule tracking approaches will be required to elucidate the precise mechanisms underlying these TF-ChrFs dependencies^63^. In any case, our results highlight TF-ChrF partnerships as a key regulatory layer determining cellular states in fibroblasts and other cell types across organs and pathological conditions.

In the second part of our study, we have used patient samples to prove that KAT5 is required for cardiac fibrotic transformation also in humans. Furthermore, these experiments also demonstrate a striking conservation at the molecular level highlighted by the dependency of TEAD TFs on KAT5 in both human and mouse fibrotic responses. While these results are promising, *in vivo* interrogation of Kat5 roles in fibrosis is required to confirm the therapeutic potential of Kat5 inhibitors. In fact, *in vivo* treatment with TH1834, a different Kat5 inhibitor, has been shown to improve cardiac function, cardiomyocyte survival and tissue scarring in a murine model of MI^64^. This study did not examine cardiac fibroblast dynamics, however our results suggest that the improved heart function reported for mice treated with the Kat5 inhibitor is due, at least in part, to an attenuated fibrotic response by cardiac fibroblasts. Indeed, we have also validated *in vitro* that TH1834 reduces cardiac fibroblast activation, providing further evidence of the role of Kat5 in fibrosis. Moreover, our results suggest that Kat5 is a universal mediator of fibrotic transformation in fibroblasts from different organs, therefore a comprehensive interrogation of Kat5 roles in different models of fibrosis could broaden the applicability of Kat5 inhibition to other fibrotic diseases.

Finally, our work highlights how multi-level functional genomics approaches are powerful tools for therapeutic target discovery. We believe that a similar but refined system built from human cardiac organoids^65^ that incorporates a wider variety of fibrotic stimuli^66–69^ will boost the predictive potential of this approach. In addition, *in vivo* functional genomic screens in murine models have emerged as revolutionary tools as they enable to interrogate potential targets in the physiological context in which cells function^70,71^. Recently, efficient *in vivo* CRISPR-mediated editing of cardiac cells has been accomplished using AAV-delivery highlighting the feasibility of *in vivo* Perturb-seq studies in the heart^72–74^. Thus, given the conservation between human and mouse cardiac fibrotic regulators demonstrated here, we advocate for *in vivo* functional studies of disease models in which heart fibrosis is the major pathological outcome. Such studies will aid in developing effective and selective therapies for this widespread clinical condition.

## Methods

### Animal models

All animal requisitions, housing, treatments, and procedures were performed according to all state and institutional laws, guidelines and regulations. All studies were fulfilled under the Guidelines of the Care and Use of Laboratory Animals and approved by the Ethics Committee for Animal Research at the University of Navarra and the Government of Navarra. B6J.129(Cg)-*Gt(ROSA)26Sor^tm1.1(CAG-cas9*,-EGFP)Fezh^*/J (stock Jackson #026179) mouse was purchased from the Jackson Laboratory (USA) and has been previously described^75^.

### Isolation and culture of cardiac fibroblasts

#### Mouse Samples

Primary murine cardiac, lung, skin and kidney fibroblasts were obtained from individual 8-week-old mice (equal ratio of males and females), as previously described^10^. Briefly, after euthanasia, the thorax was opened, the heart was perfused with ice-cold PBS pH 7.6 (Lonza) and was excised together with both lungs, kidneys and a piece of skin from the mouse back (previously shaved) and placed in ice-cold culture media (see below). First, the heart, lungs and kidneys were minced using a sterile scalpel while the skin was cut into small pieces with scissors. Then, pieces of heart, lung and kidney tissues were incubated on an orbital shaker for 7 min at 37°C in the presence of Liberase TH (125 μg/ml) (Roche) in HBSS (Gibco), and skin pieces of tissue were incubated for 45 min at 37°C. After enzymatic incubation, partially digested tissues were mechanically dissociated by slowly pipetting to generate a single-cell suspension. The digestion was repeated with the sedimented pieces, and the supernatants were pooled together. The total time for enzymatic digestion was 20 min for the heart, lung and kidney and 90 min for the skin. Next, the total collected supernatants were filtered through a cell strainer to discard cardiomyocytes (heart) or cell clumps (40 μm, nylon; Falcon) and erythrocytes were removed using RBC lysis buffer (eBioscience). Finally, heart and kidney cell pellets were resuspended in 200 μl of MACS buffer (PBS pH 7.6, 2 mM EDTA, and 0.5% BSA) and incubated with 20 μl of Feeder Removal Microbeads (mEF-SK4) (Miltenyi) for 20 min at 4°C. A positive selection of cardiac and kidney fibroblasts was then performed using LS columns (Miltenyi) according to manufacturer’s instructions. The positive selected fraction (i.e. mEFSK4^+^ cardiac and kidney fibroblasts bound to the column) was resuspended either in serum-free (DMEM KO + 20% knockout serum replacement (KSR) (Gibco) + 1% non-essential amino acids (NEAA) (Gibco) + 100 μM beta-mercaptoethanol (Gibco) + 1% P/S + 1% L-Glutamine) or serum-containing culture media (DMEM + 10% fetal bovine serum (FBS) + 1% P/S + 1% L-Glutamine) containing 10 ng/ml of basic fibroblast growth factor (bFGF) (Peprotech) and plated on 0.1% gelatin-coated (Sigma) wells. One digested heart, two digested lungs, four digested kidneys and the digested piece of skin were seeded into 10 cm^2^ wells. Lung and skin cell pellets were directly plated without Feeder Removal MicroBeads incubation. During the culture process, the growth media was freshly replaced every 2 days and cells were kept in culture for 4 days before being split and expanded. On day 8 of culture, cells were either treated or not with recombinant TGF-β (2.5 ng/ml) (Peprotech) and recombinant IL-1β (20 ng/ml) (ImmunoTools) for 24h.

#### Human Samples

Primary human cardiac fibroblasts from discarded surgical tissue from patients undergoing cardiac surgery were obtained from the University of Navarra Clinic (Spain). Samples were anonymized and the researcher had no access to any personal data. The samples were not obtained for diagnosis purposes and their use was approved by the Ethics Committee of Universidad de Navarra. Tissue enzymatic digestion was performed with forceps to remove fatty pieces and blood and subsequently after transferring to a clean sterile plate, tissue was minced into small pieces using a single-edge blade. Then, 1 ml of collagenase Type II digestion cocktail (0.2 U/ml) (Roche) and trypsin (5×10^-3^ mg/ml) (Gibco) were added and continued mincing until pieces were small enough to transfer with a 1 ml micropipette into a 50 ml conical tube. The conical tube was incubated at 37°C for 20 min with rocking or agitation, was neutralized with 1 ml FBS (Sigma) and centrifuged at 600 g and 4°C for 10 min. Next, the supernatant was removed and the pellet resuspended in 10 ml of DMEM (Gibco) supplemented with 10% FBS and filtered through a 40 μm cell strainer. Tissue pieces were transferred into a new plate and waited for the explant outgrowth. When enough fibroblast outgrowth was observed, the plate was trypsinized and cells were cultured and maintained in fibroblast complete medium (DMEM + 10% FBS + 2.5 mM L-Glutamine (Gibco) + 1% P/S (Gibco) + 0.1% Fungizone Amphoterin B + 1.5% HEPES (Lonza) + 10 ng/ml bFGF (Peprotech)), up to passage five changing the medium every 3 days until experiment began. TGF-β (2.5 ng/ml) (Peprotech) treatment was performed for 24h.

### Flow cytometry and fluorescence-activated cell sorting

Cells were harvested by trypsinization, centrifugated and resuspended into 100 μl of sorting buffer (PBS pH 7.6, 5 mM EDTA, 50 mM Hepes, and 3% FBS) before staining with the respective antibodies against specific surface markers (Supplementary Table 2) for 30 min at 4°C in the dark, following antibody datasheet and manufacturer instructions. Non-stained cardiac fibroblasts were used as negative control and to set the gating strategy. Immunolabelled cells were washed twice with 2 ml of sorting buffer and resuspended in 300 μl of sorting buffer to be analyzed by flow cytometry. For α-SMA intracellular flow cytometry analyses, Cytofix/Cytoperm Fixation/Permeabilization kit (BD Bioscences) was used, following manufacturer instructions.

Flow cytometry was performed on a FACSCANTO (BD Bioscience) Flow Cytometer, whereas fluorescence-activated cell sorting (FACS) was performed using a FACSAria Flow Cytometer (BD Biosciences) or a MoFLoAstrios Flow Cytometer (Beckman Coulter). Standard, strict forward scatter width versus area criteria were used to discriminate doublets and gate only singleton cells. Viable cells were identified by staining with a viability marker (7-AAD, SYTOX Blue or Zombie NIR). The files generated were processed using FlowJo Software (Tree Star, Ashland, USA).

### RNA-seq

#### Mouse Samples

Low input 3’-end bulk RNA-seq was performed using molecular crowding single-cell RNA barcoding and sequencing (mcSCRB-seq^76^) protocol adapted for bulk RNA-seq. Briefly, 3,000 to 10,000 fibroblasts were homogenized in 100 μl of Lysis/Binding Buffer (Ambion) vortexed and stored at -80°C until further processing. Poly-A RNA was extracted with Dynabeads Oligo dT (Ambion) and eluted in 8 μL Tris Cl pH 7.5. Before denaturing at 72°C for 3 min, 1 μL of 2 μM barcoded oligo-dT primer was added. Reverse transcription was performed at 42 °C for 90 min in a mixture containing: 1X Maxima RT buffer, 7.5 % PEG 8000, 1 mM dNTPs, 5 μM of unblocked template-switching oligo, and 240 U of Maxima H, Minus Reverse Transcriptase (Thermo Scientific). Primer excess was removed through digestion with Exonuclease I (NEB). cDNA was purified with a 1.2X SPRI clean up (Agencourt AMPure XP, Beckman Coulter), then subjected to PCR amplification using Terra polymerase (Takara Bio) and SINGV6 primer with the following thermal cycling parameters: 3 min at 98°C followed by 6 cycles of 15 s at 98°C, 30 s at 65°C, and 4 min at 68°C, and a final elongation step of 10 min at 72°C. Upon purification (1X SPRI clean-up), 0.8 ng of double-stranded cDNA were tagmented using Nextera-XT Tn5 (Illumina). PCR amplification using P5NEXTPT5 primer and i7 indexed primers allowed 3’ enrichment and secondary barcoding of the libraries as well as addition of Illumina adaptor sequences. Libraries were quantified using a Qubit 3.0 Fluorometer (Invitrogen) and their size profiles examined in Agilent’s 4200 TapeStation System (Agilent). Sequencing was carried out in an MGI DNBSEQ-G400 instrument at a depth of 10 million reads per sample.

#### Human Samples

Briefly, 75,000 to 100,000 human fibroblasts were homogenized in 100 μl of Lysis/Binding Buffer (Ambion) vortexed and stored at -80°C until further processing. Poly-A RNA was extracted with Dynabeads Oligo dT (Ambion) and eluted in 8 μL Tris Cl pH 7.5. RNA was reverse transcribed with PrimeScript Reverse Transcription reagent kit (Takara) according to manufacturers’ protocols. Double-stranded cDNA was obtained by second-strand synthesis (SSS) using the SSS kit (Invitrogen) starting with 3 ng of cDNA following the manufacturer’s protocol. Purified cDNA was then converted to sequencing library using the Illumina Nextera XT DNA library prep with 1 ng of cDNA and 12 cycles of PCR amplification. The quality and quantity of the libraries were verified using Qubit dsDNA HS Assay Kit (Thermo Fisher Scientific) and 4200 Tapestation with High Sensitivity D1000 ScreenTape (Agilent Technologies). Libraries were then sequenced using a NextSeq2000 sequencer (Illumina). 20-30 million pair-end reads (100 bp; Rd1:51; Rd2:51) were sequenced for each sample and demultiplexed using bcl2fastq.

#### Bioinformatic analyses

RNA-seq data analysis was performed using the following workflow: (1) the quality of the samples was verified using FastQC software (https://www.bioinformatics.babraham.ac.uk/projects/fastqc/) and the trimming of the reads with trimmomatic^77^; (2) alignment against the mouse (mm10) or human (GRCh38) reference genome was performed using STAR (v.2.7.3)^78^; (3) gene expression quantification using read counts of exonic gene regions was carried out with *featureCounts*^79^; and (4) the gene annotation reference was Gencode v45^80^. Data filtering and normalization, as well as differential gene expression analysis, was carried out with the edgeR^81^ (v4.0.14) and limma^82^ (v3.58.1) packages. Gene set enrichment analysis (Fig. 6e, Extended Data Fig. 10e) was performed with *fgsea* (v1.28.0)^83^, which estimates p-values using an adaptive multi-level split Monte-Carlo scheme and calculated corrected p-values using the Benjamini–Hochberg method with the R function *p.adjust*. Gene expression profiles (Extended Data Fig. 10d) were compared by (1) removing batch effects in normalized counts via the function *removeBatchEffect* in limma and (2) calculating Pearson correlation coefficients rij (R function cor) and using these coefficients as distances 1 − rij for hierarchical clustering (R function *hclust(method = “complete”)*).

### RNA extraction, reverse transcription and quantitative real-time PCR

Poly-A RNA from cells was extracted using Dynabeads Oligo dT (Ambion) and reverse-transcribed with PrimeScript Reverse Transcription reagent kit (Takara) according to manufacturers’ protocols. Quantitative real-time PCR (qPCR) was carried out using PowerUp SYBR Green Master Mix (Applied Biosystems) for mouse samples or using TaqMan Fast Advanced Master Mix (Applied Biosystems) for human samples. All primers used for qPCR are listed in Supplementary Tables 3-4. QuantStudio 5 Real Time PCR Systems (ThermoFisher) or AriaMx Real-Time PCR System (Agilent Technologies) were used to run qPCR. Once the reaction was finished, data were exported and analyzed using QuantStudio or AriaMx softwares. All quantifications were normalized to endogenous control genes (*Gapdh*, *Rpl4*) and were analyzed in terms of relative quantification expressed as fold change difference using the 2^-ΔΔCt^ method.

### FACS-based CRISPR screens

#### CRISPR library construction

sgRNA-CRISPR library (Supplementary Table 1) was ordered from Integrated DNA Technologies and cloned using a Gibson assembly master mix (New England Biolabs) in the CRISPR sequencing (CRISP-seq-BFP) backbone (stock Addgene #85707). The Gibson assembly product was electroporated in Endura Electrocompetent Cells (Lucigen #60242) and plated on Bioassay LB agar plates. After 20h at 30°C bacteria were harvested, and plasmid preparations were performed using NucleoBond Xtra Midi kit (Machery-Nagel) following the manufacturer’s protocol.

#### Cloning of individual sgRNAs

Individual oligonucleotides were cloned in the CRISP-seq-BFP backbone using a Golden-Gate reaction with 100 ng of vector backbone, 1 μl of annealed sgRNA oligonucleotides, 1 μl Esp3I (New England Biolabs) and 1 μl T4 DNA Ligase (New England Biolabs) using the following program: 10x (5 min at 37°C, 10 min at 22°C), 30 min at 37°C and 15 min at 75°C. The Golden-Gate product was heat-shock transformed in One Shot Stbl3 *E. coli* Cells (Invitrogen #C737303) and plated on LB agar plates for 20h at 30°C. Individual colonies were picked and grown overnight. Plasmids were isolated with the NucleoSpin Plasmid MiniPrep Kit (Macherey-Nagel) and sequenced using a U6 forward primer (Supplementary Table 5).

#### Lentiviral production

HEK293T cells were transfected with the CRISP-seq-BFP vectors, pMD2-G (stock Addgene #12259) and psPAX2 (stock Addgene #12260) using Lipofectamine 3000 (Invitrogen) according to the manufacturer’s protocol. After 8h, media was replaced with Opti-MEM (Thermo Fisher Scientific). The viral supernatant was collected 48h after transfection, filtered using 0.45-μM filters (Millipore) and concentrated via centrifugation at 3,000g at 4°C using a 100 kDa Amicon Ultra-15 Centrifugal Filter (Millipore).

#### Cell culture and experimental design

Murine cardiac fibroblasts were isolated from Rosa26-Cas9 mice as previously described. Two days after plating in serum-free medium, cells were transduced with the CRISPR library to reach a MOI of approximately 30%. After 24h, fresh medium was added, and cells were maintained in culture and expanded. Then, six days after infection, cells were treated with recombinant TGF-β (2.5 ng/ml) for 24h. Finally, cells were collected and stained with PDGFRα-PE antibody (Supplementary Table 2) plus a viability marker (7-AAD) for separately sorting naïve (15% PDGFRα-high) and fibrotic (15% PDGFRα-low) fibroblasts from the viable transduced (BFP^+^) Cas9 (GFP^+^) population. An exemplar gating strategy can be found in Supplementary Fig. 1. The experiment was performed in parallel using non-Cas9 expressing cardiac fibroblasts extracted from C57BL/6J mice.

#### FACS-based CRISPR library preparation, sequencing and preprocessing

Sorted cells were lysed in 40 μl of 0.2% SDS and 2 μl of proteinase K (New England Biolabs) at 42°C for 30 min. Then, gDNA was isolated with a 2x solid-phase reversible immobilization (SPRI) cleanup and NGS libraries were prepared from purified gDNA with a two-step PCR protocol using 2x KAPA HiFi Master Mix (Roche). First PCR: 10 μM Read1-U6 and Read2 scaffold primer mix (Supplementary Table 5); 3 min at 98°C, 20x (10 s at 98°C, 10 s at 62°C, 25 s at 72°C), 2 min at 72°C. Second PCR: 10 μM P5 and P7 index primer mix (Supplementary Table 5); 3 min at 98°C, 10x (10 s at 98°C, 10 s at 62°C, 25 s at 72°C), 2 min at 72°C. Libraries were purified with 1.6x SPRI cleanup and sequenced at a depth of 5 million reads per sample in a NextSeq 2000 system. Raw data were processed with bcl2fastq (v.2.20) into FASTQ files and then processed using a custom script to isolate the 20-mer protospacers; then, they were mapped using Bowtie2 (v.2.3.4.2) using an index file containing the sgRNA sequences (Supplementary Table 1).

#### Computational analysis of FACS-based CRISPR screens

For the bulk screen analysis detailed in Fig. 1, Extended Data Fig. 2a, sgRNA counts from bulk phenotypic screens were normalized against mean counts from NTC guides using the geometric mean, and the sgRNA-level log2-fold enrichment/depletion was calculated using normalized sgRNA counts comparing two paired screens of interest (i.e., Cas9 *vs*. wildtype cells, or Fibrotic *vs*. Naïve). The replicates were also used to calculate a p-value via a t-test. To calculate gene-level scores from sgRNA-level log2-fold changes and p-values, we took the median over sgRNAs that target the same genes and p-values were combined using the Fisher method. The Benjamini-Hochberg false discovery rate (FDR) was applied to calculate corrected p-values for the 726 genes tested.

#### Validation of single candidates with flow cytometry

For individual validations of candidate factors, isolated cardiac fibroblasts from Rosa26-Cas9 mice were transduced at day 2 of culture with the LV-CRISP-seq-BFP vector containing either a sgRNA against the candidate factor (*Tgfbr1, Smad4, Srcap*, *Kat5*, *Kat8*, *Hcfc1*, *Wd82, Ino80, Smarca4, Brd9, Paxip1*) or a NTC guide. Then, cells were maintained in culture until day 8 when they were stimulated, or not, with recombinant TGF-β (2.5 ng/ml) for 24h. After that, cells were collected and stained with PDGFRα-PE antibody (Supplementary Table 2) for flow cytometry analysis.

#### Validation of single candidates with Indel-seq

For assessing CRISPR/Cas9 gene editing in target genes, isolated cardiac fibroblasts from Rosa26-Cas9 or non-Cas9 (wildtype) mice were transduced at day 2 of culture with the LV-CRISP-seq-BFP vector containing a sgRNA against the candidate factor (*Tgfbr1*, *Smad4*, *Srcap*, *Kat5*, *Kat8*, *Hcfc1*, *Wd82*). Cells were maintained in culture for one week and then, viable transduced (BFP^+^) Cas9 (GFP^+^) cells, were FACS-sorted and processed for Indel-seq analysis. For that, cells were lysed in 40 μl of 0.2% SDS and 2 μl of proteinase K (New England Biolabs) at 42°C for 30 min. Then, gDNA was isolated with a 2x solid-phase reversible immobilization (SPRI) cleanup and NGS libraries were prepared from purified gDNA with a two-step PCR protocol using 2x KAPA HiFi Master Mix (Roche). First PCR: 10 μM Read1-gDNA_*target* and Read2-gDNA_*target* primer mix (Supplementary Table 5); 3 min at 98°C, 22x (10 s at 98°C, 10 s at Tm primers, 15 s at 72°C), 2 min at 72°C. Second PCR: 10 μM P5 and P7 index primer mix (Supplementary Table 5); 3 min at 98°C, 8x (10 s at 98°C, 10 s at 62°C, 25 s at 72°C), 2 min at 72°C. Libraries were purified with 1.6x SPRI cleanup and sequenced at a depth of 1 million reads per sample in a NextSeq 2000 system. Raw data were processed with bcl2fastq (v.2.20) into FASTQ files and then mapped to the reference genome using Bowtie2 (v.2.3.4.2). Final files were visualized using IGV (v2.18.2) software.

#### Validation of single candidates with Western blot

For assessing CRISPR/Cas9 consequences at target protein levels, isolated cardiac fibroblasts from Rosa26-Cas9 mice were transduced at day 2 of culture with the LV-CRISP-seq-BFP vector containing either a sgRNA against the candidate factor (*Tgfbr1*, *Smad4*, *Kat5*) or a NTC guide. The transduction of fibroblasts was achieved at a high MOI to ensure at least 70-90% of transduced cells within the culture. Then, cells were maintained in culture and collected one week after transduction for Western blot analysis. For that, total protein was extracted using a commercial Pierce RIPA buffer (Thermo Scientific) supplemented with 1x cOmplete Mini, EDTA-free (Roche), and quantified using Bradford Protein Assay (BIO-RAD). Proteins were separated by electrophoresis on 10% acrylamide gel and blotted onto nitrocellulose membranes (Bio-Rad, 0.45 μm). Total protein quantification for normalizing target signals was calculated using Revert700 total protein stain (LI-COR) and an Odyssey imaging system (Osyssey FC, LI-COR). Membranes were blocked in the blocking solution (5% non-fat dry milk powder in TBS-0.05% Tween-20) for 1h at room temperature and then the corresponding antibody was diluted according to the instructions (Supplementary Table 6) and incubated overnight at 4°C. Membranes were subsequently washed using TBS-0.05% Tween-20 solution and incubated with the HRP-conjugated secondary antibody (Supplemental Table 6) for 1h at room temperature. Finally, immunoblots were visualized by an enhanced chemiluminescence detection kit (SuperSignal West Femto, Thermo Scientific) under a chemiluminescence imaging analysis system (ChemiDoc MP, Bio-Rad). Uncropped versions of blots (for both total and target proteins) can be found in Supplementary Fig. 2.

### Perturb-seq

#### Perturb-seq library construction

For each target, we cloned the top two performing sgRNAs in the LV-Perturb-seq-BFP vector, which was built modifying the original LV-CRISP-seq-BFP vector by replacing the original sgRNA scaffold for a sgRNA scaffold containing the 10x capture-sequenced CR1Cs1^84^, as previously described^85^. Lentiviral particles were prepared as specified for FACS-based CRISPR screens.

#### Cell culture and experimental design

Murine cardiac fibroblasts were isolated from Rosa26-Cas9 mice and transduced at day 2 of culture with the Perturb-seq library to reach a MOI of approximately 10%. After 24h, fresh medium was added, and cells were maintained in culture and expanded. Then, six days after infection, cells were treated for 24h under three different stimulation conditions: TGF-β (2.5 ng/ml), IL-1β (20 ng/ml) or without stimulation. Finally, cells were collected and viable (7-AAD^-^) transduced (BFP^+^) Cas9 (GFP^+^) fibroblasts were FACS-sorted and processed in the Chromium Controller aiming to reach 200 single cells per sgRNA. An exemplar gating strategy can be found in Supplementary Fig. 3.

#### Perturb-seq library preparation and sequencing

Single-cell libraries were generated using the Chromium Next GEM Single Cell 3′ Reagent Kits v.3.1 (Dual Index) following the manufacturer’s recommended protocol. The resulting libraries were sequenced in a NextSeq 2000 system to a final coverage of ∼200,000 reads per cell for 3′ Gene Expression libraries and 10,000 reads per cell for CRISPR Feature Barcode libraries.

#### Computational analysis of Perturb-seq

Analysis was primarily performed in Python (v.3.10.12) using the *scanpy* package (v1.10.3) leveraging much of the inbuilt functionality. Cells were log-normalized with a 10^4 scaling factor. For visualizations, we used *scanpy* in-built functions as well as R (v4.3.2), in particular *ggplot2* (v3.4.4) and *ComplexHeatmap* (v2.18.0) packages. Details are outlined below.

Clustering (Fig. 2b onwards) was performing using the Leiden algorithm with standard *scanpy* presets was employed; and cell labels were chosen via maximum overlap with predefined lists of markers, calculated using the Jaccard index. Differentially expressed genes (DEGs) were calculated using the inbuilt *scanpy* functionality.

Gene signature scores (Extended Data Fig. 2d-e) were calculated from log-normalized counts by calculating the mean expression over all genes associated with a signature. Fibroblast subtype signatures were obtained from Supplementary Table 5 from Buechler *et al.*^11^, Supplementary Table 27 from Koenig *et al.*^12^, Supplementary Table 3 from Forte *et al.*^13^, Supplementary Table 10 from Amrute *et al*^6^., Supplementary Table 11 from Chaffin *et al*.^8^, Additional File 6 from Fu *et al*.^14^ and Supplementary Table 13 from Kuppe *et al*.^9^. Human genes were mapped to mouse homologs using a mapping downloaded from Ensembl BioMart on 2021-07-27.

Enrichment analysis of cells (Fig. 2f, Extended Data Fig. 5) for specific epigenetic perturbations across the different fibroblast states were determined by testing whether the abundance of a target in a cluster (relative to NTCs) was greater than the abundance of this target in the remaining cells. Fisher’s exact test was used to obtain odds ratios and p-values. Multiple test correction was performed using the Benjamini–Hochberg method with the R function *p.adjust*.

Log-fold changes of genes were subjected to gene set enrichment analysis (Extended Data Fig. 6b) as implemented in *fgsea* (v1.28.0)^83^, as described for bulk RNA-seq data. Gene sets comprised KEGG_2019_Mouse, MsigDB_Hallmark_2020, Reactome_2022, and WikiPathways_2019_Mouse, which were downloaded from *Enrichr*^86^. Genes were ranked according to their log-fold change of expression relative to NTCs.

Analysis of gene expression patterns as they relate to TF signatures (Extended Data Fig. 3) was performed using the Python implementation of the SCENIC algorithm^87^ using default settings (pyscenic; v0.12.1).

### Functional assays in fibrosis

#### Cell culture and experimental design

For CRISPR/Cas9 loss-of-function, murine cardiac fibroblasts were isolated from Rosa26-Cas9 mice as previously described and plated in serum-free medium. At day 2 of culture, fibroblasts were transduced with the LV-Perturb-seq-BFP vector containing individual sgRNAs against the candidate factor (*Srcap*, *Kat5*, *Kat8*, *Hcfc1*, *Wd82*) or a NTC guide. A LV-Perturb-seq-BFP vector containing sgRNA against *Tgfbr1* gene was also included as a positive control. The transduction of fibroblasts was achieved at a high MOI to ensure at least 80-90% of transduced cells within the culture. Then, cells were maintained and expanded in culture until day 6 after transduction, when perturbed fibroblasts were collected, counted, and plated for ELISA, α-SMA immunofluorescence (IF) or contraction assay, as detailed below. Otherwise, cell proliferation and apoptosis were evaluated across all the culture process.

For chemical inhibition of *Kat5* activity, murine (cardiac, lung, skin and kidney) and human fibroblasts were extracted as previously described. At day 7 of culture, seeded fibroblasts were collected and plated for ELISA, α-SMA or H2AZac IF and treated with 5 μM of NU-9056 inhibitor (Abcam), whereas control fibroblasts were treated with the same volume of vehicle (DMSO) for 24h. Following this treatment, cells were stimulated with TGF-β (2.5 ng/ml) for 24h.

#### ELISA

The content of Collagen type I in cell supernatants was determined using the Sandwich ELISA kit for Collagen Type I (Antibodies online) for mouse samples and Human Pro-Collagen I alpha 1 ELISA Kit (R&D systems) for human samples, according to the manufacturer’s protocols. Briefly, for mouse samples, 100 μl of each standard dilutions, blank and samples were added to the wells, and the plate was incubated for 1h at 37°C. Then, the liquid was removed from each well and 100 μl of Detection Reagent A Working Solution was added to each well, and incubated at 37°C for 1h. The solution was aspirated and 350 μl of 1x Wash Solution was added into each well and was soaked for 1-2 min. The remaining liquid was removed from all wells and washed 3 times with wash Buffer. 100 μl of the Detection Reagent B Working Solution was added to each well, and incubated for 30 min at 37°C. The aspiration/washing procedure was repeated a total of 5 times. 90 μl of Substrate Solution was added to each well and incubated for 10-20 min at 37°C, protecting it from light. 50 μl of Stop Solution was added to each well and the microplate reader was run and immediately the measurement was taken at 450 nm. For human samples, 50 μL of Assay Diluent RD1W was added to each well, followed by 50 μL of sample. The plate was incubated for 2h at room temperature on a shaker at 500 rpm, and wells were aspirated and washed 4 times with 400 μL of Wash Buffer. Next, 200 μL of Human Pro-Collagen I α1 Conjugate was added to each well, and the plate was incubated for 1h at room temperature on the shaker and wells were washed as before. 200 μL of Substrate Solution was added, and the plate was incubated for 30 min at RT. 50 μL of Stop Solution was added and measured in a microplate reader at 450 nm.

#### α-SMA IF

For IF staining, Zinc formalin-fixed cells were permeabilized with 0.1% triton for 20 min and incubated with 0.4% gelatin for 20 min at room temperature to avoid unspecific antibody binding. Then, cells were incubated with a 1:200 dilution of the anti-α-SMA antibody (Sigma, A5228) overnight at 4°C in the dark. Mounting medium (PBS:Glicerol; 1:1) without Hoechst for perturbed cells and with Hoechst for inhibitor-treated cells was used to visualize the preparation. Images were taken in a cell observer Axio Imager M1Esc microscope (ZEISS) for genetically perturbed fibroblasts and in a LSM 800 confocal microscope (ZEISS) for inhibitor treated fibroblasts. α-SMA fibers formation was indirectly assessed as a measure of the staining intensity within delimited single cells, subtracting the background fluorescence intensity (i.e. areas with no cells): a.u. = Icytoplasm – Ibackground. Only transduced (BFP^+^) cells were considered in the analysis for the perturbed fibroblasts. Image analysis was performed using the ImageJ software.

#### H2AZac IF

For H2AZac IF staining, Zinc formalin-fixed cells were permeabilized with 0.1% triton for 20 min and incubated with 0.4% gelatin for 20 min at room temperature to avoid unspecific antibody binding. Then, cells were incubated with a 1:400 dilution of the anti-H2AZac antibody (Diagenode, C15410202) overnight at 4°C in the dark. On the next day, cells were incubated with a 1:500 dilution of the fluorescence-conjugated secondary antibody, anti-rabbit 594 (Invitrogen, A11012) 1h at room temperature in the dark. Mounting medium (PBS:Glicerol; 1:1) with Hoechst was used to visualize cell nuclei and images were taken with a 20x objective in a LSM 800 confocal microscope (ZEISS). The levels of H2AZac were quantified by first counting the total number of cells and then determining the number of H2AZac-positive cells. Finally, the percentage of H2AZac cells was calculated based on these values. Image analysis was performed using the ImageJ software.

#### Contraction assay

Cell contraction assay was performed as previously described^4^ with minor modifications. Briefly, 45,000 cells were mixed with collagen-I (Corning) and allowed to solidify for 60 min at 37°C. Then, fibroblasts embedded into the collagen pads were induced to differentiate during 24h with recombinant TGF-β (2.5 ng/ml). Gels were released, and the areas of floating collagen pads were measured and compared to the control conditions (NTC) 24h later.

#### Proliferation

Growth was monitored by flow cytometric assessment of BFP^+^ cells in culture at 2, 4, 5, 6, 7 and 8 days after transduction. Fold change in BFP^+^ cells at each time point was calculated relative to the proportion of BFP^+^ cells at day 2.

#### Apoptosis

Apoptosis was evaluated by flow cytometry using the Annexin V DY634-APC kit (Immunostep), according to the manufacturer’s protocol with minor modifications. Harvested cells were trypsinized, centrifuged, and washed in PBS. Cells were resuspended in 100 μl of 1x Annexin V Binding Buffer (BD Biosciences) at 1-5×10^6^ cells/ml. 2.5 μl of Annexin V-APC was added to 100 μl of the cell suspension and incubated for 10-15 min at RT, protecting it from the light. Finally, 400 μl of 1x Annexin V Binding Buffer plus a viability marker (7-AAD) were added to be analyzed by flow cytometry.

### Chromatin accessibility analysis of chromatin factor knockouts

For CRISPR/Cas9 loss-of-function, murine cardiac fibroblasts were isolated from Rosa26-Cas9 mice as previously described and plated in serum-free medium. At day 2 of culture, fibroblasts were transduced with the LV-Perturb-seq-BFP vector containing individual sgRNAs against the candidate factor (*Srcap*, *Kat5*, *Kat8*, *Wd82*) or a NTC guide. Then, cells were maintained and expanded in culture until day 6 after transduction, when perturbed fibroblasts were either stimulated or not with TGF-β (2.5 ng/ml) for 24h. Afterwards, viable (7-AAD^-^) transduced (BFP^+^) Cas9 (GFP^+^) fibroblasts were FACS-sorted for subsequent ATAC-seq protocol.

For chemical inhibition of *Kat5* activity, murine and human cardiac fibroblasts were extracted as previously described. At day 7 of culture, seeded fibroblasts were treated with 5 μM of NU-9056 inhibitor whereas control fibroblasts were treated with the same volume of vehicle (DMSO) for 24h before TGFβ (2.5 ng/ml) stimulation. After 24h of stimulation, fibroblasts were collected and counted for subsequent ATAC-seq protocol.

ATAC-seq was performed according to the Fast-ATAC protocol previously described^10^. Briefly, 10,000 freshly sorted cells were centrifuged at 500 g for 5 min at 4°C and pellet resuspended into 25 μl of the transposase mix which includes 1x TD buffer (Illumina), 0.05% digitonin (Sigma) and 1 μl of TDE1 (Illumina). Transposition reactions were incubated at 37°C for 30 min with shaking at 450 rpm. Then, reactions were stopped at 4°C for 5 min. To release tagmented DNA, samples were incubated with 5 μl of cleanup buffer (900 mM NaCl (Sigma), 30 mM EDTA (Millipore), 2 μl of 5% SDS (Millipore) and 2 μl of Proteinase K (NEB)) for 30 min at 40°C. The isolated tagmented gDNA was purified with a 2x SPRI Cleanup. Finally, tagmented genomic regions were amplified by PCR using KAPA HiFi DNA Polymerase (Roche) and 5 μM P5 and P7 Nextera Indexing Primers (Supplementary Table 5), using the following program: 5 min at 72°C, 2 min at 98°C, 10x (98°C for 20 s, 60°C for 30 s, 72°C for 1 min) and 5 min at 72°C. The ATAC-seq libraries were sequenced at 30 million reads on a NextSeq 2000 system. Two replicate experiments (murine fibroblasts) and two patients (human fibroblasts) were analyzed.

#### Data processing and analysis

ATAC-seq data was processed using as a guideline the bioinformatic pipeline available in nf-core^88^. Briefly, low-quality bases and adapter sequences were removed with Trim Galore (v0.6.6)^89^, ensuring high-quality read inputs. Reads were aligned to the reference genome (GRCm38/mm10 or GRCh38, including decoy sequences) using Bowtie2 (v2.3.4.2)^90^ with parameters set to -X 1000 --no-discordant --no-mixed --very-sensitive. We then removed duplicates with Picard (v2.25.4) and excluded mitochondrial DNA (chrM), Epstein-Barr virus (chrEBV), and any sequences classified as ’chrUn’, ’random’ or overlapping with ENCODE blacklist regions (v2.0)^91^. Finally, we corrected the Tn5 insertion bias using alignmentSieve (v3.5.1)^92^ with --ATACshift parameter and used bamCoverage (v3.5.1) to get CPM scaled Bigwig files.

To identify accessible regions, we pooled BAM files from biological replicates and converted them to single-read BED format using *bamToBed* (BEDTools v2.27.1)^93^. Then, we ran MACS2 (v2.2.7.1)^94^ with parameters --broad -f BED --keep-dup all --nomodel --shift -75 --extsize 150. The obtained peak coordinates were used to assess differential TF motif (HOMER’s known vertebrate motifs) enrichment between ChrF-KO and NTC conditions in TOBIAS (v0.13.2)^95^. Specifically, we generated a consensus peak set by integrating peaks from both control and KO samples. Then, we generated TF footprint Bigwig files using TOBIAS’s *ATACorrect* function (parameters: --read_shift 0 0) and *ScoreBigwig* with default settings. Finally, we identified differentially bound TF motifs between conditions using *BINDetect* from TOBIAS.

### ChIP-seq

#### Dual cross-linking

Primary cardiac fibroblasts were isolated as previously described and cultured up to passage 2 in serum-free medium. Cells were either treated or not with TGF-β (2.5 ng/ml) for 24h and then cross-linked. For that, supernatant was removed, and cells were washed with PBS. Enough buffer (0.1% MgCl (Invitrogen)-PBS) was added to cover properly the cells. 3 mM of ethylene glycol bis(succinimidyl succinate), disuccinimidyl glutarate and dimethyl adipimidate (Thermo Fisher Scientific) were added next and incubated with agitation for 20 min at RT. 1% formaldehyde (Thermo Fisher Scientific) was added and incubated for another 5 min at RT. Then, glycine was added to 130 mM and incubated for 5 min to quench the cross-linkers. Cells were placed on ice and 1x cOmplete Protease Inhibitors (Roche) and 0.5% BSA were added and incubated 10 min on ice. Supernatant was removed and cells were washed with PBS-0.5% BSA + 0.1x Protease Inhibitors. Finally, PBS-0.5% BSA + 1x Protease Inhibitors was added, cells were scraped, aliquoted in dolphin tubes and centrifuged at 1,000 g for 10 min at 4°C. Supernatant was removed and samples were flash-frozen at −80°C. Two replicates per condition were analyzed.

#### Chromatin immunoprecipitation

Cross-linked cells were thawed and resuspended in 1.5 ml ice-cold cell lysis buffer (10 mM HEPES, pH 7.5, 10 mM NaCl, 0.2% NP-40 (Thermo Fisher Scientific)) plus 1x cOmplete Protease Inhibitors for 10 min on ice. Then, nuclei were pelleted at 5,000g for 7 min, resuspended in sonication buffer (0.5% SDS, 5 mM EDTA) and pelleted again at 8,000g, then resuspended in 50-100 μl sonication buffer and sonicated for five cycles (30s ON, 30s OFF) in a Bioruptor Nano (Diagenode). Then, chromatin extracts were diluted in four volumes of ChIP dilution buffer (25 mM HEPES, 185 mM NaCl, 1.25% Triton X-100 plus 1x cOmplete Protease Inhibitors) and incubated with 2 μg of H2Az-Ac antibody (#C15410202, Diagenode) at 4°C for 10-12h. The following day, 25 μl Magna ChIP Protein A + G (Merck Millipore) were added and incubated for 3h at 4°C. Bead-bound chromatin was washed twice with radioimmunoprecipitation assay (RIPA) buffer (10 mM Tris-Cl, pH 8, 150 mM NaCl, 0.1% SDS, 1% Triton X-100, 1 mM EDTA), twice with RIPA-500 buffer (10 mM Tris-Cl, pH 8, 500 mM NaCl, 0.1% SDS, 1% Triton X-100, 1 mM EDTA), twice with LiCl buffer (10 mM Tris-Cl, pH 8, 550 mM LiCl, 0.5% sodium deoxycholate, 0.5% NP-40, 1 mM EDTA) and once with TE buffer. ChIPped DNA was reverse-cross-linked by 30 min incubation with 2 μl proteinase K in 50 μl ChIP elution buffer (10 mM Tris-Cl, pH 8, 300 mM NaCl, 0.2 mM EDTA, 0.4% SDS) at 55°C followed by 1h incubation at 68°C. Finally, the ChIPped DNA was purified with a 2.2x SPRI cleanup and quantified using the Qubit dsDNA HS Assay Kit (Thermo Fisher Scientific).

#### Preparation of ChIP–seq libraries

ChIP–seq libraries were prepared from 5 ng of ChIPped DNA using the Next Ultra II kit (New England Biolabs) following the manufacturer’s instructions. ChIP–seq libraries were sequenced to 50 million reads per sample (paired-end 50 bp) in a NextSeq 2000 system and demultiplexed using bcl2Fastq (v.2.20).

#### ChIP–seq data processing and analysis

Raw sequencing data, provided in FASTQ format, were processed through a detailed bioinformatics pipeline for quality control, alignment, peak identification, and subsequent analytical steps. We evaluated the sequence quality of the data with FastQC (v0.11.9. Adapters and bases below a Q20 quality threshold were trimmed using Cutadapt (v3.4), ensuring high-quality read inputs for alignment.

Alignment to the reference genome (mm10) was performed with Bowtie2 (v2.4.2), followed by sorting and indexing of alignment outputs via Samtools (v1.10). We then removed duplicates and excluded mitochondrial DNA (chrM), Epstein-Barr virus (chrEBV), and any sequences classified as ’chrUn’, ’random’ or included ENCODE blacklist regions v2.0. Furthermore, we identified enriched ChIP signal regions from peak calling with MACS2 (v2.2.7.1) using the ’--broad’ flag suitable for histone marks and a broad-cutoff of 0.1.

Consensus peak regions were derived by merging overlapping peaks across biological replicates with HOMER’s *mergePeaks* function. Regions were then annotated with annotatePeaks.pl from HOMER. Differential binding analysis utilized *featureCounts* (Subread package v2.0.1) for read count aggregation within consensus peaks, and DESeq2 (v1.30.0) to compare signal enrichment between conditions. Significant peaks were selected based on an absolute log2(fold change) greater than 0.75 and an adjusted P-value (Padj) less than 0.01, ensuring the identification of biologically meaningful differences. Visualization and additional exploratory analyses, such as principal component analysis (PCA), scatter-plot and hierarchical clustering, were enabled by custom R scripts.

#### Motif analysis in ChIP–seq peaks

Motif analysis was conducted on significantly enriched or depleted H2AZac peaks identified in pairwise comparisons between conditions. We employed *findMotifsGenome* function from HOMER (v4.10) by comparing altered peaks to a background set composed of all peaks identified across conditions, excluding the peaks of interest to maintain an unbiased background. The search for known motif enrichment was centered around a 200 bp window surrounding the peak summits to ensure specificity to the regions of highest binding affinity.

### CRISPR/Cas9 in human cardiac fibroblasts

To induce CRISPR loss-of-function in primary human cardiac fibroblasts, cells were simultaneously transduced with the LV-Cas9-GFP vector (stock Addgene #82416) and the LV-CRISP-seq-BFP vector containing a sgRNA against *KAT5* or a NTC guide (Supplementary Table 7). Then, cells were maintained and expanded in culture until day 5 post-infection, when the medium was replaced by fibroblast minimum medium (DMEM + 2.5 mM L-Glutamin (Gibco) + 1% Penicillin/Streptomycin (Gibco) + 0.1% Fungizone Amphoterin B + 1.5% HEPES (Lonza) + 0.5 ng/ml bFGF (Peprotech)). The next day, cells were treated with TGF-β (2.5 ng/ml) for 24h. Finally, viable (7-AAD^-^) transduced (BFP^+^) Cas9 (GFP^+^) fibroblasts were FACS-sorted and processed for qPCR analysis or Indel-seq validation.

## Data availability

The data used in and generated from this publication have been deposited in the Gene Expression Omnibus (GEO) database (accession IDs GSE261742 for murine data and GSE280438 for human count matrices) and in the European Genome-Phenome Archive (EGA) database (accession number EGAS50000000835). All data is available from https://doi.org/10.5281/zenodo.14794723.

## Code availability

Code for performing the analyses and generating all figures is available from GitHub (https://github.com/csbg/fibroblast-perturb-seq-analysis) and has been archived to Zenodo (https://doi.org/10.5281/zenodo.14717236).

## Acknowledgments

The CRISP-seq-BFP backbone (stock Addgene #85707) was produced by I. Amit, Weizmann Institute of Science. pMD2.G (stock Addgene #12259) and psPAX2 (stock Addgene #12260) were produced by D. Trono, École Polytechnique F.d.rale de Lausanne. LV-Cas9-GFP (stock Addgene #82416) was produced by D. Feldser, University of Pennsylvania. Schematic depictions for Figures 1-6, and Extended Data Figure 1 were created with BioRender.com. This research was funded by the Instituto Salud Carlos III (ISCIII) and co-funded by the European Regional Development Fund-FEDER (PI19/00501, PI22/00029), MCIN/AEI/10.13039/501100011033/ European Union (M-Era.Net 2022), and Gobierno de Navarra (20-2022) to BP, Gobierno de Navarra 0011-0537-2019-000012 to LPA and Gobierno de Navarra Doctorados Industriales 0011-1408-2021-000013 to NG. AG is funded by Instituto de Salud Carlos III (CIBERCV CB16/11/00483, and PI21/01999 co-financed by FEDER ERDF funds) and the European Commission H2020 Programme (CRUCIAL project 848109).

## Extended Data Figures

**Extended Figure 1.**
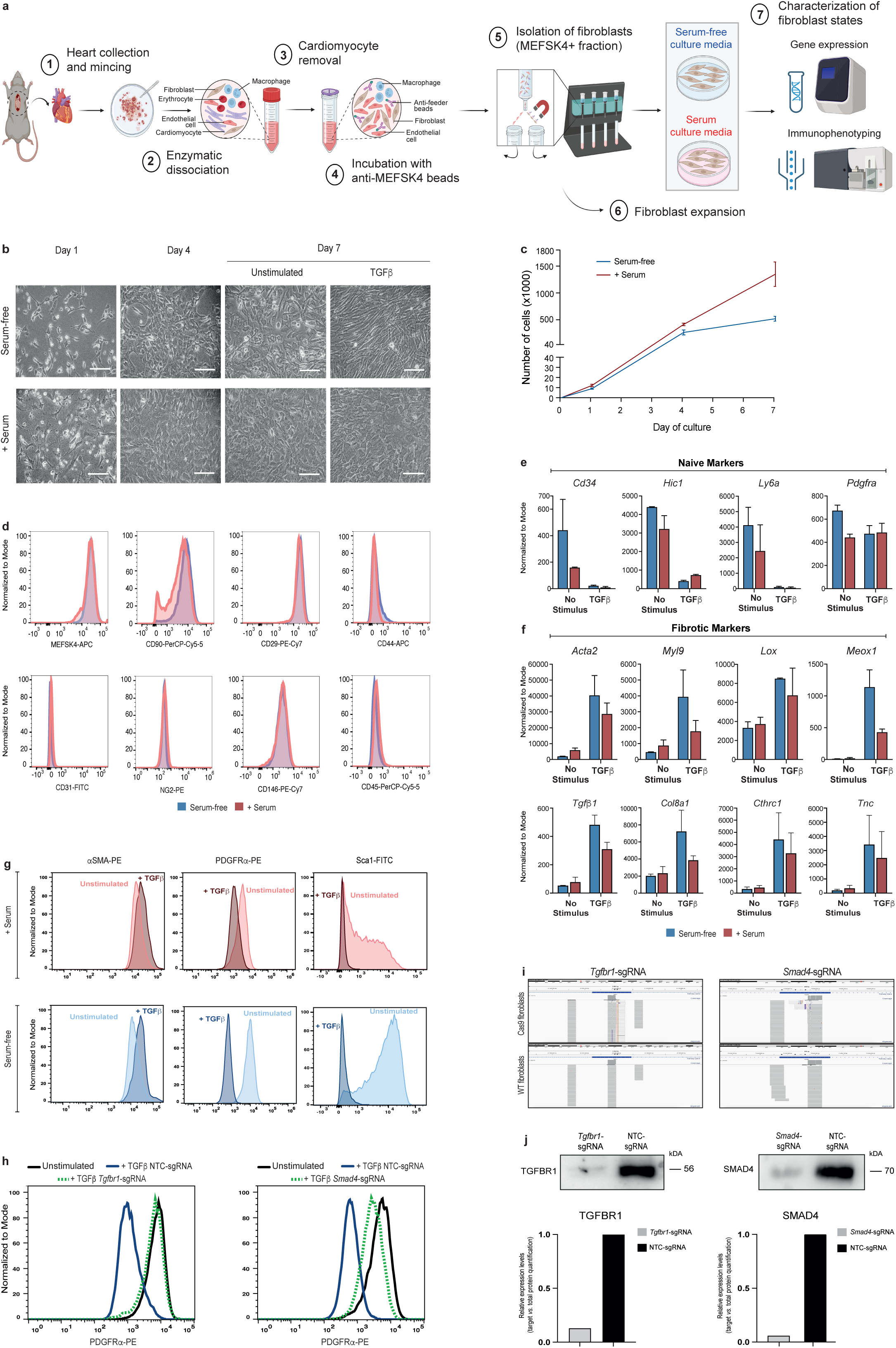
Characterization of the fibroblast *ex vivo* screen system. **(a)** Schematic depiction of *ex vivo* murine cardiac fibroblast expansion. **(b)** Images comparing the morphology of primary murine cardiac fibroblasts in serum-free and serum-containing culture media at days 1, 4 and 7 of culture, and after 24h of TGF-β stimulation at day 7. Scale bars: 30 μm. **(c)** Cell proliferation curves of cardiac fibroblasts grown in both culture media. n=3/time-point. **(d)** Representative FACS analysis of fibroblast-stromal (MEFSK4, CD90, CD44, CD29), pericytes (CD146, NG2), endothelial (CD31) and hematopoietic (CD45) cell markers in fibroblasts grown under serum-free (blue) and serum-containing (red) conditions at day 4 of culture (n=3). **(e-f)** Expression levels of naïve **(e)** and fibrotic **(f)** markers in serum-free (blue) and serum-containing (red) conditions. Values are normalized read counts from two biological replicates. Data are mean ± SD. **(g)** Representative FACS analysis of intracellular alpha smooth actin (α-SMA), PDGFRα and Sca-1 in unstimulated and TGF-β stimulated fibroblasts grown under serum-free (blue) and serum-containing (red) conditions (n=3-4). **(h)** Representative FACS analysis for PDGFRα in Cas9 fibroblasts transduced with NTC sgRNA (blue) and *Smad4* and *Tgfbr1* sgRNAs (green). PDGFRα signal in unstimulated conditions (black) is shown for reference (n=2-4). **(i)** Genome browser snapshots of Indel-seq signal for *Tgfbr1* and *Smad4* loci in Cas9 and wildtype (WT) murine cardiac fibroblasts transduced with *Tgfbr1*- or *Smad4*-sgRNAs. **(j)** Western blot of Tgfbr1 and Smad4 in Cas9 fibroblasts transduced with the target-gene or control (NTC) sgRNAs. Relative expression levels were calculated using total protein content loaded stained with Revert700 as a normalization factor.

**Extended Figure 2.**
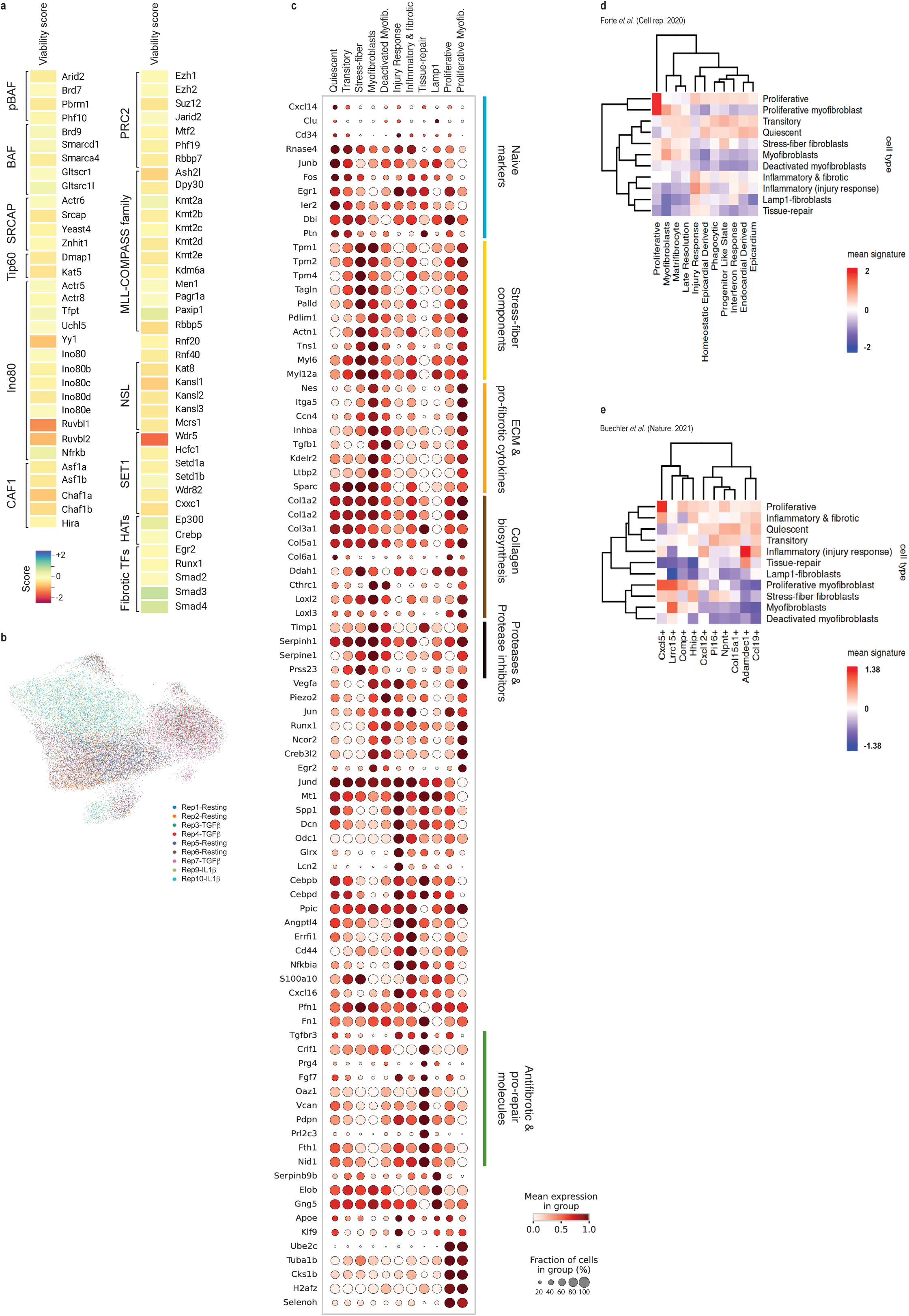
Single-cell perturbation characterization of *ex vivo* fibroblast cultures. **(a)** Heatmap showing viability score of Chromatin Factors grouped by Chromatin Complex membership derived from FACS-based CRISPR screen. **(b)** UMAP projection showing the replicates performed for the Perturb-seq screen. **(c)** Plot showing expression of fibroblast markers across fibroblast subpopulations. Markers for naïve fibroblasts, stress-fiber components, ECM and pro-fibrotic cytokines, collagen biosynthesis, proteases and protease inhibitors, antifibrotic and pro-repair molecules and others are included. Dot size represents percentage of cells in the cluster expressing the marker gene. Color represents scaled expression values. **(d-e)** Heatmap showing enrichment of *in vivo* expression signatures shown in Forte *et al.*^13^ **(d)** and Buechler *et al*^11^. **(e)** over *ex vivo* fibroblast populations from this study.

**Extended Figure 3.**
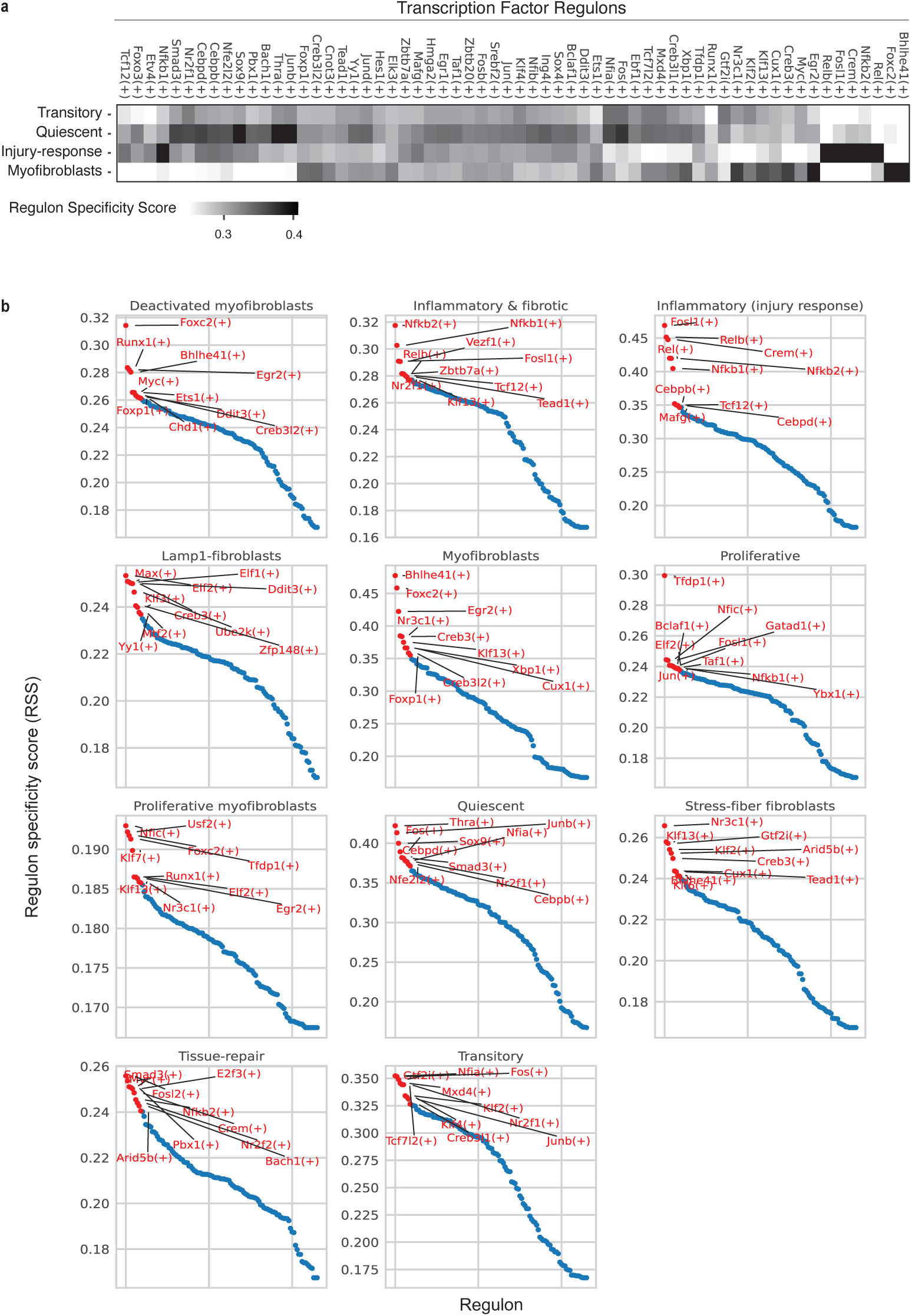
SCENIC analysis of TF regulons active in the *ex vivo* fibroblast subpopulations characterized in this study. **(a)** SCENIC analysis of TF regulons active in the *ex vivo* fibroblast subpopulations characterized in this study. **(b)** TF regulons are ordered decreasingly by their regulon specificity scores. The top 10 regulons for each subpopulation are labelled in red.

**Extended Figure 4.**
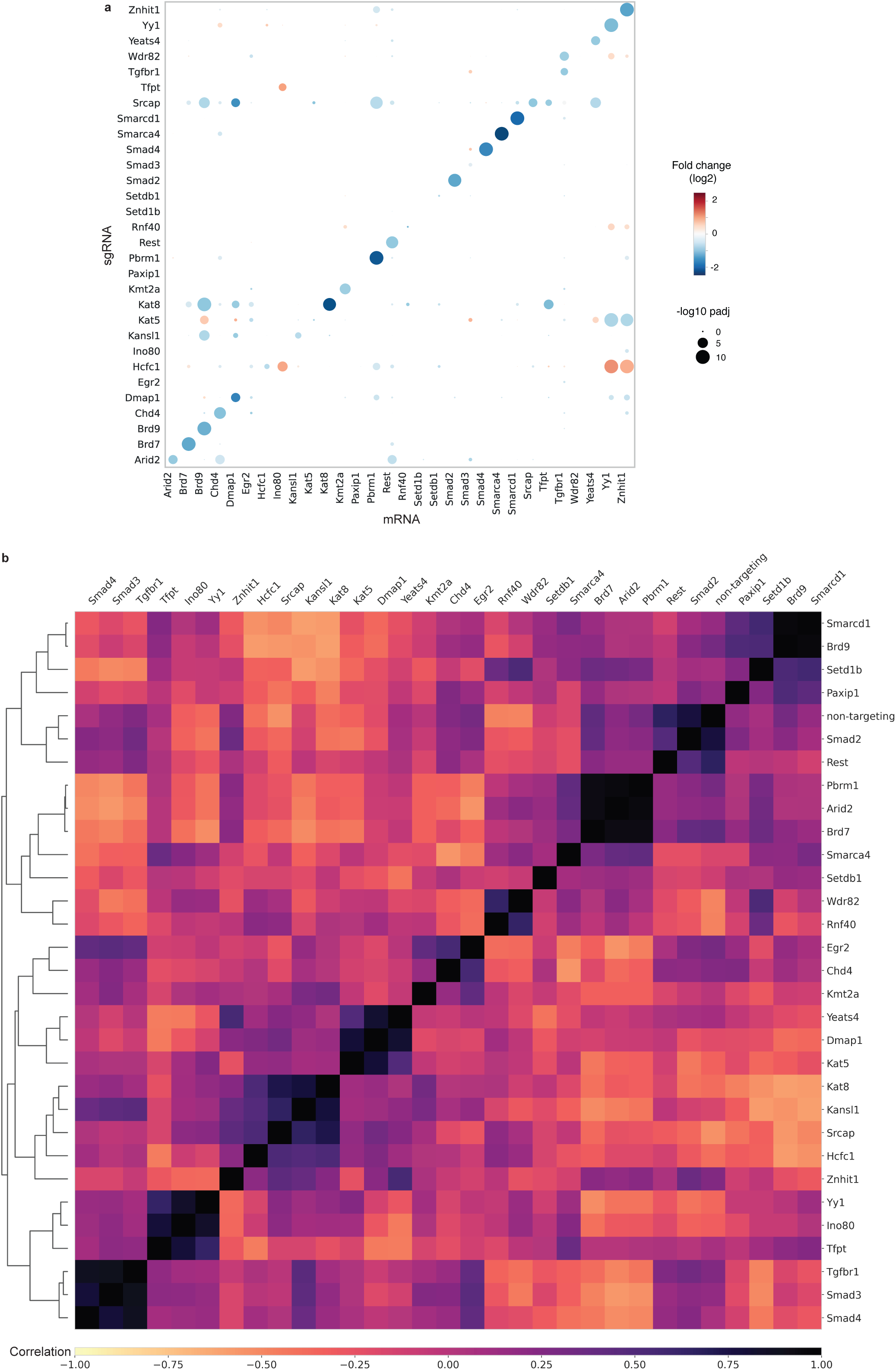
Single-cell perturbation analysis of epigenetic regulators in primary fibroblasts. **(a)** Effect of epigenetic perturbations (Y-axis) on the expression of their target genes (X-axis). Differential gene expression in each epigenetic regulator KO was calculated compared to NTC and is represented as log2 fold change value. Dark blue to dark red colour gradient denotes lower to higher log2 fold changes. **(b)** Correlation plot of gene expression profiles generated through the epigenetic perturbations in primary fibroblasts. Dendrogram showing hierarchical clustering of epigenetic regulator KOs is shown on the left side of the plot. White to black colour gradient denotes lower (-1) to higher (1) coefficients of Pearson.

**Extended Figure 5.**
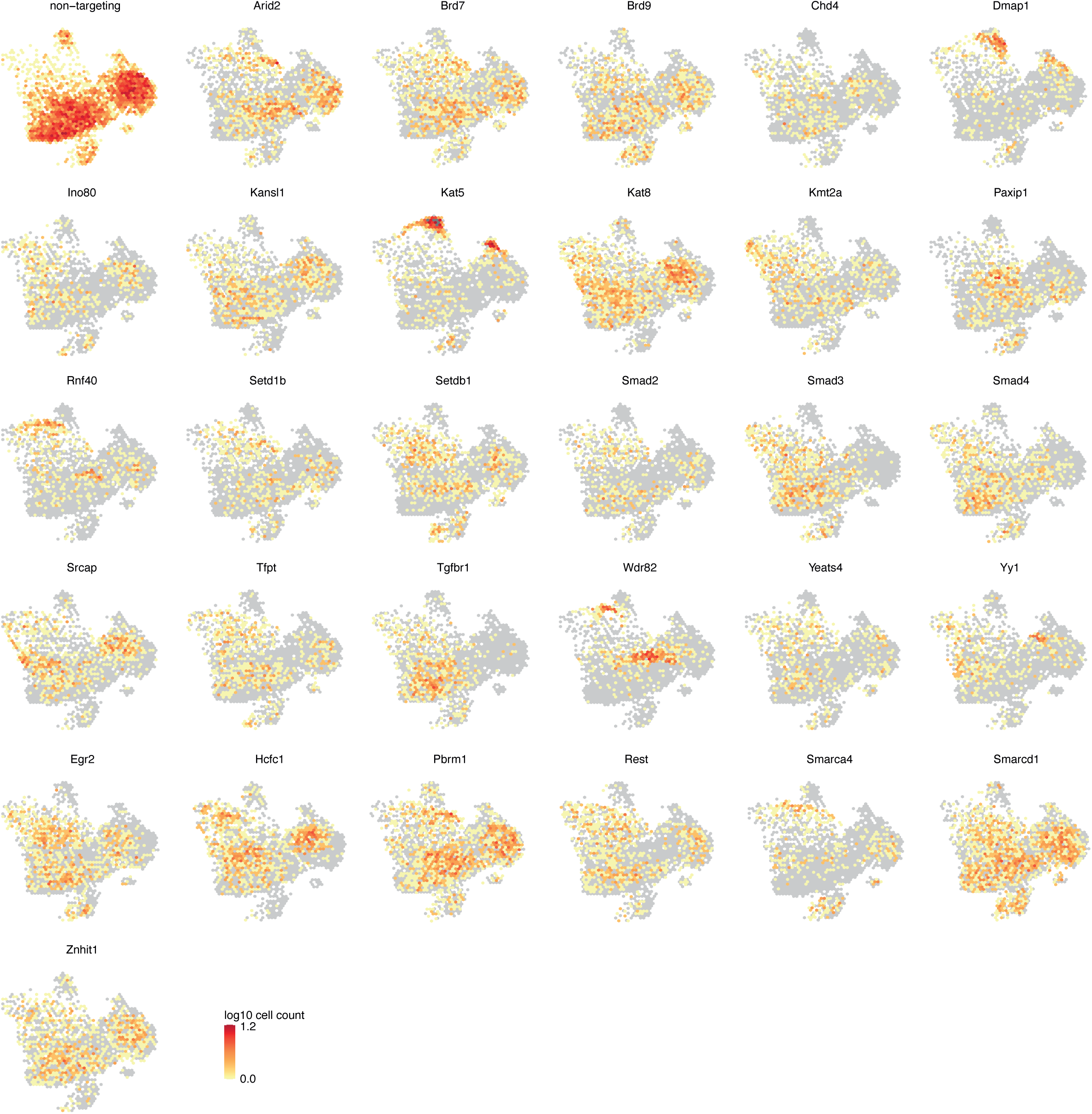
UMAPs showing the distribution of unperturbed fibroblasts (non-targeting) and *Tgfbr1*, *Smad2*, *Smad3*, *Smad4* and ChrF-KOs in *ex vivo* fibroblasts.

**Extended Figure 6.**
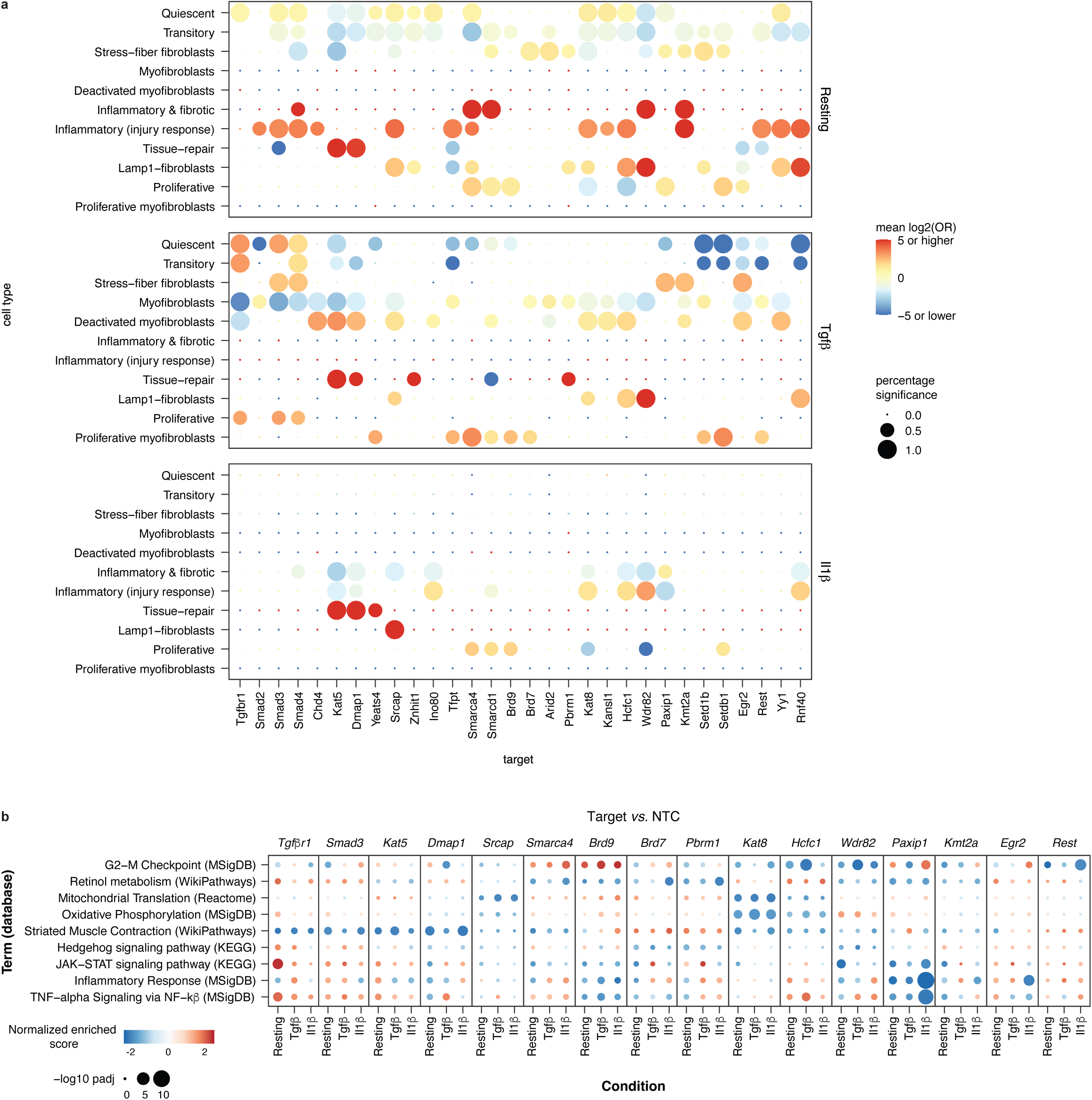
Single-cell perturbation analysis of epigenetic regulators in primary fibroblasts identifies regulators of specific fibroblast states. **(a)** Enrichment analysis of cells with specific epigenetic perturbations across the different fibroblast states identified in resting, TGF-β and IL-1β conditions. Dot color and size relate to the log2 odds ratio and the percent of significant enrichments (one test was performed per NTC), respectively. The analysis is based on measurements of two merged sgRNAs per target. **(b)** Gene set enrichment analysis (GSEA) of differentially expressed genes in across representative knockouts. The color of each dot represents normalized enrichment score, the size represents the –log10 adjusted p-value.

**Extended Figure 7.**
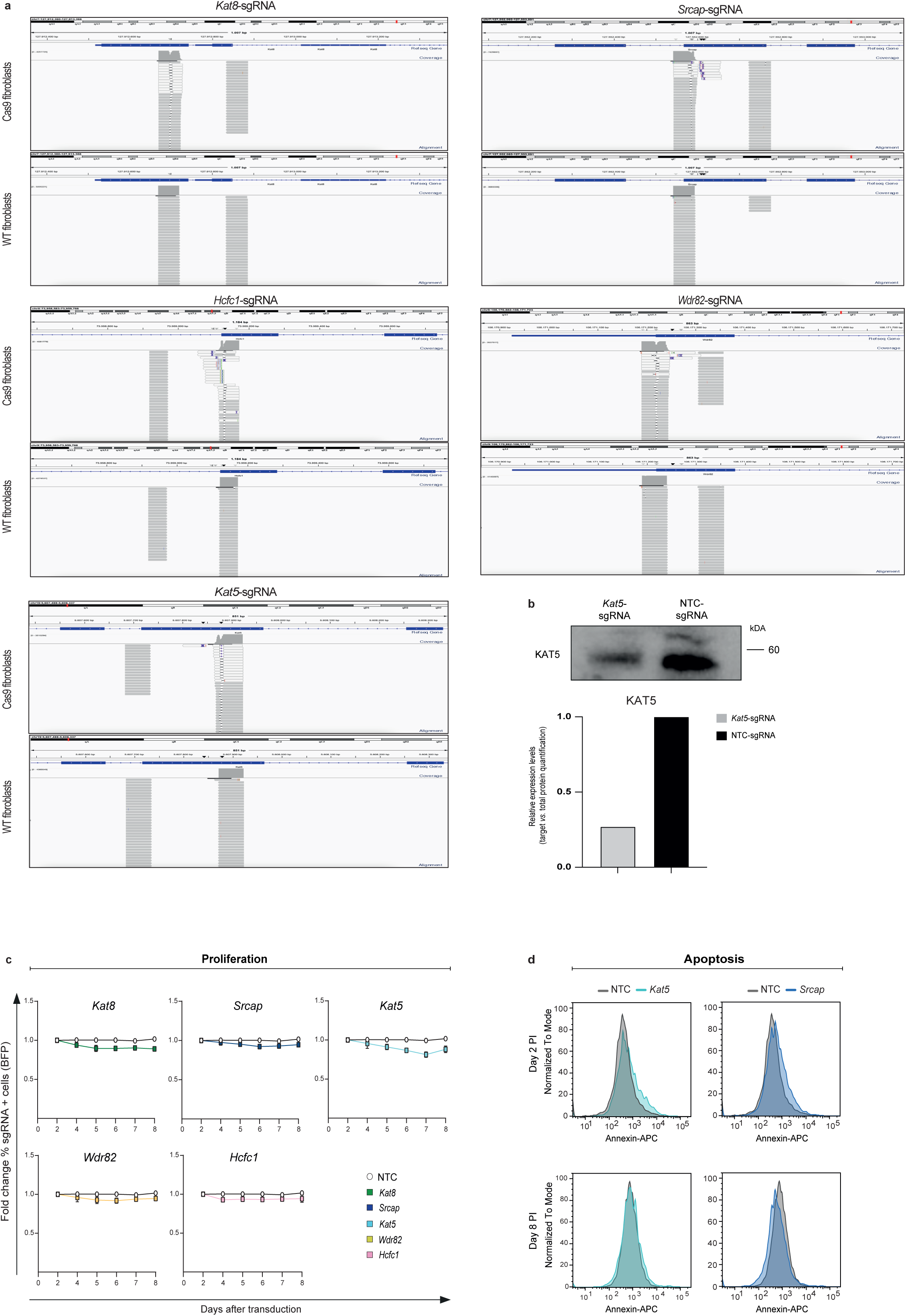
Validation of the knockout produced with the different guides by CRISPR. **(a)** Genome browser snapshots of Indel-seq signal for *Kat8, Srcap, Hcfc1, Wdr82* and *Kat5* loci in Cas9 and wildtype (WT) murine cardiac fibroblasts transduced with *Kat8-, Srcap-, Hcfc1-, Wdr82-* or *Kat5*-sgRNAs. **(b)** Western blot of Kat5 in Cas9 fibroblasts transduced with the target-gene or control (NTC) sgRNAs. Relative expression levels were calculated using total protein content loaded stained with Revert700 as a normalization factor. **(c)** Proliferation curves of control fibroblasts and *Kat8-*, *Srcap*, *Kat5-Wdr82* and *Hcfc1*-KO fibroblasts. The assay measures the change in the proportion of BFP (sgRNA) expressing cells over time performed in three replicate experiments. Statistical significance was analyzed by two-way ANOVA test. No significant differences were observed. Error bars are SD. **(d)** Representative FACS plots showing apoptosis levels (Annexin V + 7AAD^-^) in unperturbed (NTC) and *Srcap* and *Kat5*-KO *ex vivo* cardiac fibroblasts (n=2).

**Extended Figure 8.**
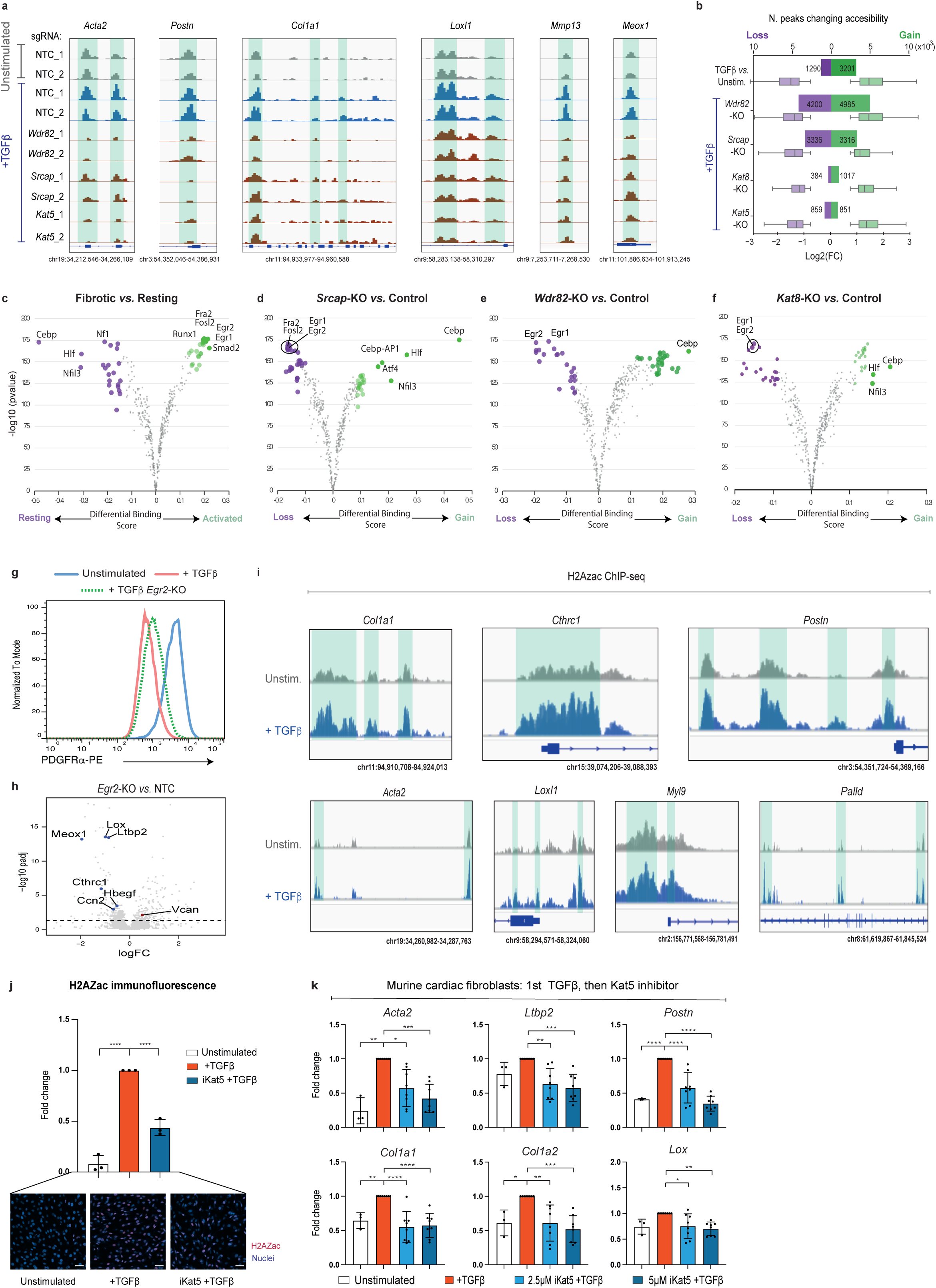
Epigenetic mechanisms underlying candidates’ pro-fibrotic roles. **(a)** Genome browser snapshots of ATAC-seq signal for fibrotic loci in cardiac fibroblasts: unstimulated (grey), 24h TGF-β-stimulated (blue) and *Wdr82*, *Srcap* and *Kat5* knockouts in TGF-β stimulated conditions (red). Peaks for two replicate samples of each condition are shown. **(b)** Plot showing number of changing peaks (log2FC >1) and fold-change distribution in unstimulated *vs.* TGF-β-stimulated (24h) unperturbed fibroblasts and in *Wdr82, Srcap, Kat8* and *Kat5* knockouts. **(c)** Volcano plot showing differential TF motif footprints between TGF-β-stimulated (24h) and resting conditions. **(d-f)** Volcano plot showing differential TF motif footprints between conditions in TGF-β-stimulated (24h) conditions: **(d)** *Srcap* knockout *vs.* control fibroblasts (NTC), **(e)** *Wdr82* knockout *vs.* control fibroblasts (NTC), **(f)** *Kat8* knockout *vs.* control fibroblasts (NTC). Differential footprinting analysis was performed with TOBIAS **(g)** Representative validation of *Egr2*-KO using PDGFRα FACS readout in Cas9 fibroblasts grown under TGF-β conditions (n=2). **(h)** Volcano plot showing differentially expressed genes derived from Perturb-seq between *Egr2*-KO and control (NTC) fibroblasts in TGF-β-stimulated (24h) conditions. **(i)** Genome browser snapshots of H2AZac ChIP-seq signal at representative fibrotic loci in unstimulated and TGF-β-stimulated (24h) fibroblasts. **(j)** Quantification of the percentage of H2AZac positive cells, following TGF-β stimulation in control and NU-9056 pre-treated murine cardiac fibroblasts. The measurements were performed in three replicate experiments. Representative images of H2AZac immunostaining assays are shown at the bottom. Scale bars: 50 μm. **(k)** Expression of fibrotic marker gene expression in unstimulated control or following TGF-β stimulation for 24h, followed by treatment with NU-9056 or vehicle for 24h in murine cardiac fibroblasts. The measurements were performed in three-eight replicate experiments.All data are shown as fold change *vs.* TGF-β values and are mean ± SD. Statistical significance was analyzed by one-way ANOVA or Kruskal-Wallis tests and is indicated as follows: *P <0.05, **P<0.01, ***P<0.001, ****P <0.0001.

**Extended Figure 9.**
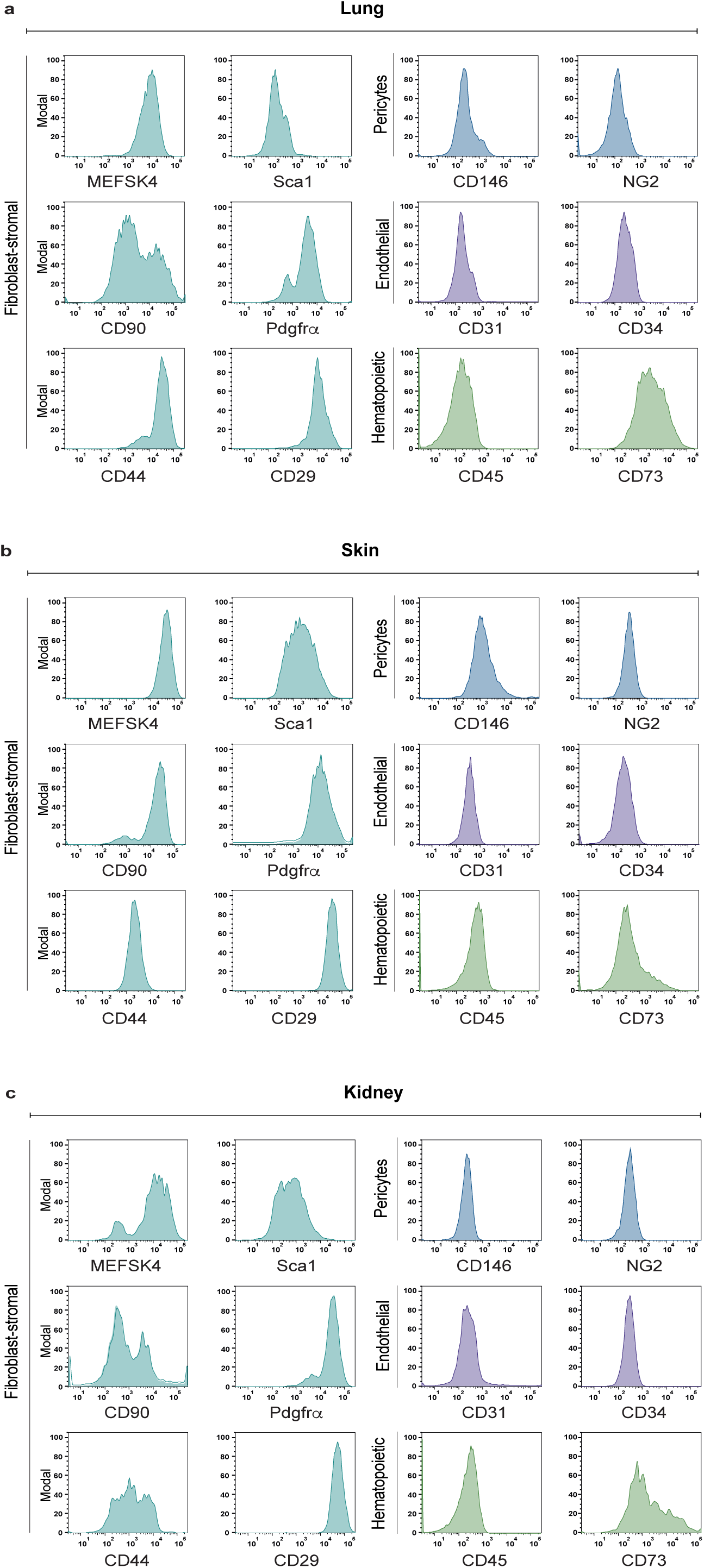
Characterization of lung, skin, and kidney fibroblasts through flow cytometry. **(a-c)** Representative FACS analysis of fibroblast-stromal (MEFSK4, CD90, CD73, CD44, CD29), pericytes (CD146, NG2), endothelial (CD31, CD34) and hematopoietic (CD45) cell markers in unstimulated lung **(a)**, skin **(b)** and kidney **(c)** fibroblasts at day 7 of culture (n=3).

**Extended Figure 10.**
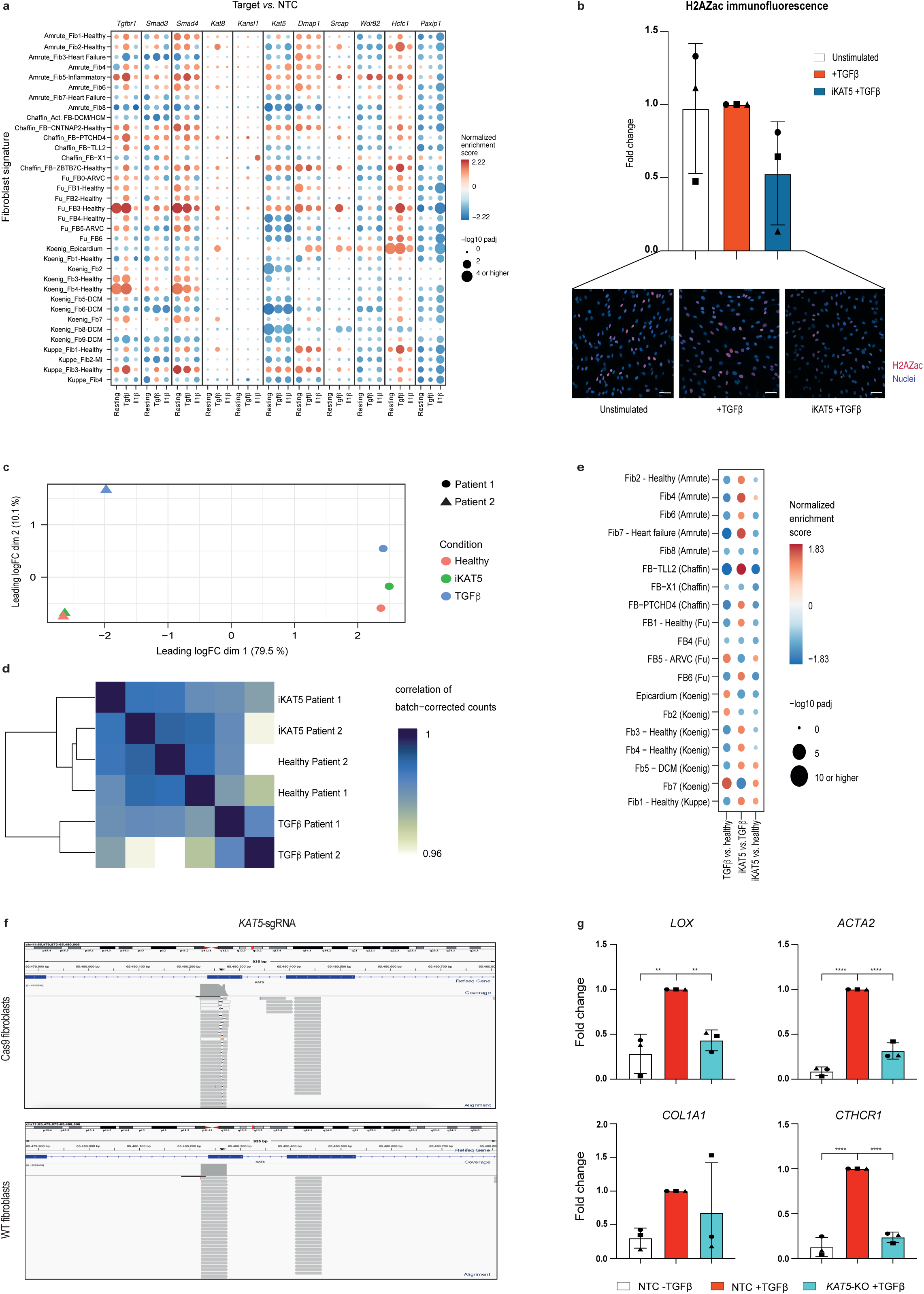
KAT5 chemical and genetic perturbation attenuates fibrotic responses in human primary cardiac fibroblasts. **(a)** Gene set enrichment analysis of differentially expressed genes across representative knockouts. Fibroblast expression signatures are taken from references in Fig. 6a. The color of each dot represents normalized enrichment score, the size represents the –log10 adjusted p-value. **(b)** Quantification of the percentage of H2AZac positive cells, following TGF-β stimulation in control and NU-9056 pre-treated human cardiac fibroblasts. The measurements were performed in three patients. Representative images of H2AZac immunostaining assays are shown at the bottom. Scale bars: 50 μm. **(c)** Principal component analysis of gene expression in patient-derived cardiac fibroblasts. Dimension 1 corresponds to variability of gene expression between patients, while dimension 2 captures effects of TGF-β or KAT5 inhibitor on the transcriptome. **(d)** Correlation analysis of gene expression demarcates samples treated with TGF-β from unstimulated and KAT5-inhibited samples. **(e)** Gene set enrichment analysis of differentially expressed genes across treatments. Fibroblast expression signatures are taken from references in Fig. 6a. The color of each dot represents normalized enrichment score, the size represents the –log10 adjusted p-value. **(f)** Genome browser snapshots of Indel-seq signal for *KAT5* loci in Cas9 and wildtype (WT) human cardiac fibroblasts transduced with *KAT5*-sgRNA. **(g)** Gene expression analysis of fibrotic markers in human cardiac fibroblasts depleted for KAT5 using CRISPR/Cas9 and then stimulated with TGF-β. The measurements were performed in cell cultures from three different patients. Each patient is represented by a different symbol. All data are shown as fold change *vs.* TGF-β values and are mean ± SD. Statistical significance was analyzed by one-way ANOVA test and is indicated as follows: **P<0.01, ****P <0.0001.

**Supplementary Figure 1.**
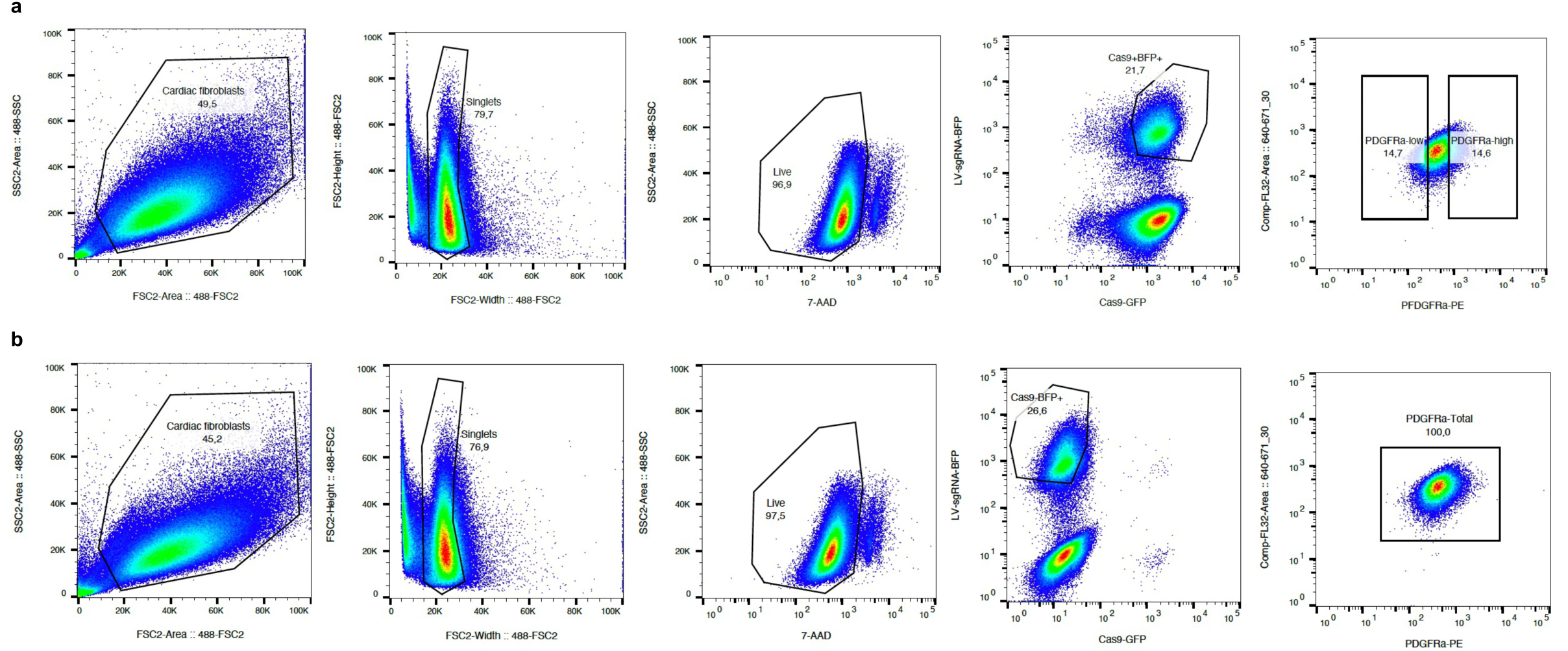
FACS Readout for Bulk CRISPR screens. **a)** Gating strategy for separately sorting naïve and fibrotic Cas9 cardiac fibroblasts. **b)** Gating strategy for sorting non-Cas9 (wildtype) cardiac fibroblasts.

**Supplementary Figure 2.**
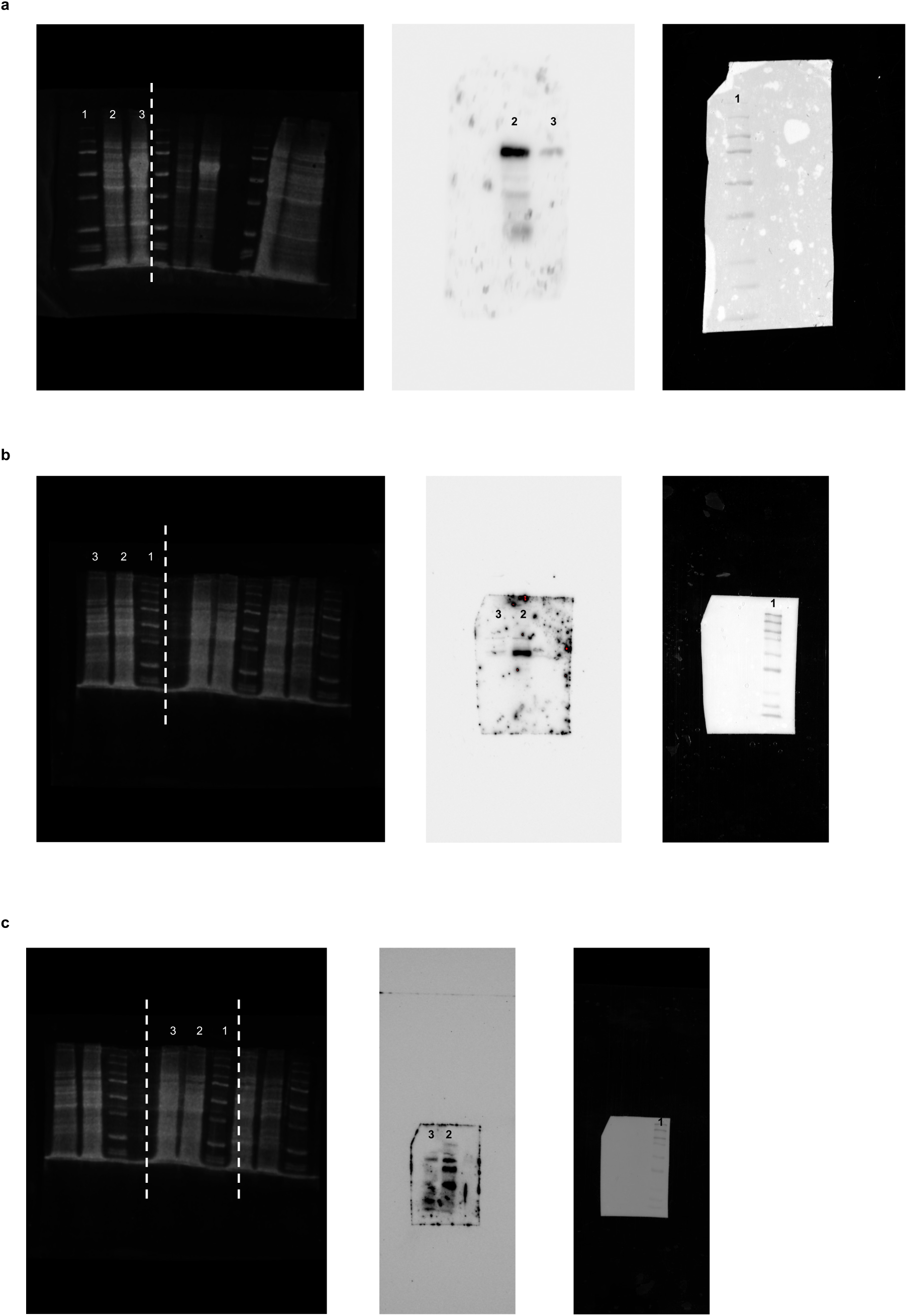
Uncropped versions of blots for total protein (left), target protein (middle) and Precision Plus Protein Standard (right) of Smad4 (a), Tgfbr1(b) and Kat5 (c) Western Blots. In all the blots shown, lane 1 corresponds to Precision Plus Protein Standard loaded (#161-0376, Bio-Rad), lane 2 corresponds to NTC-sgRNA sample loaded and lane 3 corresponds to target-sgRNA sample loaded.

**Supplementary Figure 3.**
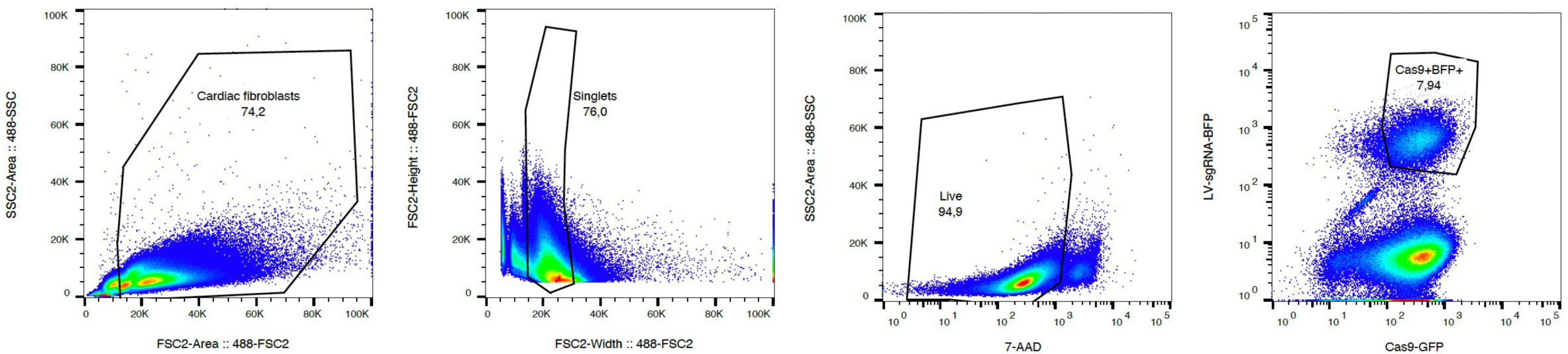
Perturb-seq gating strategy of Cas9 cardiac fibroblasts.

