## Supplementary Table 1 for "Identification of epigenetic regulators of fibrotic transformation in cardiac fibroblasts through bulk and single-cell CRISPR screens"

**Supplementary Table 1. List of chromatin factors and sgRNAs sequences analyzed in FACS-based bulk CRISPR screens.**

| gRNA sequence | gRNA ID | Gene ID |
|---|---|---|
| ATGTCTGGGACGTCCCACGA | Acta2_1 | Acta2 |
| CCTACCAGTTGTACGTCCAG | Acta2_2 | |
| TGAGGTAGTCGGTGAGATCT | Acta2_3 | |
| GTCCCAGGTTGATTTCCCCA | Actl6a_As_76114 | Actl6a |
| CCTTCTCTAGTGGAACACCA | Actl6a_As_76115 | |
| GCACCACACCGATAGCCGTG | Actl6a_As_76116 | |
| CGAATGCTACAGGATCCCGA | Actr5_As_76054 | Actr5 |
| GCTGAAGTATCCAGGCCACC | Actr5_As_76055 | |
| GGAGCATAGCTACATTGCCG | Actr5_As_76056 | |
| TGATGCAGCACAGCTCAGAA | Actr6_As_76050 | Actr6 |
| GTAAGGAACAATGTGCGTGA | Actr6_As_76051 | |
| TTTGGATCTGAACTGACAAT | Actr6_As_76052 | |
| AGATGTCAAACGGTACAAGG | Actr8_As_76046 | Actr8 |
| AGCCAGTTGTCCTTGTACAG | Actr8_As_76047 | |
| GAATGTTGTAACAGTCCAGG | Actr8_As_76048 | |
| CATGAGAGGAGTTTAAACCG | Ahctf1_R2.Br_57108 | Ahctf1 |
| CGACACGAACCGTCAAACAG | Ahctf1_R2.Br_57109 | |
| GAGCCTATAATTAACCATGG | Ahctf1_R2.Br_57110 | |
| TACTTACAATTTGCTAACAG | Ankra2_R2.Br_33062 | Ankra2 |
| CTGGCTACTCGTATTGAACA | Ankra2_R2.Br_33063 | |
| TGCCGCAGCCCACATCAGGG | Ankra2_R2.Br_33064 | |
| AGGAGCCTGGGATTTCACGG | Ankrd32_As_74146 | Ankrd32 |
| TTAAGTTTAATATTCCTCTA | Ankrd32_As_74147 | |
| ACAGCCTGTATACAATGTAG | Ankrd32_As_74148 | |
| GTTCTTCAAACTCATCCGTG | Anp32a_As_73962 | Anp32a |
| AAGCGAAAACAGAATCTCAG | Anp32a_As_73963 | |
| AGGCAAAATCGAAGGCCTCA | Anp32a_As_73964 | |
| GGTTCTGCTTACGGCTGCCG | Anp32b_As_73958 | Anp32b |
| TCAACTTAGGTAGCTTCGGG | Anp32b_As_73959 | |

| | | |
|---|---|---|
| CTTGGACAATTGCAAAGCAA | Anp32b_As_73960 | |
| GGAGTTGAAGAACAGAGCCC | Anp32e_As_73954 | Anp32e |
| CCTCTTCATAGCCTTCCGGT | Anp32e_As_73955 | |
| GAGTCATCTCACCTCCTCCG | Anp32e_As_73956 | |
| GCAGCTGCGAAGATATCGGG | Arid1a_As_73073 | Arid1a |
| TGGCTACCAGGGCTACCCCG | Arid1a_As_73074 | |
| GAGTGCATCTGTCCTCCAGA | Arid1a_As_73075 | |
| GTACCCATCCCATACAACTG | Arid1b_As_73069 | Arid1b |
| GATTCTGACTGGCTTCCAGG | Arid1b_As_73070 | |
| GCTCAGCACCCCGTACCCCG | Arid1b_As_73071 | |
| CTTTACTGCTCGCTAATGCG | Arid2_As_73065 | Arid2 |
| TTAATGCCCAGCTTTCGAGG | Arid2_As_73066 | |
| GAGTGGTTCTGAAATCCACA | Arid2_As_73067 | |
| TTAATCTTCCAGATTGACAG | Arid4a_As_73049 | Arid4a |
| TGTCAGTGCCAAGTACCGAG | Arid4a_As_73051 | |
| AGACTGGTCTATCATCAGGG | Arid4a_TF2.Br_61181 | |
| ACAGTCCCCAGAACCTAAGA | Arid5a_As_73041 | Arid5a |
| GCTGCCCTGAGGCCTACAAG | Arid5a_As_73042 | |
| TGAGAAGCTGAAGAAGGCCA | Arid5a_As_73043 | |
| ATAGGTTTACCACCTCGCCA | Arid5b_As_73037 | Arid5b |
| GGGCTCTCTGTAAAGCAGCA | Arid5b_As_73038 | |
| GTACTCGCAGACACTACGAA | Arid5b_As_73039 | |
| GCATAGCCGGGAGGGCCCAG | Arntl_As_72846 | Arntl |
| AGGCTCTTACTGGTAGTCAG | Arntl_As_72847 | |
| CCCACAGTCAGATTGAAAAG | Arntl_As_72848 | |
| ACTCACCCACGGGGTCCACG | Arrb1_As_72798 | Arrb1 |
| GAGTATCTCAAAGAAAGGCG | Arrb1_As_72799 | |
| CCGACCAGCACTCACCCACG | Arrb1_As_72800 | |
| CTGATTACTTGCACCTACCG | Asf1a_B_72578 | Asf1a |
| AATAAATTCTTGACCTCGGT | Asf1a_B_72579 | |
| AACGGGTTGTAGAAAGGCGA | Asf1a_B_72580 | |

| | | |
|---|---|---|
| GGTTCAGGTGAACAATGTAG | Asf1a_B_72581 | |
| ATTCGAAGCTGATCTCGAAC | Asf1b_B_72574 | Asf1b |
| GATCAGCTTCGAATGCAGTG | Asf1b_B_72575 | |
| AAGCTGATCTCGAACCGGAA | Asf1b_B_72576 | |
| TATGCCTTCCTGCAGGGACA | Asf1b_B_72577 | |
| GCCACCACTTTAGTCCACCG | Ash1l_B_72562 | Ash1l |
| AGGACGGTAGAAGTTTCAGG | Ash1l_B_72563 | |
| GCATCCATCTTCATTCACAG | Ash1l_B_72564 | |
| CCTGACTTAAAAAAGAAACG | Ash1l_B_72565 | |
| AGAAGTGATGGATACCCAGG | Ash2l_As_72558 | Ash2l |
| CTGCCACGTCAAATTCGCCA | Ash2l_As_72559 | |
| CTGTGGATGAGGAGAATGGG | Ash2l_As_72560 | |
| TGAGTGTGAAAAGACTAATG | Asxl1_R1.Br_57764 | Asxl1 |
| GTGTTAACAAGAATGGACCC | Asxl1_R1.Br_57765 | |
| TACTAAAACCGACTTAGCAG | Asxl1_R1.Br_57767 | |
| AGTCAGTCAGAACCGACATG | Asxl2_R1.Br_41526 | Asxl2 |
| GTCAGTCAGAACCGACATGA | Asxl2_R1.Br_41527 | |
| AGAACCGACATGAGGGAAAA | Asxl2_R1.Br_41528 | |
| ACGCCACTCTCTTATCCCCT | Atad2_As_72430 | Atad2 |
| TCTTAGGCAGAGAAAAACAA | Atad2_As_72431 | |
| CCATAGACCCTGCTTTACGG | Atad2_As_72432 | |
| CCTGGTGCTACATTACCAAA | Atad2b_As_72426 | Atad2b |
| CATCTTGCAGATCCACACCA | Atad2b_As_72427 | |
| GTTTGGCTGGAGGAGATACG | Atad2b_As_72428 | |
| ACTCGTCCCGAGCAACCAGG | Atf1_1 | Atf1_ |
| GATCCCGTGAGCTTTCTGTG | Atf1_2 | |
| TGTACTATCTGAGAAACGTG | Atf1_3 | |
| CCAGCGCAGAGGACATCCGA | Atf3_1 | Atf3 |
| TATACATGCTCAACCTGCAC | Atf3_2 | |
| TCAAATACCAGTGACCCAGG | Atf3_3 | |
| ATCACATGTGTCATCCAACG | Atf4_1 | Atf4 |

| | | |
|---|---|---|
| CAAACCCGACTGGTCGAAGG | Atf4_2 | |
| CAAACCCGACTGGTCGAAGG | Atf4_3 | |
| AACCACAGAGACGACCCATG | Atf6_1 | Atf6 |
| GAACACGAGTCTGTGGACCG | Atf6_2 | |
| GAGCAGACGACAGAACCGCA | Atf6_3 | |
| CTAGAAGCAGGACTGTCAGG | Atf7_1 | Atf7 |
| GTAGGGGTAGAGATCACAAC | Atf7_2 | |
| ATAAGATTGAGAAATCTCCA | Atm_As_72274 | Atm |
| TGAAAAATCACACAAGGCGG | Atm_As_72275 | |
| ATAGTGACAAACCTAGGCCA | Atm_As_72276 | |
| GTAGGGCCTTAAAGACCCCA | Atn1_As_72266 | Atn1 |
| ATATCGCATTGGAGGCCCAG | Atn1_As_72267 | |
| GCCCAAAAAGACCAAAACCG | Atn1_As_72268 | |
| GCCAAGCTCTGAGATGCGGG | Atoh8_1 | Atoh8 |
| GCGACGCGGAAGAATCCGGG | Atoh8_2 | |
| TCTCCAAACTGGCCATCCTG | Atoh8_3 | |
| CTACCTGGCCAACTCCCGCA | Atr_As_71906 | Atr |
| CCTTCTGCCATTAAGCCAGA | Atr_As_71907 | |
| GCTCAACAGGAGAATCTCAG | Atr_As_71908 | |
| GCTGACAATCTCAAAAAACG | Atrx_As_71886 | Atrx |
| ATCTGATGATGAAAAACCCA | Atrx_As_71887 | |
| GACAATCTCAAAAAACGCGG | Atrx_As_71888 | |
| CCGCTGCTCCGCCCGCCGCG | Atxn7_As_71858 | Atxn7 |
| CTGCTGCTGCTGCTGCCGGG | Atxn7_As_71859 | |
| AGACACATTCCTTAACCCAG | Atxn7_As_71860 | |
| GATTCTGCTTTGAAGTACAC | Atxn7l3_As_71846 | Atxn7l3 |
| TGGATCTTCGAGGGGAATTA | Atxn7l3_As_71847 | |
| AGCGGGAAGCAGCAATACTG | Atxn7l3_As_71848 | |
| TTTCGGACCGAAGGGAACAG | Aurka_As_71802 | Aurka |
| TCTCAGACCACTGTTCCCTT | Aurka_As_71803 | |
| GCAGATCCTAGGTTCTGAGA | Aurka_As_71804 | |

| | | |
|---|---|---|
| CTTACCGTCTTTGAGCCGTA | Aurkb_As_71794 | Aurkb |
| CTTGTTCTGGGATCCTTGCG | Aurkb_As_71795 | |
| GCAAGGATCCCAGAACAAGT | Aurkb_As_71796 | |
| CTGATCCTGATACCCAACCA | Bach1_1 | Bach1 |
| TCTAAGCGTACACAATATCG | Bach1_2 | |
| TGCTCAGCCTCAATGACCAG | Bach1_3 | |
| AATTACGGACAGCCCCACGT | Bach2_1 | Bach2 |
| CTCCTCGTATTCCTACGCAG | Bach2_2 | |
| TGCGCAGGAACTCAGCACAG | Bach2_3 | |
| GTAGTGGCGGAAGATCTCGT | Baz1a_B_71346 | Baz1a |
| CATCGCCGAAGACTTCAGGA | Baz1a_B_71347 | |
| GGATAAAGAGAAAAAAAGGG | Baz1a_B_71348 | |
| GCATGGCCACAGTTACACCA | Baz1a_B_71349 | |
| GTTCACTGTATCTTTCCAGG | Baz1b_B_71342 | Baz1b |
| CTACCAGCCTGAATGCTGGG | Baz1b_B_71343 | |
| GTACTGGTGAGAATGCACCT | Baz1b_B_71344 | |
| GTATTCCTCCTCTTGGTAGA | Baz1b_B_71345 | |
| ATGAGGCAGAAGGTCCAACG | Baz2a_R1.Br_50492 | Baz2a |
| GCGCAGTTCTCGGATCATGG | Baz2a_R1.Br_50494 | |
| TTGGTGTAAAGACTCCACAG | Baz2a_R1.Br_50495 | |
| GAATCGAGACAATGTGTCCG | Baz2b_R1.Br_73753 | Baz2b |
| GGAGGCCGAGAAACGAACCA | Baz2b_R1.Br_73754 | |
| GGAGAATCCAGTATCTGCGT | Baz2b_R1.Br_73755 | |
| TCAATGATCATAATTCCGTG | Bbx_R2.Br_35854 | Bbx |
| TTAGATCCTACTCAGATGGG | Bbx_R2.Br_35855 | |
| AGAACCCTTGACCCCTACCG | Bbx_R2.Br_35856 | |
| GTGCCAGATGAATTTCCCAC | Bcl11a_As_70959 | Bcl11a |
| TTTCATCTCGATGGGAGAAG | Bcl11a_As_70960 | |
| TCGATGGGAGAAGGGGAAGG | Bcl11a_As_70961 | |
| CGTGGCGGTGGGCTGAAGAG | Bcl11b_As_70955 | Bcl11b |
| CCCACTCATGAATTTCCTGG | Bcl11b_As_70956 | |

| | | |
|---|---|---|
| AGAGCAATCTCATCGTGCAC | Bcl11b_As_70957 | |
| TGATATCAAGAGGGTCATGG | Bcl7a_As_70896 | Bcl7a |
| CCTGGGATGATGGACATGCA | Bcl7a_As_70897 | |
| GCTGTTCTCTTGCTTGATGG | Bcl7a_As_70898 | |
| CCATTTCCGCACTTTCTCGA | Bcl7b_As_70892 | Bcl7b |
| CTGTGGGTGATACCTCCCTG | Bcl7b_As_70893 | |
| TCCCTCCTCCTTGAATTCCA | Bcl7b_As_70894 | |
| GCCTTGGCAGAGGTCACCCG | Bcor_As_70864 | Bcor |
| GTGCAGACAGGAGAATACAG | Bcor_As_70865 | |
| TCAATGGTGCTAGTTACCTG | Bcor_As_70866 | |
| GGTTGACTGCTCGCCAGCCG | Bcorl1_As_70860 | Bcorl1 |
| ACAGTTTCACTGGAACCAAG | Bcorl1_As_70861 | |
| TCTACAGCACCGCTCTACAG | Bcorl1_As_70862 | |
| TTGTATACAAATTAGTCCCA | Bmi1_As_70578 | Bmi1 |
| CCTCATCTGCAACTTCTCCT | Bmi1_As_70579 | |
| ACCACTCCTGAACATAAGGT | Bmi1_As_70580 | |
| CATCCCGTCGATGAAATACA | Bptf_As_70358 | Bptf |
| GCAAAGATAGCAAAGCTAAG | Bptf_As_70359 | |
| CCATGTCCTTCCTTACCAGG | Bptf_As_70360 | |
| GATGGGTCTGTGTATAGGGG | Brcc3_As_70334 | Brcc3 |
| GCTCTGGAGAAATTTCTACA | Brcc3_As_70335 | |
| GCTGTCTGCAGCTTCAACAG | Brcc3_As_70336 | |
| GGCACAGCACGCAATCCGCA | Brd1_R1.Br_70330 | Brd1 |
| GTAGCATGTCAGTTTCCACC | Brd1_R1.Br_70331 | |
| GTTCATCGAGAAGTCAGCCG | Brd1_R1.Br_703321 | |
| CTGGTACAGAAGCCATTGTG | Brd2_R1.Br_5865 | Brd2 |
| AAAAGGCGTTAAACGGAAAG | Brd2_R1.Br_5866 | |
| GTGCCATAGCCACAACATCG | Brd2_R1.Br_5867 | |
| GTGGTGGGGGAGACTCACTC | Brd3_R1.Br_30606 | Brd3 |
| GCAGTACATGCAGAATGTAG | Brd3_R1.Br_30608 | |
| GAAAACTGTCCGAGCACCTG | Brd3_R1.Br_30609 | |

| | | |
|---|---|---|
| ACTGCAATATCTGCTCAGAG | Brd4_R1.Br_25574 | Brd4 |
| TTTGGTACCGTGGATACACC | Brd4_R1.Br_25575 | |
| CCGCCAAGCTGGGTCCTCGG | Brd4_R1.Br_25577 | |
| GGTCGCTTCTGTCTTCGAAG | Brd7_As_70314 | Brd7 |
| TGTCCACCTCATTCTCTGCA | Brd7_As_70315 | |
| GTCGGACCGCCACTTCTACG | Brd7_As_70316 | |
| GGTGGCACAAGCTTAACTGG | Brd8_R1.Br_44308 | Brd8 |
| TTGGAGTCAAGTCTCCAAGT | Brd8_R1.Br_44309 | |
| GAAGTCTCCATACCAGTCTG | Brd8_R1.Br_44310 | |
| TTACGGATGCAATTGCTCCT | Brd9_As_70306 | Brd9 |
| GCTGCATGTTTGAGCCTGAA | Brd9_As_70307 | |
| GGAAAAGCAAAAAATCCATG | Brd9_As_70308 | |
| GACTGCGAGTCACAGATCGG | Brdt_R1.Br_50332 | Brdt |
| GAAGGTTGTCAGGTCTAAGG | Brdt_R1.Br_50333 | |
| ATGGCCCTTCTATAATCCTG | Brdt_R1.Br_50335 | |
| GTACCCCAGGAACACTCTGA | Brwd3_As_70214 | Brwd3 |
| GTTCACTGCAGTAATAACAG | Brwd3_As_70215 | |
| CTACCCAGATCGAAGCCGGT | Brwd3_As_70216 | |
| TGTGGACCAGAAACAAAAGA | Bub1_As_70038 | Bub1 |
| TCTCCCCAGTAGGGAACAAA | Bub1_As_70039 | |
| GCAGGGCATCTGGAAGCCCA | Bub1_As_70040 | |
| CCTTGTCCTTGAAGTCCGTG | Carm1_B_69141 | Carm1_ |
| GAGCCAAAACACATACCGTG | Carm1_B_69142 | |
| AGACCTCCTCTACTTTGCCA | Carm1_B_69143 | |
| ACAACATCCTGAAAACCTGT | Carm1_B_69144 | |
| AACCCATACCATCCAATCTG | Cbfb_1 | Cbfb |
| AACCCATACCATCCAATCTG | Cbfb_2 | |
| CGATCTCCGAGCGACCGTCG | Cbfb_3 | |
| TTACAGTCAGAAAAGCCACG | Cbx1_R1.Br_2217 | Cbx1 |
| GAATATCTTCTAAAGTGGAA | Cbx1_R1.Br_2219 | |
| TGGTGGAAAAAGTTCTTGAT | Cbx1_R1.Br_2220 | |

| | | |
|---|---|---|
| GATCCCATTAGAAAGAAACG | Cbx2_B_68897 | Cbx2 |
| GAGTACCTGGTCAAGTGGCG | Cbx2_B_68898 | |
| AAGAGAGGCAAGAGACCCAG | Cbx2_B_68899 | |
| GTCGTCTTCCTCTTCGTCCG | Cbx2_B_68900 | |
| ACACAGTGCTGATAATACTT | Cbx3_B_68893 | Cbx3 |
| AGTATTTCCTGAAGTGGAAG | Cbx3_B_68894 | |
| TGGACCGTCGTGTAGTGAAT | Cbx3_B_68895 | |
| CCTTCCCATTCACTACACGA | Cbx3_B_68896 | |
| CCAAGGGCTTCACGGCCCCG | Cbx4_B_68889 | Cbx4 |
| GTATTCACCTGCACCACCAG | Cbx4_B_68890 | |
| TGGAGAGCATCGAAAAGAAG | Cbx4_B_68891 | |
| GCTCCTGCCCCTCCCCCCAA | Cbx4_B_68892 | |
| GGATGAGGAGGAATATGTGG | Cbx5_B_68885 | Cbx5 |
| GGGTGAAAACAATAAGCCCA | Cbx5_B_68886 | |
| ATATGTGGTGGAAAAGGTGT | Cbx5_B_68887 | |
| ACATGGGAAAGAAGACCAAG | Cbx5_B_68888 | |
| GCTGGAGTGCAGCTTGGGCG | Cbx6_B_68881 | Cbx6 |
| TGAAGCCCCGGGAGCCCAAG | Cbx6_B_68882 | |
| CCACGACCAGACCCACAAGG | Cbx6_B_68883 | |
| GGGGACGGCGAGACATACGG | Cbx6_B_68884 | |
| ACCTCTCTTCCTATACCCCG | Cbx7_B_68877 | Cbx7 |
| CGTAGGCCATGACAAGGCGA | Cbx7_B_68878 | |
| CTATGGCTGACAGCTCCATG | Cbx7_B_68879 | |
| AGAGAGGTCCGAAACCCAGG | Cbx7_B_68880 | |
| CTCTATGGCCCCAAAAAGCG | Cbx8_R2.Br_21794 | Cbx8 |
| GGGGACAGCTCCAAGAAACG | Cbx8_R2.Br_21795 | |
| ACACGGGGCTACCCCCACCA | Cbx8_R2.Br_21796 | |
| ACAATGACTCAGAGCCCCCA | Ccdc101_As_68833 | Ccdc101 |
| TCGAGAATCAGCAGACACAA | Ccdc101_As_68834 | |
| AGTCTCTGTTGGAAGAGAGG | Ccdc101_As_68835 | |
| AGGAACCCTTGATTGTGACA | Cd200_1 | Cd200 |

| | | |
|---|---|---|
| CACCTACAGCAAAACCCATG | Cd200_2 | |
| GAAACAATGCTTAGCAACAG | Cd200_3 | |
| AAAGACCAGATGCAACCAGG | Cdc73_1 | Cdc73 |
| ACCCAGAGAGTACTACACAT | Cdc73_2 | |
| TCTACTTGGATATCTCAATG | Cdc73_3 | |
| ATATAACTGCCCTTAAACAG | Cdc73_As_67380 | |
| TAGCAGAAGCAAAGAAACCA | Cdc73_As_67381 | |
| GCTGCACGCCGGACATAAAC | Cdc73_As_67382 | |
| GAAGATCAGACTTGAAAGCG | Cdk1_1 | Cdk1 |
| GATACGAGTGTACACACACG | Cdk1_2 | |
| GCACTCCTAACAACGAAGTG | Cdk1_3 | |
| AGTCTCTGTGAAGAACTCGC | Cdk1_As_67224 | |
| AATCCATGAACTGCCCAGGA | Cdk1_As_67225 | |
| ACTTACGGTGTGGTGTATAA | Cdk1_As_67226 | |
| GATGAAGTCCAGTCACCTAC | Cdk17_As_67192 | Cdk17 |
| AGACTGTCCCTCACGCTCAG | Cdk17_As_67193 | |
| GCAGACATCAGGATACCTGA | Cdk17_As_67194 | |
| GGTCCCTATGTAGCACGTTG | Cdk5_As_67160 | Cdk5 |
| GAGTAGACAGATCTCCCGGA | Cdk5_As_67161 | |
| TGAGTAGACAGATCTCCCGG | Cdk5_As_67162 | |
| GGAGTCGGCCGACTTCTACG | Cebpa_As_66924 | Cebpa |
| GATCAAACAAGAGCCCCGCG | Cebpa_As_66925 | |
| GCCGCCTTTGGCTTTCCCCG | Cebpa_As_66926 | |
| ACTTCTACGAGGTGGAGCCG | Cebpa_B_66927 | |
| CATCAGCGCCTACATCGACC | Cebpa_R1.Br_27141 | |
| CAGTTCCAGATCGCGCACTG | Cebpa_R1.Br_27161 | |
| ATATGTGAGAAGTTCAGCCG | Cenpa_1 | Cenpa |
| GCAAACCGCAGACCCCAAGG | Cenpa_2 | |
| TCACATATTTCTCTAACCTG | Cenpa_3 | |
| GGGTCTGTCTTTGCAATCCA | Chaf1a_B_66372 | Chaf1a |
| TCTCCAACCAGCTCTCCTGA | Chaf1a_B_66373 | |

| | | |
|---|---|---|
| AATACTGGAGTCCTGCCCCG | Chaf1a_B_66374 | |
| GCTAAAGGAGGAAAAACGCA | Chaf1a_B_66375 | |
| ATATTTGCTGGGCAACCGAT | Chaf1b_B_66368 | Chaf1b |
| TCCACAGGCTGGCATCCGCG | Chaf1b_B_66369 | |
| GACGGCGGTGTCCACCCCCG | Chaf1b_B_66370 | |
| AGAGGTCATCAAATTCCCAT | Chaf1b_B_66371 | |
| TAGCACTTAATCCGTAGTAG | Chd1_R1.Br_2801 | Chd1 |
| GGAAACGGCCGAAGAAACGT | Chd1_R1.Br_2802 | |
| GTCTCACATCCACAACACAT | Chd1_R1.Br_2804 | |
| GGAGGGAGTAGAGCTCCCGG | Chd1l_As_66320 | Chd1l |
| GCCAATTGACTCCTTCCAGT | Chd1l_As_66321 | |
| CGTGCTGCTGACGACATACG | Chd1l_As_66322 | |
| GAGCTTTGATCAGATAACAG | Chd2_B_66316 | Chd2 |
| AAGCAACCTAAGATTCAGCG | Chd2_B_66317 | |
| CTTATAATACTGTTTCTGAA | Chd2_B_66318 | |
| AGTCAGTCGGAAAGTGAGCA | Chd2_B_66319 | |
| AGGCACTGACACGATCCTGG | Chd3_As_66312 | Chd3 |
| GTGCCAATAACCCCTTCAAA | Chd3_As_66313 | |
| TATGCTTCGGAGACTCAAGG | Chd3_As_66314 | |
| TTATGTGCAGAAAGAAGCAT | Chd4_As_66308 | Chd4 |
| AGAAGAAGAGCAAATCCAAG | Chd4_As_66309 | |
| TCTTAGGTCCCAATGCTCGG | Chd4_As_66310 | |
| CAAACGTTGTGATGACGACG | Chd6_R1.Br_36774 | Chd6 |
| CCGACCCAGACAAATCACCC | Chd6_R1.Br_36776 | |
| TCACTCGAAACTCCTATGAG | Chd6_R1.Br_36777 | |
| ACTGTTTGACGGGAATCCCA | Chd7_B_66296 | Chd7 |
| TCGCAGTCAGAGCAGCAGGT | Chd7_B_66297 | |
| GCACGTAGCACAGATGACCG | Chd7_B_66298 | |
| ATTCTCCGTGGAGAGCAGCG | Chd7_B_66299 | |
| ACCGTTACAAAATACCGTAG | Chd8_R1.Br_31506 | Chd8 |
| TTGTCCTCCAGTATTACTGG | Chd8_R1.Br_31507 | |

| | | |
|---|---|---|
| TGAGAAATACAATACCATGA | Chd8_R1.Br_31509 | |
| AGAAACCATTAGTTTCCGCG | Chd9_R1.Br_49364 | Chd9 |
| CAATTGAATCAGAAGGACGT | Chd9_R1.Br_49365 | |
| GCCAGAGCCAGACCATTCCG | Chd9_R1.Br_49366 | |
| AGGTGTAAATTGGCTCCGAA | Chmp1a_R1.Br_60284 | Chmp1a |
| CTGGATGTGCACACATCGGT | Chmp1a_R1.Br_60285 | |
| GGCCCTTCAGCAGAAAAATG | Chmp1a_R1.Br_60286 | |
| TCAATGGCACAAGCCATGAA | Chmp2a_B_66196 | Chmp2a |
| GTATCTTGAGGGACACAGCT | Chmp2a_B_66197 | |
| CCAGGAAAAGAAAATCATTG | Chmp2a_B_66198 | |
| ACTACTTCGGCAAAACCAGA | Chmp2a_B_66199 | |
| TGGTAATAAGGAAGCGTGCA | Chmp2b_R2.Br_33658 | Chmp2b |
| GGCACCGGCCATCTTCATCT | Chmp2b_R2.Br_33659 | |
| AGAGCTGCGTGGCACCCAGA | Chmp2b_R2.Br_33660 | |
| CATCATGAAGAGCTCTCCCG | Chrac1_As_66128 | Chrac1 |
| GGTGTCGAGCATCAACCAGG | Chrac1_As_66129 | |
| CGAGGTGTCGAGCATCAACC | Chrac1_As_66130 | |
| TTAAAAAGTTCAATTCCTCA | Chuk_As_65952 | Chuk |
| GCATCCACACTACCTTCCGG | Chuk_As_65953 | |
| AGGCCTTTACAACATTCGCA | Chuk_As_65954 | |
| GGCGCTCCGAGAGAAAAGTG | Cir1_R2.Br_29526 | Cir1 |
| TGCCTACCTGAGCCATCAGT | Cir1_R2.Br_29527 | |
| GCTGCTATCTGATGTCTCCG | Cir1_R2.Br_29529 | |
| GTCGCTTCTGAACAGATACG | Cit_As_65852 | Cit |
| TCTTCGCCTGAGAAGCCCGA | Cit_As_65853 | |
| GATGTTGTTGAAGGTCCGGG | Cit_As_65854 | |
| ATGTCCTCATTAAGGAGCTG | Clock_As_65384 | Clock |
| GTATCAACTTCAACACACAA | Clock_As_65385 | |
| ACTAGCATCTGACTGTGCAG | Clock_As_65386 | |
| CATACTCGAGATATCCGATG | Crebbp_R2.Br_34091 | Crebbp |
| TAATGAATCAGGCTCAACAA | Crebbp_R2.Br_3411 | |

| | | |
|---|---|---|
| TGAACCTACTGAATCCAAGG | Crebbp_R2.Br_3412 | |
| ATTAGCCAGCTGCTCTACGA | Cstf1_As_63561 | Cstf1 |
| AAAGAGTTCGGATCACTGGG | Cstf1_As_63562 | |
| GACACGTGACTTCATCCACG | Cstf1_As_63563 | |
| GCGCGCGCGGCCTTACCCAG | Ctbp1_As_63533 | Ctbp1 |
| GAGTGTAGAGCAGATCCGAG | Ctbp1_As_63534 | |
| GCTCACGCGCTGTAGCCCCA | Ctbp1_As_63535 | |
| GAGTGTGGAACAGATCCGTG | Ctbp2_As_63529 | Ctbp2 |
| CCACACCATCACCCTCACCA | Ctbp2_As_63530 | |
| TGGATAAGCACAAAGTCAAG | Ctbp2_As_63531 | |
| GGTAAGGTGTGACATATCAT | Ctcf_As_63517 | Ctcf |
| ATGAAGACTGAAGTCATGGA | Ctcf_As_63518 | |
| GTTCAGATGATTCCTCAGGA | Ctcf_As_63519 | |
| GCAGCAGCAGAAGCAGCAGA | Cul4b_As_63237 | Cul4b |
| CCTGGCTAAATCTTCTACCG | Cul4b_As_63238 | |
| GCAAAAGCTAAAAGAAGCAG | Cul4b_As_63239 | |
| GATCACTGAGAAGATGGCCA | Cxxc1_As_63081 | Cxxc1 |
| CCGACAGTACCATTCCCGGA | Cxxc1_As_63082 | |
| TGTCACGGCAGAAGTCACAG | Cxxc1_As_63083 | |
| AGGAACAGACAAAAGTACCG | Cxxc5_R1.Br_30642 | Cxxc5 |
| CACAGAGATGCTAAAGCGCG | Cxxc5_R1.Br_30643 | |
| GCTCTTGTTCCGACGCTCAG | Cxxc5_R1.Br_30644 | |
| GGACAAGTCCTACCTCTCCG | Ddb1_As_61933 | Ddb1 |
| TGATTTGCCGGAGCTCCTGA | Ddb1_As_61934 | |
| TATTGTGTGCCATAACCGGG | Ddb1_As_61935 | |
| CTCAGGAGCAAAAGTCGCAG | Ddb2_As_61929 | Ddb2 |
| TGTTTCTGCCAAGAGCAGAG | Ddb2_As_61930 | |
| GACGTAGTCCTCCTGTCAAA | Ddb2_As_61931 | |
| GCTTTGATAGGAAATACCTG | Ddx21_As_61829 | Ddx21 |
| ATACCTGTGGAGCAGAAAGA | Ddx21_As_61830 | |
| AACCAAAGAGATAATAACAG | Ddx21_As_61831 | |

| | | |
|---|---|---|
| CCCCATCTAAAGATAGGAAG | Ddx23_As_61825 | Ddx23 |
| AATTCCTCTCTAAAGCAGAG | Ddx23_As_61826 | |
| AGAAGATGCTCGAAGAGGAG | Ddx23_As_61827 | |
| CCAAAGCTACTACCTTCAGG | Ddx27_As_61809 | Ddx27 |
| TCTGATCTACAAGCCCCGCC | Ddx27_As_61810 | |
| TCTCAGTGAAAACAAAGTCA | Ddx27_As_61811 | |
| GGCTTTCTTGGCCAAAACAG | Ddx31_As_61801 | Ddx31 |
| GTGATGCTCCACCAGCAAAA | Ddx31_As_61802 | |
| GTTCCTGGAGTGTTGATGGG | Ddx31_As_61803 | |
| GACAGGCTCAATCTGCTGCA | Ddx39_As_61797 | Ddx39 |
| CCTGGTCATGTGCCACACGA | Ddx39_As_61798 | |
| GGCTTTGAGCATCCTTCAGA | Ddx39_As_61799 | |
| GCTGTCTGAAGGGCTCCAGA | Ddx47_As_61761 | Ddx47 |
| GGAGCCCACGAGCAAGATGG | Ddx47_As_61762 | |
| GCTTCGATCTGGATCTTGGT | Ddx47_As_61763 | |
| CTAACACGTGAACAAAAGGA | Ddx50_As_61749 | Ddx50 |
| AGAAGAAAAGACCTGCCAAA | Ddx50_As_61750 | |
| GTAGCTTTGAAAGATCCAAA | Ddx50_As_61751 | |
| AGGCTCCTAGGGCGAAGCGG | Ddx51_As_61745 | Ddx51 |
| TGGCGCTGTTCCACATAGCG | Ddx51_As_61746 | |
| TCATACTCACCTGTACCACG | Ddx51_As_61747 | |
| GCAGGCCCCCAGCATGTCGG | Dek_As_61488 | Dek |
| GGTTCCTTCTCGGATGAGGG | Dek_As_61489 | |
| GTGAGGTTCTTGATTTAGAG | Dek_As_61490 | |
| GCAGACGCAGAAGAACAAGG | Dhx36_As_61176 | Dhx36 |
| GGTTTGCTCCTGAGGATCAT | Dhx36_As_61177 | |
| CCTGCATCTCTATATATCGA | Dhx36_As_61178 | |
| TGGCATACTGAGTGAGATCG | Dhx8_R2.Br_55104 | Dhx8 |
| TAATTGTAATTGGAGAGACA | Dhx8_R2.Br_55105 | |
| CCCAGGTCGAACCTATCCAG | Dhx8_R2.Br_55106 | |
| CATTAGGTATGGATGATAAA | Dido1_As_61128 | Dido1 |

| | | |
|---|---|---|
| ACTGCTTTCAACGCTCCCCT | Dido1_As_61129 | |
| AAATACAGAGGAAAACCCCA | Dido1_As_61130 | |
| CTAGAACTCGGGGGTCCAGA | Dmap1_As_60920 | Dmap1 |
| GCTTCGTAGTTATTCACGAT | Dmap1_As_60921 | |
| CTGACCTTCAAGAGGCCTGA | Dmap1_As_60922 | |
| TGAAACATCATCCAGACAAA | Dnajc2_As_60640 | Dnajc2 |
| AAGCAAAAGCAGAGGCCAGG | Dnajc2_As_60641 | |
| GCTGCTCCTGCCGAGCGCCG | Dnajc2_As_60642 | |
| ACCTGCCCAACATACAGGTG | Dnmt1_1 | Dnmt1 |
| CATCACGGCTCACTTCACGA | Dnmt1_2 | |
| TAATGTGAACCGGTTCACAG | Dnmt1_3 | |
| TTGTTTGTAGAGCTGCCAAA | Dnmt1_B_60504 | |
| TGAAACTTCACCTAGTTCCG | Dnmt1_B_60505 | |
| CCACCAGGAACAGTTCCAGG | Dnmt1_B_60506 | |
| ACATCCTTACCTCTGTCCCA | Dnmt1_B_60507 | |
| AGTTTACACCGACATGTGGG | Dnmt3A_1 | Dnmt3A |
| GATCATTGATGAGCGCACAA | Dnmt3A_2 | |
| GCGGAGTGAACCCCAACCTG | Dnmt3A_3 | |
| AATGAAGAGTGGGTGCTCCA | Dnmt3a_B_60500 | |
| TTGGGGCTCCAGGATGGCCG | Dnmt3a_B_60501 | |
| GCACTGCAGAAGGAGCTCAG | Dnmt3a_B_60502 | |
| GCTCTCCAATGCCAAAGCCC | Dnmt3a_B_60503 | |
| GCCTCCCCCAGAATCACCCG | Dnmt3b_B_60496 | Dnmt3b |
| TGGTAGCCGGAAACTCCACA | Dnmt3b_B_60497 | |
| CATGAAGTCGACGCTGGCAG | Dnmt3b_B_60498 | |
| CAGCCTTCTGAATTACACGC | Dnmt3b_B_60499 | |
| GAGACTACCAGAATGCTATG | Dnmt3l_As_60492 | Dnmt3l |
| CTAAGCCCCTACTCACCTGA | Dnmt3l_As_60493 | |
| GGTACATGTTCCAGTTCCAC | Dnmt3l_As_60494 | |
| AGAGGACCTCAAAATCACAG | Dnttip2_As_60472 | Dnttip2 |
| AGATGTGTCCCAATTCTCCG | Dnttip2_As_60473 | |

| | | |
|---|---|---|
| CCAAAATTTGCATATCCAAG | Dnttip2_As_60474 | |
| GTTCCCTCCATTCCTCTGAG | Dot1l_As_60360 | Dot1l |
| GTTGATCCTGAAGTTCAGAG | Dot1l_As_60361 | |
| GCAAGTACACTCACGTAGAC | Dot1l_As_60362 | |
| ATAGGAGTATAACTGTCCAG | Dpf2_As_60324 | Dpf2 |
| GCAATACTACAAAGATGCCA | Dpf2_As_60325 | |
| TGGATGGAAAAGCGACACCG | Dpf2_As_60326 | |
| GGCATGACTTACCTGGGCCG | Dpf3_As_60320 | Dpf3 |
| GCAAGAACAGGACCAGAGGA | Dpf3_As_60321 | |
| GCTTGGGGACCAGTTCTACA | Dpf3_As_60322 | |
| CATCTGCTCCGACTCCATGG | Dpy30_As_60212 | Dpy30 |
| GAGTCGGAGCAGATGCTGGA | Dpy30_As_60213 | |
| AAAGTCATCGAAACAGAAGG | Dpy30_As_60214 | |
| TCTTATTTATAGCAGCTCTG | Dr1_As_60180 | Dr1 |
| GCTGGTAGTGAACTGCTGCA | Dr1_As_60181 | |
| ATCATCGTCGTTGCCAGACG | Dr1_As_60182 | |
| GATGATGTGAGGGCAGACAG | Dtx3l_As_59992 | Dtx3l |
| CCGCAGCCGTACGAGTAGCG | Dtx3l_As_59993 | |
| AAATGTCAGCAGGAGAGCCG | Dtx3l_As_59994 | |
| TACAACTCATAGCCAGACTG | Dzip3_As_59706 | Dzip3 |
| CTACTTGTTCTGTCTCCAGC | Dzip3_As_59707 | |
| CCAATCAGTGCTCTTCCCAA | Dzip3_As_59708 | |
| GCTGGGCAGCGGCATGAACG | Ebf1_As_59558 | Ebf1 |
| GGTGCTGGACGCCAACACAG | Ebf1_As_59559 | |
| GATTTCCGCAGGTTAGAAGG | Ebf1_As_59560 | |
| AGTATGAAGGAAGAGCCGCT | Ebf1_B_59561 | |
| CTTTCCTCACTCACCCACCA | Egr1_1 | Egr1 |
| GCTCATCCGAGCGAGAAAAG | Egr1_2 | |
| GTTGTGGAAACAGATAGTCA | Egr1_3 | |
| AAAATCCCAGTAACTCTCAG | Egr2_1 | Egr2 |
| CAGATGAACGGAGTGGCGGG | Egr2_2 | |

| | | |
|---|---|---|
| GTGGTGCGGATTATAAGGGG | Egr2_3 | |
| AGAAGTCCTACTATCCTCGG | Ehmt1_1 | Ehmt1 |
| GGGCCGGTGTACAAACAGCG | Ehmt1_2 | |
| TCGCAGACAAAGGTGCCCAG | Ehmt1_3 | |
| GGATTTCTACCTTTGCAGCC | Ehmt1_B_59110 | |
| CTGCGGCTACTTCTGCACGG | Ehmt1_B_59111 | |
| CTTTGCTTAGTACCTGCCCG | Ehmt1_B_59112 | |
| CCTCACCTCAGCATCAGCGG | Ehmt1_B_59113 | |
| GGTGAGCTACACGAAAGTCG | Ehmt2_1 | Ehmt2 |
| TTCGGGGCCTACCTCGCCCT | Ehmt2_2 | |
| TTGGCAATTAATTACCAGCG | Ehmt2_3 | |
| GCTGCACTCACCTCTCTCGG | Ehmt2_B_59106 | |
| TCTTCACTGGAGACAGAACG | Ehmt2_B_59107 | |
| TGAAGCCATCTAGAAAACGG | Ehmt2_B_59108 | |
| CTGCGGCTACTTCTGCACAG | Ehmt2_B_59109 | |
| CCAGCGCGGTCTGTTCCCCG | Eid1_1 | Eid1 |
| CGACGAGTTTGATGACTGGG | Eid1_2 | |
| GCCCGGCGAGGGCTATATGG | Eid1_3 | |
| CTATATGGAGGTAGGCCGCG | Eid1_As_59098 | |
| GGCTATATGGAGGTAGGCCG | Eid1_As_59099 | |
| GCTCGCTGCTCCCGTCTGCG | Eid2_As_59094 | Eid2 |
| GCGCGACCTGGGGAGACCCG | Eid2_As_59095 | |
| AGCGGCCCGCGAAGGCCCGG | Eid2_As_59096 | |
| CGTCGTCTACGAAATCCTCG | Eif3b_R1.Br_212421 | Eif3b |
| AAAGCAATGATGTTACCACC | Eif3b_R1.Br_212431 | |
| GATGCTGTGAAAAACGCCGA | Eif3b_R1.Br_212441 | |
| TCAGTATGCATATTTCCATG | Eif3d_R1.Br_239021 | Eif3d |
| CGGATGAGATTCGCACAGGT | Eif3d_R1.Br_239031 | |
| GCCGACTTACCGGTAAGCGA | Eif3d_R1.Br_239051 | |
| ACATTGTTGTTGGTACTCCA | Eif4a2_As_58950 | Eif4a2 |
| GCAGATGAAATGTTGAGCCG | Eif4a2_As_58951 | |

| | | |
|---|---|---|
| AGTACCAACAACAATGTGAG | Eif4a2_As_58952 | |
| TTTGATCCTGGCTCCAACCA | Eif4a3_As_58946 | Eif4a3 |
| GTGTCCGTAAACTTCTCCGG | Eif4a3_As_58947 | |
| GCTGGACTACGGACAGCACG | Eif4a3_As_58948 | |
| CCAGGCTTTGACCAGTTAAG | Elp5_As_58714 | Elp5 |
| TTATCAAGAAATCTGCACTT | Elp5_As_58715 | |
| GCTTGACAGAGTGTAACACA | Elp5_As_58716 | |
| GTACTGGAATTCGAAAACAG | Ep300_As_58406 | Ep300 |
| TGATATTGTGAAGAGCCCCA | Ep300_As_58407 | |
| TTACTGGTAGTCATTCCTGT | Ep300_As_58408 | |
| TTAGCTGTTGAGTCTTCACA | Ep400_As_58402 | Ep400 |
| GCAGAGAAAGTAGCAATCGG | Ep400_As_58403 | |
| TGAGCCACTCACAGGTACAA | Ep400_As_58404 | |
| GTAAACAGTCTTGCTCCTGA | Ercc6_As_58130 | Ercc6 |
| GGTCATGTACAACATCCCTG | Ercc6_As_58131 | |
| TTTGTAGAGGATGTTCCACG | Ercc6_As_58132 | |
| AATCAAGTCTGATTGTAAGG | Esco1_As_57982 | Esco1 |
| GTAGTTCTCAGTGCAACCAA | Esco1_As_57983 | |
| GCTGTAGTCTTTGTGATCTG | Esco1_As_57984 | |
| CAACTTGTACTCCAAGAAAG | Esco2_As_57978 | Esco2 |
| TTATTCTTCTTGGTACAGAC | Esco2_As_57979 | |
| TTATGTTTCTTATTAGGTGA | Esco2_As_57980 | |
| GCACTCACCAGCTCTCCCCC | Exosc2_As_57644 | Exosc2 |
| TCATCCATGAACCTACCTGG | Exosc2_As_57645 | |
| GCCGGGGGACACGATCACCA | Exosc2_As_57646 | |
| ATTGTGGGCAGGCATCAAGA | Eya1_As_57592 | Eya1 |
| TTAAAGAGACTGACTCCGAG | Eya1_As_57593 | |
| TCTTATCCCGGCTTTGGCCA | Eya1_As_57594 | |
| CGGATGAGACATCATCCACG | Eya2_R1.Br_5253 | Eya2 |
| TCCAGGCATCGCCAAATCAG | Eya2_R1.Br_5254 | |
| GTTGGGTACGCTGTATAGGG | Eya2_R1.Br_5255 | |

| | | |
|---|---|---|
| AACATGACTGTCAAGAACCG | Eya3_R2.Br_5257 | Eya3 |
| TATAATCATTGGATGAGCGA | Eya3_R2.Br_5258 | |
| CAGGAGAGCTTACCGAAAGG | Eya3_R2.Br_5260 | |
| GCTGTGGGGTAGAAAACCCA | Eya4_As_57580 | Eya4 |
| GCTGGTGTAGAAAGAATGTG | Eya4_As_57581 | |
| CTTATAGTAATAACAAGCAG | Eya4_As_57582 | |
| GCGCTTATATACATTTGGGG | Ezh1_As_57576 | Ezh1 |
| GCTGTCGTCTTGCTTTCCAT | Ezh1_As_57577 | |
| GATCACTACCTTCTTCACCG | Ezh1_As_57578 | |
| AGAGTACATTATAGGCACCG | Ezh2_1 | Ezh2 |
| GACACCACCTAAACGCCCAG | Ezh2_2 | |
| TAGCAAAGATGCCTATCCTG | Ezh2_3 | |
| TTATCAGCAGGAAATTTCCG | Ezh2_B_575721 | |
| ACACGCTTCCGCCAACAAAC | Ezh2_B_575731 | |
| AGAGTACATTATAGGCACCG | Ezh2_B_575741 | |
| TGAGACTGAGACAGCTCAAG | Ezh2_B_57575 | |
| TCTTCTTTCTCCTTTCCAGA | Fam208a_As_56880 | Fam208a |
| GCAAACCTCCGAAGCGGCCG | Fam208a_As_56881 | |
| GTTAAACCTTTAGGAGACCG | Fam208a_As_56882 | |
| CTGCAGTGACGAGCGCCGCG | Fbxw9_As_55973 | Fbxw9 |
| GACTGCAGTGACGAGCGCCG | Fbxw9_As_55974 | |
| CCATGGATCTTTCCTCAGGG | Fbxw9_As_55975 | |
| ATGGTGAAGACCGTGTCAGG | Fos_1 | Fos |
| CAGACCTCCAGTCAAATCCA | Fos_2 | |
| TGCTGGGGCTTACGCCAGAG | Fos_3 | |
| AGGTACTGAGACTCGGCGGA | Fosb_1 | Fosb |
| CAGCTAAGTGCAGGAACCGT | Fosb_2 | |
| GCTGACAGATCGACTTCAGG | Fosb_3 | |
| CGTGTGCAAAATCAGCCCCG | Fosl2_1 | Fosl2 |
| TGGATAGGGATTGGACATGG | Fosl2_2 | |
| TGGATAGGGATTGGACATGG | Fosl2_3 | |

| | | |
|---|---|---|
| CTAAGAGCAGTGAGAAGCAG | Foxp2_As_55013 | Foxp2 |
| CTTCAGTTCTGAATGCAAGG | Foxp2_As_55014 | |
| ATGGCCATAGAGAGTGTGCG | Foxp2_As_55015 | |
| CTGCAGAGCAGAAGACCGAA | Gadd45a_As_54393 | Gadd45a |
| CGTTCTCGCAGCAGAACGCA | Gadd45a_As_54394 | |
| CATTACGGTCGGCGTGTACG | Gadd45a_As_54395 | |
| GCTGGTGGCGAGCGACAACG | Gadd45b_As_54389 | Gadd45b |
| GGCCGCCTCGTACACCCCCA | Gadd45b_As_54390 | |
| GAGGCGATCCTGACGCTGCG | Gadd45b_As_54391 | |
| ATGACTCTGGAAGAAGTCCG | Gadd45g_As_54385 | Gadd45g |
| GGCGGACTCGTAGACGCCAG | Gadd45g_As_54386 | |
| GTCGCCCTCATCTTCTTCAT | Gadd45g_As_54387 | |
| AGTATGGAGGGAATTCCTGG | Gata1_B_541411 | Gata1 |
| CCCTCCATACTGTTGAGCAG | Gata1_B_541421 | |
| CCACTGCTCAACAGTATGGA | Gata1_B_541431 | |
| TGGACACCTTGAAGACGGAG | Gata1_B_54144 | |
| GCAGGGGGTAGTGTAGCCCG | Gata2_B_541371 | Gata2 |
| GAGAGCATGAAGATGGAAGG | Gata2_B_541381 | |
| GCCTCTGTCTGTTTACCCAG | Gata2_B_541391 | |
| GCTGCGCATTCAATACGGCG | Gata2_B_54140 | |
| CCTGGGCTGTGCAACAAGTG | Gata2_R1.Br_61771 | |
| GGCCGGGAGTGTGTCAACTG | Gata2_R1.Br_61781 | |
| ACAGCTGCTGCCTCCCGACG | Gata2_R1.Br_61791 | |
| CCGTACTATGCTCAGATCAG | Gatad1_R1.Br_30230 | Gatad1 |
| AGACTACGTCGTCCTCCATG | Gatad1_R1.Br_30231 | |
| TGCAGAGTCAATCTTCTACA | Gatad1_R1.Br_30233 | |
| TCAGACAGCATTCCTCTACG | Gatad2a_R1.Br_60072 | Gatad2a |
| GGGCGCACGAACCTGAAGTG | Gatad2a_R1.Br_60073 | |
| AAATTCCACTCCCACCAGTG | Gatad2a_R1.Br_60075 | |
| TTGCCCAGTGAGCCAGACCG | Gatad2b_B_54109 | Gatad2b |
| GCAAAACGACTGAAAATGGA | Gatad2b_B_54110 | |

| | | |
|---|---|---|
| GCAGCTTATCAAGCAACTGA | Gatad2b_B_54111 | |
| TGGATATGAGTGCTAGACGG | Gatad2b_B_54112 | |
| GTTGAGTGTGGTGGTCACTG | Gltscr1_As_53229 | Gltscr1 |
| GCATTCCCAGGACTGCCATT | Gltscr1_As_53230 | |
| CTTTCAGTGGAGCAGCTGCA | Gltscr1_As_53231 | |
| CGACGATCTGAGGAGAGGCG | Gltscr1l_As_53225 | Gltscr1l |
| GCACGGACCTAGCAGTAAAT | Gltscr1l_As_53226 | |
| GCGATGATGTGACGAACGCA | Gltscr1l_As_53227 | |
| GGCTACAGGAGCGCACGACC | Gltscr2_R2.Br_32262 | Gltscr2 |
| AGGGCAAGGTCAACCCGTAA | Gltscr2_R2.Br_32263 | |
| GGCCCCAGAAACAAGAAGCG | Gltscr2_R2.Br_32264 | |
| AGGTGAATATCCACAAACTG | Gon4l_R2.Br_42362 | Gon4l |
| TCTACATGCCCAGCCCATGG | Gon4l_R2.Br_42363 | |
| GGTTACCAGAGAAGACCGAG | Gon4l_R2.Br_42364 | |
| AGTGCCTGCCCCACACCACA | Gse1_As_50576 | Gse1 |
| CAGAGACACTGGGTTCCAGA | Gse1_As_50577 | |
| AGTCCTGTACAACTGAGCCG | Gse1_As_50578 | |
| TGAAATCCCTCCAGGAAGTG | Gtf2a1_R2.Br_45324 | Gtf2a1 |
| CTCATTCCTGCATCCCAGCA | Gtf2a1_R2.Br_45325 | |
| AGTGGATGAACAAGTCCTGA | Gtf2a1_R2.Br_45326 | |
| TCAGCCGGCTCTTATACCCG | Gtf2h3_R2.Br_52752 | Gtf2h3 |
| CCAGCACGCAGGCGTCGATG | Gtf2h3_R2.Br_52753 | |
| TGCAAACGAGGTGATCGCTG | Gtf2h3_R2.Br_52754 | |
| AAACAAGCCCGAACTCACGC | Gtf2h4_R2.Br_7093 | Gtf2h4 |
| GGGCTCAGGAGGAAAGTACG | Gtf2h4_R2.Br_7094 | |
| AAGCCAGCGGAAGTGATGCA | Gtf2h4_R2.Br_7096 | |
| CATCGCGGATGCGTTCGTCG | Gtf3a_R2.Br_28730 | Gtf3a |
| GCATGTCCTGATTCACACCG | Gtf3a_R2.Br_28731 | |
| GCACCTATGCAAACACACGG | Gtf3a_R2.Br_28732 | |
| TCCGGACGCGGCAAGACCGG | H2afx_As_49968 | H2afx |
| TCTACCTCGTACACTATGTC | H2afx_As_49969 | |

| | | |
|---|---|---|
| TCGTACACTATGTCCGGACG | H2afx_As_49970 | |
| GCTGTCGCCATGTCGAGCCG | H2afy_As_49964 | H2afy |
| GCAGCAAGAGACAACAAGAA | H2afy_As_49965 | |
| ACAGCTAACAGGATGTGCCG | H2afy_As_49966 | |
| GCGTCATCTTTCCAGTAGGC | H2afy2_As_49960 | H2afy2 |
| CGAGGGACAACAAGAAGGCA | H2afy2_As_49961 | |
| GCTGGCCGTTGCCAACGACG | H2afy2_As_49962 | |
| AGAGTGCGCCCTCTACTGGA | H3f3a_As_49948 | H3f3a |
| ATAGCGTCTGATTTCACGGA | H3f3a_As_49949 | |
| GATAGCGTCTGATTTCACGG | H3f3a_As_49950 | |
| GGTGGAGTATAAGAGTGCCG | Hat1_B_49817 | Hat1 |
| AGCCTTGGAGAAATTTCTGG | Hat1_B_49818 | |
| GTGTTTGTGCAAAACCCAGG | Hat1_B_49819 | |
| ACAGCAATCGAGCTGAAACT | Hat1_B_49820 | |
| ACAGCAGTATGTGACTCCCG | Hcfc1_As_49709 | Hcfc1 |
| GCAGGTCTCATGGTAACCAG | Hcfc1_As_49710 | |
| ACCCCAAGAACAACATTCCG | Hcfc1_As_49711 | |
| AGAAGAAGAGAAAACCAAGG | Hdac1_1 | Hdac1 |
| CTTACCGACAGAGCCTCCCG | Hdac1_2 | |
| TAAAGGGAGTTCTCACCCGT | Hdac1_3 | |
| GGATTCGGTGAGGCTTCATG | Hdac1_As_49657 | |
| CCTCTTCCACGCCATCGCCA | Hdac1_As_49658 | |
| TGATCCTAGGTACCACCAGA | Hdac1_As_49659 | |
| AGAAGCTTCCATGCTCATAG | Hdac10_1 | Hdac10 |
| CATATCAGGCTGGAAACCAT | Hdac10_2 | |
| TGTACTGCTTAGACAGTGCG | Hdac10_3 | |
| AGGTGGTTCTCACCTCACCA | Hdac10_B_49653 | |
| GGTCTGGGTCTTCTGCACCA | Hdac10_B_49654 | |
| ACTGTGCTAGGGACTCCAAG | Hdac10_B_49655 | |
| GTGGAGCTCCTCTTTATCCA | Hdac10_B_49656 | |
| ATGAAGGTGATGTTGTAACG | Hdac11_R1.Br_59220 | Hdac11 |

| | | |
|---|---|---|
| GCCACCACAAAGGACCACTG | Hdac11_R1.Br_59221 | |
| GGCATCGAGATCAATGATGG | Hdac11_R1.Br_59223 | |
| ACTTGATATACTCATCGCTG | Hdac2_1 | Hdac2 |
| ATCTTCTCCGACGTTAACTG | Hdac2_3 | |
| ATTGACATAGACATCCACCA | Hdac2_As_49645 | |
| TCACCTTCCACAGTTAACGT | Hdac2_As_49646 | |
| TGAGTCATCCGGATTCTATG | Hdac2_As_49647 | |
| CAATGTAGAGCACCCGAGGG | Hdac3_1 | Hdac3 |
| GACGTATGACAGGACTGACG | Hdac3_2 | |
| GCACATTGAGACAATAGTAG | Hdac3_3 | |
| TATCAATGTAGAGCACCCGA | Hdac3_As_49641 | |
| GGGGTCGTAGAAATACGCCA | Hdac3_As_49642 | |
| ATTGATATCGACATCCACCA | Hdac3_As_49643 | |
| TCACTTACCCATACCAGTAG | Hdac4_1 | Hdac4 |
| TGTGGCTGCCAAATGACACA | Hdac4_2 | |
| TTTGACGCCTACAGACAGCG | Hdac4_3 | |
| TGACGTGTAGAGAGGAAGTG | Hdac4_B_49637 | |
| GCAGCCTGAGAGCCACCCAG | Hdac4_B_49638 | |
| GCAGTGGTTCAGGTTCCGGT | Hdac4_B_49639 | |
| GACATACCTGTAACATCCAG | Hdac4_B_49640 | |
| CAGCTCATGACACTGGCTGG | Hdac5_1 | Hdac5 |
| CGTGCCCTGTACTTACGGTG | Hdac5_R1.Br_7437 | |
| GCCGGGCGCACTGTTACACA | Hdac5_R1.Br_7438 | |
| CTTTCTGTAGAGCCTTCCCG | Hdac5_R1.Br_7439 | |
| ATGATGCGTAAGATGCGCTG | Hdac6_R1.Br_7441 | Hdac6 |
| TCCCTGCACCGGTATGACCG | Hdac6_R1.Br_7442 | |
| CCTGAGACAAGAGTGCCAGT | Hdac6_R1.Br_7444 | |
| ACGTTTGTGATGCTACCCTG | Hdac7_1 | Hdac7 |
| AGAGGGTGGACATGACCTCA | Hdac7_2 | |
| GCTACCTGTAGGGAATACTG | Hdac7_B_49625 | |
| GTATCAGCACGGAGTCCCCG | Hdac7_B_49626 | |

| | | |
|---|---|---|
| GCTCTCTCAGGGAAGCCCCG | Hdac7_B_49627 | |
| GCTCCCCTCTTCCTGCACCA | Hdac7_B_49628 | |
| CCGTTTGGGGACCTTCACAA | Hdac8_R2.Br_35586 | Hdac8 |
| GTTGCAGATAGGCATCAGTG | Hdac8_R2.Br_35587 | |
| GACTCCATAGAATATGGACT | Hdac8_R2.Br_35588 | |
| CGGTCAGTAGCAGCTCTCCA | Hdac9_B_49617 | Hdac9 |
| CTATCTTTGCCTCTGAGAGG | Hdac9_B_49618 | |
| GGTCCCAGTTCACCAAACAA | Hdac9_B_49619 | |
| CCATCCTTCCGCCTGAGTAA | Hdac9_B_49620 | |
| AGTTTGGCAAGCCCAACAAG | Hdgf_As_49601 | Hdgf |
| CTTGTACTCTTTCTGCCGGT | Hdgf_As_49602 | |
| CCGCCATGTCGCGATCCAAC | Hdgf_As_49603 | |
| TATTCTTTACTAATTTCCGA | Hells_As_49505 | Hells |
| ATTGCATTGATGATTCAGAG | Hells_As_49506 | |
| GCTGTATCATGGAACCCGGG | Hells_As_49507 | |
| ATGACAAACTGATTATGGTG | Hira_R2.Br_7605 | Hira |
| TAGCTGCTGACCACTAGGAG | Hira_R2.Br_7607 | |
| AGTTCATGGATGACAACCAG | Hira_R2.Br_7608 | |
| TGGAGGCTTGCGAGACAGCG | Hirip3_R2.Br_59900 | Hirip3 |
| GGATGACAGTAACAGCACAC | Hirip3_R2.Br_59901 | |
| CACAGGTGCTAACTGCCAGG | Hirip3_R2.Br_59902 | |
| GATGAGACCATTAAACGCAG | Hltf_As_48974 | Hltf |
| CCATGTCCTATACGTTCACG | Hltf_As_48975 | |
| ACTGGGTTCTTCACCACACG | Hltf_As_48976 | |
| GGAACCTTCCGGCTTCGACG | Hmbox1_R2.Br_55812 | Hmbox1 |
| AGGGTCCTCTATTGGAATGG | Hmbox1_R2.Br_55813 | |
| ATTGGTAGTGCAAGGCAGTG | Hmbox1_R2.Br_55815 | |
| GCTTCGAGCAAAGAGACCAG | Hmg20a_B_48950 | Hmg20a |
| CTCAGGTTAACACACCCCGA | Hmg20a_B_48951 | |
| GGAGCAGTATCAGAAGACCG | Hmg20a_B_48952 | |
| TCTTCATCTGCAAACAGGGG | Hmg20a_B_48953 | |

| | | |
|---|---|---|
| CTGCTTGACAGCCACTACAA | Hmg20b_R2.Br_7657 | Hmg20b |
| GAGTTTCTGGACCAAAACAA | Hmg20b_R2.Br_7658 | |
| AGCCATCGCAGTCCACACCC | Hmg20b_R2.Br_7660 | |
| AAGGATGGGACTGAGAAGCG | Hmga1_R2.Br_7677 | Hmga1 |
| GCCTCCGGTGAGTCCTGGGA | Hmga1_R2.Br_7678 | |
| CCAACTCCGAAGAGACCTCG | Hmga1_R2.Br_7679 | |
| TCCTCGCTTCTGTGGCACCG | Hmga2_R2.Br_7681 | Hmga2 |
| CAGCCGTCCACATCAGCCCA | Hmga2_R2.Br_7682 | |
| CCTTCTGGGCTGCTTTAGAG | Hmga2_R2.Br_7683 | |
| GAAGTGCTCAGAGAGGTGGA | Hmgb1_R2.Br_7645 | Hmgb1 |
| AACCTACATCCCCCCCAAAG | Hmgb1_R2.Br_7646 | |
| CTTTGTGCAAACTTGCCGGG | Hmgb1_R2.Br_7647 | |
| GAAATGCTCCGAGAGATGGA | Hmgb2_R2.Br_46308 | Hmgb2 |
| GTGGTCTCTTCGGAGCATTG | Hmgb2_R2.Br_46309 | |
| CTGCACGAAGAAGGCGTACG | Hmgb2_R2.Br_46310 | |
| GGAGAACTCAGCAAAATTGA | Hmgb3_B_48923 | Hmgb3 |
| GGTGACCCCAAGAAACCAAA | Hmgb3_B_48924 | |
| CTTCTTTGTGCAGACATGCA | Hmgb3_B_48925 | |
| ACATAAGAAGAAAAACCCAG | Hmgb3_B_48926 | |
| CGCGAAGCCGAAAAAGGCCG | Hmgn1_B_48895 | Hmgn1 |
| GGATGGAGCCGCCAAGGCGG | Hmgn1_B_48896 | |
| TTCGAAACCTACCTTTCCCG | Hmgn1_B_48897 | |
| ACTTACGGCCGACAGCCTCG | Hmgn1_B_48898 | |
| CAAAACCAAGGTGAAGGACG | Hmgn2_As_48891 | Hmgn2 |
| ACTCACCTCGTCCTTCACCT | Hmgn2_As_48892 | |
| GCCAGAGCCCAAACCTAAAA | Hmgn2_As_48893 | |
| AGATTAGCAGAGGTGCTAAG | Hmgn3_R2.Br_46256 | Hmgn3 |
| AACCAAGCTAACTAAGCAGG | Hmgn3_R2.Br_46257 | |
| ACCAAGAAAAACATCAGCTA | Hmgn3_R2.Br_46259 | |
| AGAAGTAAAAGAACAAAACA | Hmgn5_R2.Br_22150 | Hmgn5 |
| AGCCAAAAGTGAAGATGCAG | Hmgn5_R2.Br_22152 | |

| | | |
|---|---|---|
| AGAGCACAAAGATACAGGTG | Hmgn5_R2.Br_22153 | |
| ATGATGATGTATTCCGATCG | Hmgxb3_R2.Br_48428 | Hmgxb3 |
| ACTACAACATACACCCGTCG | Hmgxb3_R2.Br_48429 | |
| ACATACTGCCATCGGCTGAG | Hmgxb3_R2.Br_48431 | |
| TCCATAGTAGTAATCGTCAG | Hmgxb4_R2.Br_36202 | Hmgxb4 |
| GTGCCAATCTCGACATCACA | Hmgxb4_R2.Br_36203 | |
| GAAACTCATTCTCTCACCAA | Hmgxb4_R2.Br_36205 | |
| AGCCGAGGCGGGTCTCACGG | Hopx_1 | Hopx |
| CCTGGAGTACAACTTCAACA | Hopx_2 | |
| TTAAGCAGCGCCTGGCAGAG | Hopx_3 | |
| CATCACCTCAGGAACGCCCA | Ing1_R2.Br_20030 | Ing1 |
| CGAATAACAAGCGGTCCAGG | Ing1_R2.Br_20031 | |
| GAGATCGACGCCAAATACCA | Ing1_R2.Br_20033 | |
| TTGTCATGTGACATAAGTCA | Ing2_R2.Br_34174 | Ing2 |
| CCGCGCTCCTGACCGGAGAG | Ing2_R2.Br_34175 | |
| CGCTGCCCCACGACATGCAG | Ing2_R2.Br_34176 | |
| GCTTCCAAGGAGAACACGCT | Ing3_R2.Br_37254 | Ing3 |
| CAAAGAAGAATAAACCCGAG | Ing3_R2.Br_37255 | |
| TGAGCGTGGTGATTGTTCAC | Ing3_R2.Br_37256 | |
| AGGTAGACAAACACATTCGG | Ing4_R2.Br_21286 | Ing4 |
| TATGACAGCTCTTCTAGCAA | Ing4_R2.Br_21287 | |
| GAGAAACTTTCAGCTCATGA | Ing4_R2.Br_21289 | |
| TGACGAGAGAGTCTTCACTG | Ing5_R2.Br_27911 | Ing5 |
| CTAGACCAGAGAACAGAAGG | Ing5_R2.Br_27912 | |
| AAGTTCCTCTGAAGTTCACA | Ing5_R2.Br_27913 | |
| TGTTTCTTCTTGGTGCCAGG | Ino80_As_46860 | Ino80 |
| GGATGTCAAGTACTTCCAGC | Ino80_As_46861 | |
| GTTGGATGAGGAAATGCGGG | Ino80_As_46862 | |
| GATGGTGGTGGATAATGAAG | Ino80b_As_46856 | Ino80b |
| CTTTCACTGTGATTCCTGAG | Ino80b_As_46857 | |
| ACTTTCACTGTGATTCCTGA | Ino80b_As_46858 | |

| | | |
|---|---|---|
| GCTGTTCCTAGCTACGGCGG | Ino80c_As_46852 | Ino80c |
| GCTGACTGCGGGCAGAATGG | Ino80c_As_46853 | |
| AGCCCTTCTCACAACAGCAG | Ino80c_As_46854 | |
| AAACCTCCAGCACCACCACA | Ino80d_As_46848 | Ino80d |
| ATTGGGCATCTTCAAGGCCA | Ino80d_As_46849 | |
| CTTCCTCGTTGAAAGCAAAG | Ino80d_As_46850 | |
| ACTGTTATCTGAAGATGCGG | Ino80e_As_46844 | Ino80e |
| GGCTCCTGCTTACCTCTTAG | Ino80e_As_46845 | |
| GGTAGTCAGAAGGGAATGGG | Ino80e_As_46846 | |
| GCTTACCTCAGGGATCCACA | Jade1_As_46076 | Jade1 |
| GGGATCCTCAAGGTGCCAGA | Jade1_As_46077 | |
| TCAATGATATGGATGCTGCG | Jade1_As_46078 | |
| GATAGCCCCACACTTGGTGA | Jade2_As_46072 | Jade2 |
| ACACACCTCAGGAATCCACA | Jade2_As_46073 | |
| GGATGACTACTACATCCTGG | Jade2_As_46074 | |
| GGTCTCCAGACAGTGAAGAA | Jade3_As_46068 | Jade3 |
| TGTAGATGCTGCTTGCTCCA | Jade3_As_46069 | |
| GGCATTCTCAAGATTCCAGA | Jade3_As_46070 | |
| CATTCCTAGGTGTGACACCG | Jarid2_R1.Br_8949 | Jarid2 |
| CTGAGCAGACCGATGCGAAG | Jarid2_R1.Br_8950 | |
| TGCCCGAGCGAAGTTTGGAA | Jarid2_R1.Br_8951 | |
| AATACAAGGAGGAACATCGT | Jmjd1c_R1.Br_49164 | Jmjd1c |
| CATGAGCACTCTCTAGACGT | Jmjd1c_R1.Br_49165 | |
| TGTGGAAAACCAACGCACAG | Jmjd1c_R1.Br_49166 | |
| GAGGTCCCGGTAGTACCGCT | Jmjd4_As_46004 | Jmjd4 |
| AGACACGCACCTTTGCCGAG | Jmjd4_As_46005 | |
| GGTGGCGGCCCCACACCCAG | Jmjd4_As_46006 | |
| GCTGCTATCGAAGATGTAAA | Jmjd6_As_46000 | Jmjd6 |
| GCGAGGCCAAGCGAAGTGCG | Jmjd6_As_46001 | |
| TGAATCCCAGTTCCAGAACG | Jmjd6_As_46002 | |
| GCAAGGACTCACGGACAACT | Jmjd8_As_45992 | Jmjd8 |

| | | |
|---|---|---|
| GTGTAGCTCACCTTTCTGGT | Jmjd8_As_45993 | |
| TTTCTGGTAGGAGTAGGTGT | Jmjd8_As_45994 | |
| CTTAGGGTTACTGTAGCCGT | Jun_1 | Jun |
| GAAGGTCCGAGTTCTTGGCG | Jun_2 | |
| GCTCTCGGACTGGAGGAACG | Jun_3 | |
| GGGTAAAAGTACTGTCCCGG | Junb_1 | Junb |
| GGGTAAAAGTACTGTCCCGG | Junb_2 | |
| TGAGCTCCCAGTCCCGACGG | Junb_3 | |
| CACGCAAGAACGCATCAAGG | Jund_1 | Jund |
| CACGCAAGAACGCATCAAGG | Jund_2 | |
| TATGGCGAGGAGGCGCTGAG | Jund_3 | |
| ATATTGAGATCCGTGCCCGA | Kansl1_As_45908 | Kansl1 |
| GCTTCTGTAACCTTCGGGCA | Kansl1_As_45909 | |
| TCACAGACAGTAGTTCCGGA | Kansl1_As_45910 | |
| GTATCCGGCTGGCTTCACTG | Kansl2_As_45900 | Kansl2 |
| GCCGCGGCTGAGAGCACGGG | Kansl2_As_45901 | |
| AACTCCTGCCCCTCCAGCCG | Kansl2_As_45902 | |
| GGTCCATCCACAGTAAGCAG | Kansl3_As_45896 | Kansl3 |
| ATCATAGCTATGCAAAACCG | Kansl3_As_45897 | |
| ACTTCCAGACTTCAGCTCGG | Kansl3_As_45898 | |
| TGGTGCCTGCCTTACCCAAA | Kat2a_B_45884 | Kat2a |
| GCTACTGGAAAAATTTCGGG | Kat2a_B_45885 | |
| GCTTGAGGAAGAGATCTACG | Kat2a_B_45886 | |
| CCGCTGCCGGAATTGAGCAG | Kat2a_B_45887 | |
| TCCCGTGGAATTGATCAATG | Kat2b_R1.Br_12574 | Kat2b |
| GTACCTGTTTGGATTCCTAG | Kat2b_R1.Br_12575 | |
| GAGGCAGACAACGATCGAGC | Kat2b_R1.Br_12576 | |
| GCCACGACGACATTGTCACC | Kat5_R1.Br_45092 | Kat5 |
| CCAAGACCCCTCATACCAAG | Kat5_R1.Br_45093 | |
| ATGAATGGGTGACTCACGAG | Kat5_R1.Br_45094 | |
| ATTGGTGGCGAGTTTGACCG | Kat6a_R2.Br_63088 | Kat6a |

| | | |
|---|---|---|
| CAAAACACTCTATTATGACG | Kat6a_R2.Br_63089 | |
| GCTTCCACACGAGAAAGACA | Kat6a_R2.Br_63091 | |
| TTGCGTAGATCATTACACGG | Kat6b_R2.Br_23334 | Kat6b |
| TCCCGAACTCTATTACTGAG | Kat6b_R2.Br_23335 | |
| CCTCCCGTTCCATCCCCACG | Kat6b_R2.Br_23337 | |
| CCTATCCTGAGGAATATGCG | Kat7_R2.Br_55052 | Kat7 |
| GACTCGGGCAGATCGGCGCG | Kat7_R2.Br_55053 | |
| GCGATAAATCTCATCACCAG | Kat7_R2.Br_55054 | |
| TGGCGAACCAGAAGTCACCG | Kat8_As_45860 | Kat8 |
| GTACGTTTCTCCGATCTCCA | Kat8_As_45861 | |
| GGGAAAATGCAGCAGTCGAG | Kat8_As_45862 | |
| GGAATAGCCGAGACCCCGGA | Kdm1a_As_45308 | Kdm1a |
| GGCCATTACAACTCACCAGA | Kdm1a_As_45309 | |
| GCTGGTCCGTCGGCCCTCCG | Kdm1a_As_45310 | |
| CAACACAGGCGTTCTCACGG | Kdm1b_R1.Br_55524 | Kdm1b |
| AAAATGGTTACACCTCCCGA | Kdm1b_R1.Br_55525 | |
| GGGGAAAGAACAATGATCGG | Kdm1b_R1.Br_55526 | |
| ACAGCATTGAAGATCGAACA | Kdm2a_R1.Br_56736 | Kdm2a |
| TCTACCCGGGGATCCCACAA | Kdm2a_R1.Br_56737 | |
| AAGTACCTCCAAAATCCACA | Kdm2a_R1.Br_56738 | |
| GGGGGACGTTTGAAGCGACG | Kdm2b_R2.Br_21670 | Kdm2b |
| CCATGCGGTTTATACGCCTG | Kdm2b_R2.Br_21671 | |
| GCGCTACTACGAGACACCAG | Kdm2b_R2.Br_21672 | |
| TTACCTGTATAGGTTTCAAA | Kdm3a_B_45292 | Kdm3a |
| GATAAGCCAGAAGTGAAAGC | Kdm3a_B_45293 | |
| GCTTGGACCCATCTACTCAG | Kdm3a_B_45294 | |
| TTACTTTCCTAGCCCCAGCC | Kdm3a_B_45295 | |
| TGTGAAGGAATCCCTGCTCG | Kdm3b_R2.Br_68904 | Kdm3b |
| GGTGCCAGTTGAATACCTTG | Kdm3b_R2.Br_68906 | |
| GACATGGTACACGCTGCCCG | Kdm3b_R2.Br_68907 | |
| GATCTTGCGGAACTCACGAA | Kdm4a_R1.Br_58440 | Kdm4a |

| | | |
|---|---|---|
| CTTCAAATTTGACTTCAGTG | Kdm4a_R1.Br_58441 | |
| AAGCACGCTCACCTGATGGT | Kdm4a_R1.Br_58442 | |
| AGCCCGCATGGTAACCATAG | Kdm4b_R1.Br_51972 | Kdm4b |
| AGCCGAGAGGAAGTTCAATG | Kdm4b_R1.Br_51973 | |
| ATGTCATCATACGTCTGCCG | Kdm4b_R1.Br_51974 | |
| GTGCACTTGTCGGAATGACA | Kdm4c_R1.Br_43132 | Kdm4c |
| CTGGCCGGAGGCTTACCAAG | Kdm4c_R1.Br_43133 | |
| TTTGGATACCAGGATACAAG | Kdm4c_R1.Br_43134 | |
| ATGCGCCCGATAAAACTCAG | Kdm5a_As_45268 | Kdm5a |
| TATCACTCTGGATTTAACCA | Kdm5a_As_45269 | |
| GCTGGACTCTAGGAGTGAAA | Kdm5a_As_45270 | |
| ATAAACTTCATTTCACCCCG | Kdm5b_As_45264 | Kdm5b |
| CTAAGATCTTTCTCTCCACG | Kdm5b_As_45265 | |
| ATTCAGCCTCTGGATCCGCG | Kdm5b_As_45266 | |
| AGACTTACCGAAGGTACTGG | Kdm5c_R1.Br_16150 | Kdm5c |
| GGATGAGGTAAAACGCACGC | Kdm5c_R1.Br_16151 | |
| AGTTCAACAGTTATGGACGA | Kdm5c_R1.Br_16153 | |
| GATTTACTCCTCGAATTCAG | Kdm5d_As_45256 | Kdm5d |
| CCTTCACTACAATAGTTGCA | Kdm5d_As_45257 | |
| TCAACTTCTACAGCAAAAGG | Kdm5d_As_45258 | |
| TTCCTCATCACCGAAAGCGG | Kdm6a_As_45252 | Kdm6a |
| TGTGCTGTACAATTGGACCA | Kdm6a_As_45253 | |
| TCAGCATTGGACAAAGTGCA | Kdm6a_As_45254 | |
| GAGTGCCCACAGAAAAGCGG | Kdm6b_B_45248 | Kdm6b |
| CATTTCAGCTGACTAAGCCA | Kdm6b_B_45249 | |
| TCTCTAAGGTCACTTCCGGG | Kdm6b_B_45250 | |
| TTAGAGAGAGCAGAGTTCAG | Kdm6b_B_45251 | |
| CTACTCTTACCCAAAGGACA | Kdm7a_B_45244 | Kdm7a |
| AATACCCTACTCTTACCCAA | Kdm7a_B_45245 | |
| AAGAATGTCTGAATTGGTGG | Kdm7a_B_45246 | |
| CATCGCACTCGATCATGAAG | Kdm7a_B_45247 | |

| | | |
|---|---|---|
| TCAGGTGCTGATCATGTCAG | Kdm8_B_45240 | Kdm8 |
| TCCTGTCCACCTTCTCACCG | Kdm8_B_45241 | |
| CGTCCTGAGCTCCTTCCAGA | Kdm8_B_45242 | |
| AGTACTGCAGAGGTGGGCAG | Kdm8_B_45243 | |
| AGAAAGGGCGGCGATCAAGG | Kmt2a_1 | Kmt2a |
| CCATGCGTAAGATCTACCGG | Kmt2a_2 | |
| GCAGCCGTTAGACCTCGAAG | Kmt2a_3 | |
| AAGAGCATAGAAAAGAAGAG | Kmt2a_As_44475 | |
| GGATCATCAAGACTCCCCGG | Kmt2a_As_44476 | |
| GTAATGATAGGAGAAGCAGA | Kmt2a_As_44477 | |
| GCTGGTGCGCGAGTTCAGCG | Kmt2b_As_44471 | Kmt2b |
| TCTTCATCCATAAATCGCCG | Kmt2b_As_44472 | |
| CCGCCCCCGCCGGAACCCCG | Kmt2b_As_44473 | |
| AAAAGGCCCATTACCCAATG | Kmt2c_1 | Kmt2c |
| AAGCATTACCTGAATCCATG | Kmt2c_2 | |
| TTATCATCAGCTCCAACCGG | Kmt2c_3 | |
| GGATCAATAGTGTTAGGCCG | Kmt2c_As_44467 | |
| ACTTTGTTCTGAAATCACAG | Kmt2c_As_44468 | |
| CCTTCATGTGACATGCTGCA | Kmt2c_As_44469 | |
| AAATGGCTGTTGATCCCATG | Kmt2d_1 | Kmt2d |
| GGACCTGGAGAAGCACACGA | Kmt2d_2 | |
| GTTCACCATTAATACCCCCA | Kmt2d_3 | |
| TGATGGGTGAAAGTTCCCCG | Kmt2d_As_44463 | |
| TGTGTTAGAGAAGAGAACCA | Kmt2d_As_44464 | |
| GCCTGGTTCTACAAGCACAG | Kmt2d_As_44465 | |
| TCTGTGTCAGACCTTCAACT | Kmt2e_2 | Kmt2e |
| TGGGCTGACAACGTTCACAT | Kmt2e_3 | |
| AGTTCTGTTCACTTTGGCAG | Kmt2e_As_44459 | |
| CAGCATTTGCCACTCCCCCA | Kmt2e_As_44460 | |
| GGATCTGTAGTTATGCGCAG | Kmt2e_As_44461 | |
| AAGAGCAAAGACACCAGGAG | Kmt5a_1 | Kmt5a |

| | | |
|---|---|---|
| GATTTCGGCCTATAAAAGGA | Kmt5b_2 | Kmt5b |
| AGGAGTGCCCAGCTGTGGCT | Kmt5c_3 | Kmt5c |
| TGCAGATTTCTCGGACCCCG | Kti12_R1.Br_46796 | Kti12 |
| TGAACAAGGGGCGATCCCAG | Kti12_R1.Br_46797 | |
| CGAGGCAGTACAGCTCGTAG | Kti12_R1.Br_46799 | |
| TGATGGCACACCACCCAATG | L3mbtl2_R1.Br_54140 | L3mbtl2 |
| GACCTGACATCTTGATGCCG | L3mbtl2_R1.Br_54141 | |
| GTCCGCGCTGTCTACACAGA | L3mbtl2_R1.Br_54142 | |
| ATGGAACCATCAGAAACCGG | L3mbtl3_R1.Br_60900 | L3mbtl3 |
| TAGCAACACAGATGACAGAA | L3mbtl3_R1.Br_60901 | |
| GTTGCTACTCTGATGAACAG | L3mbtl3_R1.Br_60902 | |
| GCACTCCCAGACCTTCACGC | Las1l_As_43773 | Las1l |
| TTGTGAATCTTATCTCAGAG | Las1l_As_43774 | |
| TCCTGAGCCAGGTATTTGAG | Las1l_As_43775 | |
| TTCAAGTGACCACTCCACAG | Lbr_R2.Br_46448 | Lbr |
| GCGCCCTGGATTGATTGGAT | Lbr_R2.Br_46449 | |
| GAGCTTGAACGAAAAACCAG | Lbr_R2.Br_46451 | |
| CGGCAAACCCATGTTTACCC | Ldb1_R2.Br_9730 | Ldb1 |
| CGTGCTCAAGCACCCCAAGG | Ldb1_R2.Br_9731 | |
| AAAATGCTTCGGAAGTAGCG | Ldb1_R2.Br_9732 | |
| GTAGAAGGCAAATACATGCA | Lrrfip1_R2.Br_9993 | Lrrfip1 |
| CGAGTCTCAGCGGCAATACG | Lrrfip1_R2.Br_9994 | |
| GCTGAACCAGATCGCGCGCG | Lrrfip1_R2.Br_9996 | |
| TTTGAGTAACATCGCCAGAG | Lrrfip2_R2.Br_36650 | Lrrfip2 |
| ACACCTCACTGAGTGAACTG | Lrrfip2_R2.Br_36651 | |
| GAAGAACAATCTGATCTACC | Lrrfip2_R2.Br_36652 | |
| CATGAAGGACCTGTTTACGG | Maml1_R2.Br_47616 | Maml1 |
| TCACACAACTAGTCAACCTG | Maml1_R2.Br_47617 | |
| CAGACCCTTGGTTATTCACG | Maml1_R2.Br_47618 | |
| TGGTTGCAAAGACTGGTTTG | Maml2_R2.Br_68568 | Maml2 |
| GGCAAATAACGGTAACAATG | Maml2_R2.Br_68569 | |

| | | |
|---|---|---|
| AGTGAGTTTAAACTCTAACC | Maml2_R2.Br_68571 | |
| AGCACAGCACGGTAGTCGAG | Maml3_R1.Br_74088 | Maml3 |
| TCCGCATTAAAATGGGTCAG | Maml3_R1.Br_74089 | |
| AATAGACGAACTGGCCAACA | Maml3_R1.Br_74090 | |
| GCTCCTGTATGACAGCCCCA | Mapkapk3_As_41299 | Mapkapk3 |
| ACGGGAACCCAAGAAGTACG | Mapkapk3_As_41300 | |
| GCTCACCTTGACATCTCGGT | Mapkapk3_As_41301 | |
| GTCCGGGCCTCGATCCGCTG | Mau2_R1.Br_40926 | Mau2 |
| GTCAGAGCTGTATTGTCAAG | Mau2_R1.Br_40927 | |
| CATTATTCCTGCTCAGCAAG | Mau2_R1.Br_40928 | |
| CGATAACGATGACATCGAGG | Max_As_41091 | Max |
| TGCACTGGAACGAAAACGTA | Max_As_41092 | |
| CACTCTCCTAGGCTGACAAG | Max_As_41093 | |
| GCAGCGTGCAAGGGAACACG | Maz_As_41087 | Maz |
| AAGGCACGAAGCCATCCACA | Maz_As_41088 | |
| GCTGGGCCTGGACTCCCGGG | Maz_As_41089 | |
| ACGTTGTGCAAAGATTGTCG | Mbd1_R1.Br_10397 | Mbd1 |
| GGGGCCTTATATTCCCCCTG | Mbd1_R1.Br_10398 | |
| ACAGGGTAAGATCACATGCA | Mbd1_R1.Br_10399 | |
| GGATCCTGTCCCTTTCCCGT | Mbd2_B_410671 | Mbd2 |
| TCCTCCTTCTTCCATCCGGG | Mbd2_B_410681 | |
| GCGACTCCGCCATAGAGCAG | Mbd2_B_410691 | |
| GCAGGAGGAGGGGGAGAGCG | Mbd2_B_41070 | |
| TCTTCCCTTTCCCAGCCCTG | Mbd3_As_41063 | Mbd3 |
| AGCACCTTCGACTTCCGCAC | Mbd3_As_41064 | |
| AGTCTGCCGTACAGGCAGCG | Mbd3_As_41065 | |
| CCGAAAGGGAGCATCAATCC | Mbd4_R1.Br_10409 | Mbd4 |
| GCCTGTGCAAGGGAGACTGT | Mbd4_R1.Br_10410 | |
| TCAGAGTCGCCAGAAAGCAG | Mbd4_R1.Br_10411 | |
| ATAGGGTCAAGAATGCCAAG | Mbd5_B_41047 | Mbd5 |
| GCAGCTATCCAGGTTCCTGT | Mbd5_B_41048 | |

| | | |
|---|---|---|
| ATTTGATCCCCCACAACCCG | Mbd5_B_41049 | |
| CCAACAGGTGAAGGTCAAAG | Mbd5_B_41050 | |
| CCTACAACTCCACTTAACGG | Mbd6_R1.Br_50088 | Mbd6 |
| CCAGCACACCAAATAAACTG | Mbd6_R1.Br_50089 | |
| AGACTGCTAGCAAGGAATGG | Mbd6_R1.Br_50090 | |
| CAGCCCCCTCCTCACCCGCG | Mbip_As_41039 | Mbip |
| AAGAGCCATACATCCCACCG | Mbip_As_41040 | |
| AGGAGCGAAAGACTTCGCAG | Mbip_As_41041 | |
| TTCTTGACCATCCAATATGG | Mbtd1_R1.Br_47540 | Mbtd1 |
| TGAGGCCACTCACTTCTAGG | Mbtd1_R1.Br_47541 | |
| ATGTCGAACAAGAGTAGCAG | Mbtd1_R1.Br_47542 | |
| GAACGAAAGACTTTCCGCTG | Mbtps1_R2.Br_24838 | Mbtps1 |
| ATGGCAAGGTAACCTGACCA | Mbtps1_R2.Br_24839 | |
| TGTTTATAGGAATTACCAGG | Mbtps1_R2.Br_24840 | |
| AAGGCAAGAGGATTACTGGG | Mbtps2_R2.Br_40979 | Mbtps2 |
| GAAGAGGAAGAATAAGGAGA | Mbtps2_R2.Br_40980 | |
| GTTGTACATGAAATTGGACA | Mbtps2_R2.Br_40981 | |
| GCGCTCCTGAACTTCCCGAA | Mcrs1_As_40820 | Mcrs1 |
| TCTGAGCCCAGGCTACCTGG | Mcrs1_As_40821 | |
| GTGACTCCTCATCCTCGGAG | Mcrs1_As_40822 | |
| GGAGATGGATGGAAAAGGCA | Mdc1_As_40792 | Mdc1 |
| GCTCTTGTGGAAGAGTACAT | Mdc1_As_40793 | |
| GTGTCACTGCCCAAACCAAA | Mdc1_As_40794 | |
| CACACGTCACAGCTTCTGTG | Mdfi_R1.Br_10517 | Mdfi |
| AAGATCCACTCACCCTGTAG | Mdfi_R1.Br_10518 | |
| CTGCACTGCCCGGTTCTGAG | Mdfi_R1.Br_10519 | |
| GACACATGTTTGGCTCCGTG | Mdfic_R2.Br_9161 | Mdfic |
| TGTGCATCTTCTGAGAAACG | Mdfic_R2.Br_9162 | |
| AGCACAGGAACCCGGCAAGG | Mdfic_R2.Br_9163 | |
| CCTCCATAGGATCTTGACGA | Mdm2_R2.Br_10530 | Mdm2 |
| ATTACAGCCTGAGTGACGAA | Mdm2_R2.Br_10531 | |

| | | |
|---|---|---|
| TTAGTGGCTGTAAGTCAGCA | Mdm2_R2.Br_10532 | |
| ACATCAGCTTCTATTAACAC | Mdm4_R2.Br_10533 | Mdm4 |
| GAGCAACATATGGTATACTG | Mdm4_R2.Br_10534 | |
| CTTGAGCGATGATACTGACG | Mdm4_R2.Br_10535 | |
| CTGCCCATACTTAGATCCAG | Mecom_R2.Br_40716 | Mecom |
| GCTGGTCACAGTCCTCACAG | Mecom_R2.Br_40717 | |
| GCTGTTCATGAAGAGTGAAG | Mecom_R2.Br_40718 | |
| CAGGCCATTCCTAAGAAACG | Mecp2_R1.Br_10554 | Mecp2 |
| TCTTGACAACAAGTTTCCCA | Mecp2_R1.Br_10555 | |
| CCGTGAAGGAGTCTTCCATA | Mecp2_R1.Br_10556 | |
| CCAACCAACACCTTTCCGGG | Med1_R1.Br_13486 | Med1 |
| TAACTTACCCCCACTCCGCG | Med1_R1.Br_13488 | |
| GTGAGCTGTAAACTCTACAA | Med1_R1.Br_13489 | |
| ACAGCAGCTGCACGATATCA | Med10_R1.Br_21326 | Med10 |
| ACACCAAAGAATGCCTGGAG | Med10_R1.Br_21327 | |
| GGAGAAGTTTGACCATCTGG | Med10_R1.Br_21329 | |
| CCAAGGAAAAGACTAACGAG | Med11_R1.Br_27658 | Med11 |
| CCTACAGTCTGGCGAACGAG | Med11_R1.Br_27660 | |
| AGTGAGGTAGCGGATTTGAG | Med11_R1.Br_27661 | |
| CTGTTGGAAAACCTCGATTG | Med12_R1.Br_26418 | Med12 |
| GAGGACCGGATTCCAACAGT | Med12_R1.Br_26419 | |
| CATATCTTTACAGTACTCAG | Med12_R1.Br_26421 | |
| CATCACTAACTGATACACGA | Med13_R1.Br_70568 | Med13 |
| ATAGGACGTAACACAGACTG | Med13_R1.Br_70570 | |
| TCATAACCCAACAAAAACGT | Med13_R1.Br_70571 | |
| GTGCGGCCAACAATGTACGG | Med13l_R1.Br_42494 | Med13l |
| TGTGGCCGAAGAATTATGCG | Med13l_R1.Br_42495 | |
| CTTCCATCCTAAACTCCGTG | Med13l_R1.Br_42497 | |
| AACCTCGAGATACCACATCA | Med14_R1.Br_20526 | Med14 |
| AGGCAGGCGTGCATGGACCA | Med14_R1.Br_20528 | |
| CACTAATGTTAATCCGAGAG | Med14_R1.Br_20529 | |

| | | |
|---|---|---|
| GCCCAACTCTAATGTCAGGT | Med15_R1.Br_46068 | Med15 |
| GCTTGTGCTTGTGACACCAA | Med15_R1.Br_46069 | |
| GCCATGAGGGGCCATTCCAG | Med15_R1.Br_46071 | |
| GACCTACGCAATGATGACCA | Med16_R1.Br_54592 | Med16 |
| CTCGTGGGCATCGATAACCA | Med16_R1.Br_54593 | |
| CCTGGACCGTGTATCCGCGG | Med16_R1.Br_54595 | |
| GCAGACCCCTCTAAATCACT | Med17_R1.Br_60320 | Med17 |
| GAGAACACACATCTCTGTAA | Med17_R1.Br_60321 | |
| AATTCAATACAGTCCACAGA | Med17_R1.Br_60322 | |
| GGATGAGGCTTTCCAAACTG | Med18_R1.Br_30254 | Med18 |
| GACCATCTCGTGATCAAGGA | Med18_R1.Br_30255 | |
| GTCCACACAGTTGCGCACCA | Med18_R1.Br_30256 | |
| TTTAATCACACACTACAACC | Med19_R1.Br_72368 | Med19 |
| GAATAGGAGGCTTCTCAATG | Med19_R1.Br_72369 | |
| GAGGCGGAAACCAGACAGCA | Med19_R1.Br_72370 | |
| GCCAGCAAGATTGAGACCCG | Med20_R1.Br_25334 | Med20 |
| AGGTGGGCACGGTCACAATG | Med20_R1.Br_25335 | |
| TTGGTGTCAGCAATGAGGCA | Med20_R1.Br_25337 | |
| ATTGGTGTATTACAGCAGTG | Med21_R1.Br_48948 | Med21 |
| GCAGCTGTAGACTCTTCACT | Med21_R1.Br_48949 | |
| CAACCAGCCAATCCTACAGA | Med21_R1.Br_48950 | |
| ACCGCAAGCTCATCACCCTG | Med22_R1.Br_16930 | Med22 |
| CATTGACCTATATGAGCTGG | Med22_R1.Br_16931 | |
| CAGGTGTCTCGGGCTACTCA | Med22_R1.Br_16932 | |
| ACGTAGAGAACTCATACACC | Med23_R1.Br_35470 | Med23 |
| GGAATACGCGTTACATAGCA | Med23_R1.Br_35471 | |
| GGAGAGAGAGGACCATATGT | Med23_R1.Br_35472 | |
| TAGACAGATGAAGTGGCACG | Med24_R1.Br_19778 | Med24 |
| GCTACACATCGCCAAACTAG | Med24_R1.Br_19779 | |
| GAAGGATTTAAGCTTCCCAG | Med24_R1.Br_19780 | |
| TCTTTGCAGGTACTTCAACG | Med25_R1.Br_41930 | Med25 |

| | | |
|---|---|---|
| GCCTAGCCTAGTCTCCACCG | Med25_R1.Br_41931 | |
| ATTCTTGAGTTCCGGAACAG | Med25_R1.Br_41933 | |
| CGGAGCTCTTACATACCCAA | Med26_R1.Br_36002 | Med26 |
| AGAATCGCAACGACATCCAA | Med26_R1.Br_36003 | |
| TCTTGTAGGAGACACGACTA | Med26_R1.Br_36004 | |
| GTCTTCGACTGCCTTAAGGA | Med27_R1.Br_33730 | Med27 |
| GAACGTTTGAGCAACCTGGT | Med27_R1.Br_33731 | |
| GAAGGTGTTGAAAGTCATTG | Med27_R1.Br_33732 | |
| AGTGCTGTTAGATGGTCTCG | Med28_R1.Br_29694 | Med28 |
| CGAGTTGGAGTCATCCTTCG | Med28_R1.Br_29695 | |
| ACCAGATCAAGTTATCAAAG | Med28_R1.Br_29697 | |
| ACAGCCACAACCGACCCAGT | Med29_R1.Br_30270 | Med29 |
| AGCTGGAACTCTGCTTGGTG | Med29_R1.Br_30271 | |
| GCAACTGAAGGAGAGTCTCC | Med29_R1.Br_30272 | |
| CCAGTATGAGCACTTTCGCA | Med31_R1.Br_30398 | Med31 |
| CTAACATATGTAAGCACTGA | Med31_R1.Br_30399 | |
| ATGATGCTGGAAATCGGCTT | Med31_R1.Br_30400 | |
| AAGATCTGGAGGTCTTGTCG | Med4_R1.Br_30602 | Med4 |
| TCAACTCACCAGGATCTGCT | Med4_R1.Br_30603 | |
| AAGGAAAAGGAGCGGATGGG | Med4_R1.Br_30605 | |
| GTGTGCCCCTTTCAGTCAGA | Med6_R1.Br_40604 | Med6 |
| GCCCCTTTCAGTCAGATGGT | Med6_R1.Br_40605 | |
| ATGAGGTGGTCAAAATGCAG | Med6_R1.Br_40606 | |
| GACTTTGAGAGTCATGATGG | Med7_R1.Br_27782 | Med7 |
| GAATGATAAGATCATCACAT | Med7_R1.Br_27783 | |
| ACATCTTAATAAGAAGTCCT | Med7_R1.Br_27785 | |
| GAACGAGAATCAGAGAGCGG | Med8_R1.Br_44756 | Med8 |
| ATGACCTGGTTTCGGAACAG | Med8_R1.Br_44757 | |
| ATATTACAGCGTCAAACTGA | Med8_R1.Br_44758 | |
| TTGATGACATTGTGAACCAA | Med9_R1.Br_51780 | Med9 |
| TCTCCTCCTTCGCGCCAGCG | Med9_R1.Br_51781 | |

| | | |
|---|---|---|
| CCTGTCGCCCAGCATGGACA | Med9_R1.Br_51783 | |
| CCAGCCATGGTGATGCACGG | Meis2_1 | Meis2 |
| GCGTTGAGGTTGCGTCATCG | Meis2_2 | |
| TCAAAAACCAGAGCTAACAG | Meis2_3 | |
| TGGCCTCACCTACTTCCCGG | Men1_As_40512 | Men1 |
| GGGAGAGGTCCACAGCGCCG | Men1_As_40513 | |
| TCTGCGCTCTATCGACGACG | Men1_As_40514 | |
| AATCGCGGGATACCCCAACG | Meox2_1 | Meox2 |
| AGAGTTCGGGGTAGGACATG | Meox2_2 | |
| GATGTCCTCTCCACCAAGCG | Meox2_3 | |
| CCATTTCCTCTGGCTTCACA | Mettl1_As_40432 | Mettl1 |
| GCGCGCCCACTCCAACCCCA | Mettl1_As_40433 | |
| ATCAGGGTATCTGGGAAGAG | Mettl1_As_40434 | |
| CGTGAACATCGACATCAGCG | Mettl13_B_40420 | Mettl13 |
| CATCCTGTCCACCTGACGCA | Mettl13_B_40421 | |
| ACACAGCTCAAAGATCTGAA | Mettl13_B_40422 | |
| GCCACGCTACACCCTCCACG | Mettl13_B_40423 | |
| CAGACCTCAGAATTTCATCA | Mettl14_As_40416 | Mettl14 |
| CATCTGCTCCAAACTCAAAA | Mettl14_As_40417 | |
| CTGAAACCAGAGAAACCTGC | Mettl14_As_40418 | |
| TCATTAGCAATGCCTCCGCT | Mettl15_As_40412 | Mettl15 |
| GCATACTGAATCTAAAGCTG | Mettl15_As_40413 | |
| GTTGGGCCAGTTCAGCCAAG | Mettl15_As_40414 | |
| ACCTAGTTCTGTAAACACCG | Mettl16_As_40408 | Mettl16 |
| GTGTTTACAGAACTAGGTGG | Mettl16_As_40409 | |
| ACTGTACCTTGGCTTCCAAT | Mettl16_As_40410 | |
| CCACACACACGTATTCACGG | Mettl17_As_40404 | Mettl17 |
| ACTGGGGTCCTCGAAGGCTG | Mettl17_As_40405 | |
| GGTGTGTCCCAGGTAGACAA | Mettl17_As_40406 | |
| TCTGACCTGTCAGCAATAGG | Mettl18_As_40400 | Mettl18 |
| TGTGTCTTAGAAAGTCAGAA | Mettl18_As_40401 | |

| | | |
|---|---|---|
| GACACAAGTTCCATCATCGA | Mettl18_As_40402 | |
| GCAGTTACCAGGCATTGTGA | Mettl2_As_40396 | Mettl2 |
| GGACAACGTGAAGTGGTCCG | Mettl2_As_40397 | |
| TCACTCCTCTGCTTCTCCAA | Mettl2_As_40398 | |
| CCAGTTGCCTCCAGTCCTGA | Mettl21a_As_40388 | Mettl21a |
| GGTACCCTATGAGGAGAGCG | Mettl21a_As_40389 | |
| CCAGTCCTGACGGATCTGGA | Mettl21a_As_40390 | |
| CGTACTCCTGGTCTCCCACG | Mettl22_As_40376 | Mettl22 |
| TTTGAAGGTGACCTCGTCCA | Mettl22_As_40377 | |
| GGACTCTTGGACAGACTCGG | Mettl22_As_40378 | |
| CGGGAGGAGCCAGTTCCCGG | Mettl23_R1.Br_40546 | Mettl23 |
| GAGGCCGGGCTCGCACGTAG | Mettl23_R1.Br_40547 | |
| TCGAAGTTGTAGGACTGACA | Mettl23_R1.Br_40548 | |
| ACGCTGCACGACAAGTTACA | Mettl25_As_40364 | Mettl25 |
| AGAGAGTTGAAAGTGCCAAA | Mettl25_As_40365 | |
| TGAGAAATCAGGCAAACCAT | Mettl25_As_40366 | |
| TTACCGTAGAGATGGCAAGA | Mettl3_As_40360 | Mettl3 |
| GAGTTGATTGAGGTAAAGCG | Mettl3_As_40361 | |
| CATCCGTCTTGCCATCTCTA | Mettl3_As_40362 | |
| CAATTGTGATTGATCCACCA | Mettl4_R2.Br_43088 | Mettl4 |
| AGTGACTGACTATAAACAAG | Mettl4_R2.Br_43089 | |
| TCTTCGAAGTTAATCCAAGA | Mettl4_R2.Br_43091 | |
| TGGAGTACTTAGCATCGGAG | Mettl5_R2.Br_41626 | Mettl5 |
| TTCTAGAACAGTATCCCACC | Mettl5_R2.Br_41627 | |
| CGATGACATTGAAAACAAAG | Mettl5_R2.Br_41629 | |
| CATCTCTCTGCATTGTACAA | Mettl6_As_40348 | Mettl6 |
| GGAGTTGAGATCATGTAGGG | Mettl6_As_40349 | |
| AAGGGCAGTTGACTACGTGA | Mettl6_As_40350 | |
| GGTACAGACCACCAGCCGCA | Mettl7a1_B_40344 | Mettl7a1 |
| CAGAAACATGGGAAGCGCCA | Mettl7a1_B_40345 | |
| GCAGATGGCGAGCCAAAAGC | Mettl7a1_B_40346 | |

| | | |
|---|---|---|
| ATAGAACTTGAAGTTGGCTC | Mettl7a1_B_40347 | |
| TGTGTTCAAAAACTTTGGCG | Mettl8_As_40333 | Mettl8 |
| GGTGGACGCCACTCTGGCAT | Mettl8_As_40334 | |
| GTTAAAATCCTCGACCCCAG | Mettl8_As_40335 | |
| CATCCTCCGCGCCAGCCACA | Mettl9_As_40329 | Mettl9 |
| CTGTACGTGAACATGACTAG | Mettl9_As_40330 | |
| TCATGTTCACGTACAGGGAG | Mettl9_As_40331 | |
| TCATTGAGTTGGCTTCTCAG | Mga_As_40193 | Mga |
| TGAGATTTCGACAAGCCTGG | Mga_As_40194 | |
| TGACCTCTGATGTACATACG | Mga_As_40195 | |
| GCTACTCCTGAGGTGCTCGG | Mgmt_As_40133 | Mgmt |
| GCTGTGAGCGAGGCCTGCAT | Mgmt_As_40134 | |
| ACAGCTCCATCTTCCCCAAA | Mgmt_As_40135 | |
| TCAAAATATTTACATCGACG | Mier1_R2.Br_36558 | Mier1 |
| CTGAAATAGAGGATCTTGCG | Mier1_R2.Br_36559 | |
| TTTAATGTCAAAGCAGCTCG | Mier1_R2.Br_36560 | |
| CGTGAAGTGTAATTTCAACG | Mier2_R2.Br_35766 | Mier2 |
| TGTACAGGGCAGTGAAACGG | Mier2_R2.Br_35767 | |
| GGTTCACACAGGTACGCACG | Mier2_R2.Br_35769 | |
| AAACGCTATAACCATCACCC | Mier3_R2.Br_55636 | Mier3 |
| CTATGAATCTACAATTCCAG | Mier3_R2.Br_55638 | |
| TTCTCAATGAGGTCTCAACA | Mier3_R2.Br_55639 | |
| AGCTTCCCTAAACCCAAGCG | Mllt1_R1.Br_26866 | Mllt1 |
| CTGCTCTCTGAGTCCTTGCG | Mllt1_R1.Br_26868 | |
| CTAAAAAGAGTGGGTCAAAG | Mllt1_R1.Br_26869 | |
| TGCATGACATGTAATAAACA | Mllt10_R1.Br_10721 | Mllt10 |
| GGCCGAGGAAGCTCGCCCCG | Mllt10_R1.Br_10722 | |
| TTGCTGAATGCAATACACAA | Mllt10_R1.Br_10723 | |
| GCTGCTATCCTTGCTCTCGT | Mllt11_R1.Br_25338 | Mllt11 |
| TAGGTGTATCTGACAGGCCC | Mllt11_R1.Br_25339 | |
| CATCCCAGAACTGGATCTGT | Mllt11_R1.Br_25340 | |

| | | |
|---|---|---|
| AACTTTCAATAACCCCACGG | Mllt3_R1.Br_35410 | Mllt3 |
| AAGGAAAAAGAGTAGCTCGG | Mllt3_R1.Br_35411 | |
| TCCACGATGTCATCAAACGG | Mllt3_R1.Br_35412 | |
| AAGGCTCATCGGGTGCACAG | Mllt4_R1.Br_10729 | Mllt4 |
| GTTTCAGTTTCAGATCGTCG | Mllt4_R1.Br_10730 | |
| CACAAACTTAAACACATGCG | Mllt4_R1.Br_10731 | |
| TCACATTTCCTGCAGAACCA | Mllt6_B_39821 | Mllt6 |
| GTGCGCCCTCTACATCCCGG | Mllt6_B_39822 | |
| GGAGACCTCTGAGAGTAGCA | Mllt6_B_39823 | |
| TAAGTCTCCCCATATTACCG | Mllt6_B_39824 | |
| GTAGCACCTGGAAATGGAGA | Morf4l1_R1.Br_17442 | Morf4l1 |
| TGAAGGGAACAATATGCAGA | Morf4l1_R1.Br_17443 | |
| GCCTCTTCTCTATGAAGCAA | Morf4l1_R1.Br_17444 | |
| AGGACTGGGACTTGGTTACG | Morf4l2_1 | Morf4l2 |
| ATCACTGGCAGTAAACAGAG | Morf4l2_2 | |
| GGCACGGGCTGACCCCACTG | Morf4l2_3 | |
| GTGAAGTCCCCCAGCCCCCT | Morf4l2_B_39513 | |
| CCAGCCCCCTCGGAAGAAAA | Morf4l2_B_39514 | |
| TCCCTGAAGAATTAAAACCG | Morf4l2_B_39515 | |
| ACCAGCTGACTTCTTTCCCG | Morf4l2_B_39516 | |
| AAAGCAAAGAAACCAAGACA | Mphosph8_R2.Br_41563 | Mphosph8 |
| TAATGATTCCCTCATTCTAG | Mphosph8_R2.Br_41564 | |
| GATCCTGGACATGAAGTGCG | Mphosph8_R2.Br_41565 | |
| TAAACAAGCCGGCATCGGCG | Msl1_R2.Br_39970 | Msl1 |
| CCTTGAGCTTACGATGCCGG | Msl1_R2.Br_39971 | |
| TATAAGGAAGAGCCCTCTCG | Msl1_R2.Br_39972 | |
| GCTGTGGAGAAAAGAATGGA | Msl3_R2.Br_38706 | Msl3 |
| GCTGCTTCTTGAGAACATCG | Msl3_R2.Br_38707 | |
| GCTGCGTTCAAGAAAGGAAA | Msl3_R2.Br_38708 | |
| ACTGTTACTACATCAACCGC | Msl3l2_R2.Br_39422 | Msl3l2 |
| AGTGGCGCACGTAGCACTCA | Msl3l2_R2.Br_39424 | |

| | | |
|---|---|---|
| CGCTGGCTCGCTCCTCAGTG | Msl3l2_R2.Br_39425 | |
| TTTGGCCTCCACATTCCCGT | Mta1_As_38606 | Mta1 |
| TGGATTGCAGCAGCTCCGTG | Mta1_As_38607 | |
| CATGGCCGCCAACATGTACA | Mta1_As_38608 | |
| GGAGGAATCAAAACAGCCAG | Mta2_As_38602 | Mta2 |
| GTTCACTTACAAAGTCCTGG | Mta2_As_38603 | |
| AGACACTTCTGGCTGATCAA | Mta2_As_38604 | |
| GCAACCCGTACCTCATCAGA | Mta3_As_38598 | Mta3 |
| TACCAAGGTGATGTCTCGGG | Mta3_As_38599 | |
| GGATTCTGTCTCATTCAGCA | Mta3_As_38600 | |
| TCAAACGTCTACCGTTACAG | Mtf2_B_38539 | Mtf2 |
| GACCATTTCCCCACTCCCAG | Mtf2_B_38540 | |
| GGTCATGCTCACCACTGTAA | Mtf2_B_38541 | |
| TCTTCTTGTGAATAACACTT | Mtf2_B_38542 | |
| ACAGGCACTAGACCTGATCG | Mybbp1a_R2.Br_12422 | Mybbp1a |
| CCAGAGATAAGTACATACGT | Mybbp1a_R2.Br_12423 | |
| AGATATGACCAAATACACTC | Mybbp1a_R2.Br_12425 | |
| CTGCTCGCGCTTCGAGTCGG | Mycbp_R2.Br_24442 | Mycbp |
| CGAAGCGCGAGCAGTTCCGG | Mycbp_R2.Br_24443 | |
| GGATACTCCAGAGCACTGGT | Mycbp_R2.Br_24445 | |
| AATAAGGATGATATTCGCTG | Mycbp2_R2.Br_48128 | Mycbp2 |
| GATGAAAGTCTATTCCAAGG | Mycbp2_R2.Br_48129 | |
| AATGGCACAAACCAAACCAG | Mycbp2_R2.Br_48130 | |
| TACACATTATAGGATCCGGA | Myef2_R2.Br_11178 | Myef2 |
| ATGTCTTACTATCATGTGAA | Myef2_R2.Br_11180 | |
| ATGGGAAATTTGGGTCCAAG | Myef2_R2.Br_11181 | |
| CCAGTAGCTGGTAAAAGACG | Myo1c_As_37951 | Myo1c |
| GTCTCCTCACCTCTAACACG | Myo1c_As_37952 | |
| TGAACAGGCGGCATATGCAA | Myo1c_As_37953 | |
| TCTTGATTCAACGTGGAGAA | Mysm1_R1.Br_37823 | Mysm1 |
| TGAGAAATACAATAAAGTGG | Mysm1_R1.Br_37824 | |

| | | |
|---|---|---|
| GCATAATTCTCCTTGATTGG | Mysm1_R1.Br_37825 | |
| ACCCATCTATAAATTACCAG | Nab1_R2.Br_11318 | Nab1 |
| CAAACACCTTACATTGCATG | Nab1_R2.Br_11319 | |
| CTGGCTCCGACCTTACCTGG | Nab1_R2.Br_11320 | |
| AAGTTGTCGCCACTCCCTGG | Nab2_R2.Br_11322 | Nab2 |
| ATCTTCCGGAGTTTCCCCAG | Nab2_R2.Br_11323 | |
| GCCGCCTGGACTCCAAAAGG | Nab2_R2.Br_11325 | |
| CCTCTAGGTATGCATCAATG | Nabp1_R2.Br_49284 | Nabp1 |
| TCTGTGTGGGATGAGATCGG | Nabp1_R2.Br_49285 | |
| CGAGTGACCAAAACCAAAGA | Nabp1_R2.Br_49286 | |
| CAAGACAAAGGACGGACACG | Nabp2_R2.Br_35170 | Nabp2 |
| TGGGCAACCTGATCCAACCT | Nabp2_R2.Br_35171 | |
| GCCTGCTGCGTGTTATACTC | Nabp2_R2.Br_35172 | |
| ACACACCTGGGCTTGCTGCG | Naca_R2.Br_11326 | Naca |
| TGAGTCAGTACCAGAGCTCG | Naca_R2.Br_11327 | |
| GGGTCTTCGACAGGTTACAG | Naca_R2.Br_11329 | |
| AGGAAGTTCATGATCTCGAG | Nap1l1_R2.Br_23011 | Nap1l1 |
| CTGGTCCGTCAAAAGAAAAG | Nap1l1_R2.Br_23012 | |
| ATGCGAGTGGAAACCAGATG | Nap1l1_R2.Br_23013 | |
| CCACAGATGCAGAGTCAGCG | Nap1l4_R2.Br_11354 | Nap1l4 |
| GCTTGGTCACTACTCACCAT | Nap1l4_R2.Br_11355 | |
| AGATGGTGAACCAGAACTCA | Nap1l4_R2.Br_11356 | |
| CCTCGGCAGATAAAATGGAG | Nasp_B_37519 | Nasp |
| AGCCGCTGCTGCCATCGCCG | Nasp_B_37520 | |
| CAGTCCCGCTTACTTGTCAG | Nasp_B_37521 | |
| GTCAGCGGAAACCAGCTCCG | Nasp_B_37522 | |
| GGCTCCTTAGACCAGATGAG | Ncapd2_B_37419 | Ncapd2 |
| ATTCTGCTAGACCATCTGGA | Ncapd2_B_37420 | |
| GCTGAGTCGAGACACAGCAG | Ncapd2_B_37421 | |
| TTCCTCCCAGTCAAAGCCAA | Ncapd2_B_37422 | |
| GTTATGGGTACGGCTTTGCG | Ncapd3_B_37415 | Ncapd3 |

| | | |
|---|---|---|
| TTTCAGGTAATCTAGTTCCA | Ncapd3_B_37416 | |
| TTATGGCTAGGGGGGCACAG | Ncapd3_B_37417 | |
| GCTTCCGCACAGATATGGCA | Ncapd3_B_37418 | |
| TTACCTTAAGTATGCTATGG | Ncapg_R2.Br_23543 | Ncapg |
| TCATCCTATAGTAAGAAACT | Ncapg_R2.Br_23544 | |
| AGCTCACTAATAAACTACTT | Ncapg_R2.Br_23545 | |
| TGATGAATGCAGCACGACTG | Ncaph_R2.Br_54348 | Ncaph |
| GGAAGAAGTTATTTCCCTCG | Ncaph_R2.Br_54349 | |
| GCGAAACACATAATGAGGTG | Ncaph_R2.Br_54350 | |
| CAGTGTTGTACCTATACAAG | Ncaph2_R2.Br_22674 | Ncaph2 |
| CCGTTCCTAGACACTTCCAT | Ncaph2_R2.Br_22675 | |
| ATTTCTCACCTGTAGCCGGG | Ncaph2_R2.Br_22676 | |
| GTACCAACTCCTGGTAAAAA | Ncl_As_37335 | Ncl |
| TTGTAGGCTGGCAAAACCCA | Ncl_As_37336 | |
| TACCTTTGCGAGCTTCACCA | Ncl_As_37337 | |
| GATGCTAATTCACCCTCCAG | Ncoa1_R1.Br_11415 | Ncoa1 |
| ACCACTAGTAATGCCAACTG | Ncoa1_R1.Br_11416 | |
| AGTTTGGACAACCAGGAGCG | Ncoa1_R1.Br_11417 | |
| AGTGCATAGTTACTACCCTG | Ncoa2_R1.Br_11418 | Ncoa2 |
| GGGAGGATTCATATTAACTG | Ncoa2_R1.Br_11419 | |
| AGTCAGATGTGTCGTCCACG | Ncoa2_R1.Br_11420 | |
| CAAGGAGAAACGATACACGG | Ncoa3_R1.Br_11422 | Ncoa3 |
| TCGTCCTCCATATAACCGAG | Ncoa3_R1.Br_11423 | |
| AGAACATCATGATTTCCCCT | Ncoa3_R1.Br_11424 | |
| CGTCGCTGATTGTTGCGCCG | Ncoa4_R2.Br_20762 | Ncoa4 |
| TTCAAGAAGTCAGCATCCAG | Ncoa4_R2.Br_20763 | |
| AATTAAAGATAATTTACGAG | Ncoa4_R2.Br_20765 | |
| GGGTGGATGACTATTGCCGG | Ncoa5_R1.Br_57824 | Ncoa5 |
| GTAGAGAGGAGCTTTATCGT | Ncoa5_R1.Br_57825 | |
| TAACAGATACCTCACTGCCG | Ncoa5_R1.Br_57827 | |
| TCACCCTCCTCCTTTCAGCA | Ncoa6_As_37303 | Ncoa6 |

| | | |
|---|---|---|
| TGTCCTGCTGAGAAGCCAGG | Ncoa6_As_37304 | |
| GCACAATGACTACAAATCAA | Ncoa6_As_37305 | |
| TGAACCATAGGATTCCGACC | Ncoa6_R1.Br_24702 | |
| GCTGTTATGACGCCCCAGGG | Ncoa6_R1.Br_24703 | |
| CAATGACTACAAATCAAGGG | Ncoa6_R1.Br_24704 | |
| AGGCCTTAAAACCCATCGAG | Ncoa7_R1.Br_53200 | Ncoa7 |
| AGTTCAACGTTACAGCAGAG | Ncoa7_R1.Br_53201 | |
| GGTACACTCTGGGTAAAAGG | Ncoa7_R1.Br_53202 | |
| TTTCCAGCGTGTTAGTGCTG | Ncor1_As_37295 | Ncor1 |
| GCTCTGTGCTGAAAGCTCCT | Ncor1_As_37296 | |
| TTGCTTTCAGAATTTCACCC | Ncor1_As_37297 | |
| GCACTATCGTGGAGGCCCAG | Ncor2_As_37291 | Ncor2 |
| GCAGATAGCCCGGTCCCACA | Ncor2_As_37292 | |
| ACGCAGCACCTACAGCCAGA | Ncor2_As_37293 | |
| TTATCCCTGAGAGGACCGTG | Nek6_As_36915 | Nek6 |
| ATTGAGAAGAAGATTGGCCG | Nek6_As_36916 | |
| CTGCACGAAGTACTTCCACA | Nek6_As_36917 | |
| CACGCTGTACCGCCGCACCG | Nek9_As_36903 | Nek9 |
| TGGCCTCCCCAAAGGCACCG | Nek9_As_36904 | |
| GGTATATAGTTCCTTCTCCA | Nek9_As_36905 | |
| CATCCTTGCCCTATTCAGTG | Nfrkb_R1.Br_60388 | Nfrkb |
| GGGTGTTACAGGCGTAACTG | Nfrkb_R1.Br_60389 | |
| AGGACTACCCCAGGTTCGGG | Nfrkb_R1.Br_60390 | |
| GGTGAGTCAGAGCGACATCG | Nipbl_R1.Br_36586 | Nipbl |
| CAACAGTCAATAAGCCCGAG | Nipbl_R1.Br_36587 | |
| AGGGTAAACTAACCTCCATG | Nipbl_R1.Br_36588 | |
| ATCCTTAGGACTCTCCACTG | Nmi_R2.Br_27054 | Nmi |
| CACAAAATGTGATATCGATG | Nmi_R2.Br_27055 | |
| GCAGAGATGGACGATATGAG | Nmi_R2.Br_27056 | |
| GGTATGTCTTGACATCCACA | Noc2l_As_35467 | Noc2l |
| CCACCTTTCCACCATGGCGA | Noc2l_As_35468 | |

| | | |
|---|---|---|
| CCGCCGCTCCCGGTTCCACG | Noc2l_As_35469 | |
| GTAGTCATTGTGGATGACCG | Nono_R2.Br_23026 | Nono |
| TACTTACGTAGTCAGCAAGA | Nono_R2.Br_23027 | |
| AGTGGATCGGAACATCAAGG | Nono_R2.Br_23028 | |
| TGTAATTTCAAAGCCCCCTA | Npm1_R2.Br_11830 | Npm1 |
| GGACGATGATGAGGACGATG | Npm1_R2.Br_11831 | |
| AGATGAGTTACACATCGTAG | Npm1_R2.Br_11832 | |
| GTTGCGATTGATGCGAACGA | Nr1d1_R2.Br_55080 | Nr1d1 |
| GAGTAGGTGAGGTCTCTAGA | Nr1d1_R2.Br_55081 | |
| CACAACAGCTGACACCACCC | Nr1d1_R2.Br_55082 | |
| ATATCGCGAATACTACGTAG | Nsd1_R1.Br_11938 | Nsd1 |
| GAATTGCTCGTTAAGACACC | Nsd1_R1.Br_11939 | |
| GAGGCGCAACCGATTCAGAG | Nsd1_R1.Br_11941 | |
| AGGAAGGCCCAAACAAGACA | Ntmt1_As_34683 | Ntmt1 |
| CAAAGCCAAGACCTACCTGG | Ntmt1_As_34684 | |
| GCTCCCGGAAGTTTCTGCAG | Ntmt1_As_34685 | |
| GGATGAGAAGGACTGTAGAA | Nudt21_As_34527 | Nudt21 |
| GAGCTGTCCTTCTCATAGAG | Nudt21_As_34528 | |
| CCCGTCTGCGAGCGATTGGG | Nudt21_As_34529 | |
| ACGCAGCGTGTAGAGCAACG | Nudt8_1 | Nudt8 |
| TCTCCTCTCCGCGAACCCCA | Nup133_As_34459 | Nup133 |
| CTATTGGCCCAGCCTCGCCA | Nup133_As_34460 | |
| CTATTCCTAGAGAAAAGGAT | Nup133_As_34461 | |
| TGACCAGCTCAACAGCACGG | Nup153_As_34455 | Nup153 |
| ACATCCTCTGCGTTCCCAAT | Nup153_As_34456 | |
| ATAACATCTCAACTACCAGT | Nup153_As_34457 | |
| TCATAGATACTTCTCACAGG | Nup35_As_34423 | Nup35 |
| TTTCCATTACTCCTACTGAA | Nup35_As_34424 | |
| GGAGTTGTAGCTCCAACAAG | Nup35_As_34425 | |
| CTTCAGAGTTGATCACAAGG | Nup43_As_34415 | Nup43 |
| GACCACAGAGACCTTTGCCA | Nup43_As_34416 | |

| | | |
|---|---|---|
| AGAAAATTAGCAAAACCCGC | Nup43_As_34417 | |
| CTTTACATTGGAGAAGGGCG | Nup50_As_34411 | Nup50 |
| GCTTGTGATAGGCATTCCCA | Nup50_As_34412 | |
| CCATTTGTCAGTCCTTCCAG | Nup50_As_34413 | |
| ACTGGTGAATGAGAACCCAG | Nup62_As_34403 | Nup62 |
| CTTCTGGCACCGGCACCGGA | Nup62_As_34404 | |
| ACTGGGTTCTCATTCACCAG | Nup62_As_34405 | |
| GCAGCCTCGTCACTAAGAAA | Nup85_As_34394 | Nup85 |
| CCTTGCAGAGAGACTCCAGA | Nup85_As_34395 | |
| TCATCAGGTCCCCAAGGACG | Nup85_As_34396 | |
| ACGAAAACCTTACCGACACA | Nupr1_As_34370 | Nupr1 |
| GGTTTAGAGGTTGCTGGGAA | Nupr1_As_34371 | |
| CTAAACCTAGAGGATGAAGA | Nupr1_As_34372 | |
| CTGGACAAGTCTAACCCGGT | Padi4_R1.Br_12774 | Padi4 |
| TCCAGTCAAGAAGAGTACCA | Padi4_R1.Br_12775 | |
| AGAATTCTCATCGGGAATAG | Padi4_R1.Br_12776 | |
| TAGAGCCGCTGGATCTCGGA | Pagr1a_As_29049 | Pagr1a |
| GCCTTACTAGCCGAGACAGA | Pagr1a_As_29050 | |
| TGTCTCGGCTAGTAAGGCCG | Pagr1a_As_29051 | |
| AGCTGCCTGGAGATGTCGCA | Pax5_B_286891 | Pax5 |
| GATGGAGTATGAGGAGCCCG | Pax5_B_286901 | |
| CTACCTGTCCTGATGGTCCG | Pax5_B_286911 | |
| AGACAGGAAGCATCAAGCCG | Pax5_B_28692 | |
| AGTGCTGTCTCTCAAACACG | Pax5_R1.Br_125381 | |
| TGGAGGAGTGAATCAGCTTG | Pax5_R1.Br_125401 | |
| TGTTATCTAGGACTCGCCGT | Paxbp1_R2.Br_30566 | Paxbp1 |
| ACAAAGACAAAAGATCGCTG | Paxbp1_R2.Br_30567 | |
| ATGCCATAGGATGCACCGTA | Paxbp1_R2.Br_30568 | |
| TCCAGAGGAGCTATTCCGCG | Paxip1_As_28665 | Paxip1 |
| CCGCGAGGTCAAGTACTACG | Paxip1_As_28666 | |
| TTGATTGTCCCAGAGCCAAA | Paxip1_As_28667 | |

| | | |
|---|---|---|
| TTGCTTTGCACATGGCCAGA | Pbk_As_28657 | Pbk |
| AAGCTTCTGCATAAATGGAG | Pbk_As_28658 | |
| TCTGCATAAATGGAGAGGCA | Pbk_As_28659 | |
| GGAGCTGTAGGAATCACCAA | Pbrm1_As_28641 | Pbrm1 |
| GGTCTCTCCGAGTCTCAAGG | Pbrm1_As_28642 | |
| GTATGGATCAGAGAGTGAAG | Pbrm1_As_28643 | |
| CAATAAAATAGCCCCCACAG | Pcgf2_R1.Br_19094 | Pcgf2 |
| TGCAGCATACCCCCTGACAG | Pcgf2_R1.Br_19095 | |
| GGACCTGGACGTCACACATG | Pcgf2_R1.Br_19097 | |
| ACCCGATATACTGTAATGGG | Pcgf3_R1.Br_34666 | Pcgf3 |
| AGGGAATTCTATCACAAACT | Pcgf3_R1.Br_34667 | |
| TCACTGTGGTTGCGTCAATG | Pcgf3_R1.Br_34668 | |
| CTGCAGGCTGGATAACACGT | Pcgf5_R1.Br_42394 | Pcgf5 |
| TTATACAGATGATATTTCGA | Pcgf5_R1.Br_42395 | |
| AGAAATACGTACATGTATGG | Pcgf5_R1.Br_42396 | |
| TGTAGGGGGTCAGCTCGACA | Pcgf6_R2.Br_36454 | Pcgf6 |
| ATGAAGACACTCTGTAATGG | Pcgf6_R2.Br_36455 | |
| CCTCGAAGCGGCCTCTCAAG | Pcgf6_R2.Br_36456 | |
| GCACCTAGAGCATGTCATAG | Pdcd2_R2.Br_12670 | Pdcd2 |
| AGCATCAGACATTAGACTGG | Pdcd2_R2.Br_12671 | |
| ATTCTGGAAACAGGAAGTTG | Pdcd2_R2.Br_12673 | |
| CAACCTGCTTGCAGACCAGA | Pdp1_As_27913 | Pdp1 |
| GTTCTCCCGTGGCCTACAAA | Pdp1_As_27914 | |
| GCTAGAGATTGAAAATGCAG | Pdp1_As_27915 | |
| AATACTCACGACCGTATCTG | Pds5a_R1.Br_36882 | Pds5a |
| TGGTGAAGAACGCCTAGCTG | Pds5a_R1.Br_36883 | |
| TGATGAATAAGTCAATCGAA | Pds5a_R1.Br_36884 | |
| GCTCCAGTGTAATGATCTCG | Pds5b_R1.Br_46960 | Pds5b |
| AGTGAGGTCACATGATCCTG | Pds5b_R1.Br_46962 | |
| ATGGCGTATTTGGCCTGCCG | Pds5b_R1.Br_46963 | |
| CGACTGCCACCAGAGCGTCG | Per1_R2.Br_12826 | Per1 |

| | | |
|---|---|---|
| CCAGATGCTTATCGCCCAGT | Per1_R2.Br_12827 | |
| GGATGCGCTCGGCAATGAGT | Per1_R2.Br_12828 | |
| GTTCTTCACGATATACTCGG | Per2_R2.Br_12830 | Per2 |
| AGCAGCACCATCGTGCATGT | Per2_R2.Br_12831 | |
| CAACGGGGAGTACATCACAC | Per2_R2.Br_12832 | |
| AGAAGCCAAGCCAATCCCGG | Per3_R2.Br_12834 | Per3 |
| ACTGAAAAATAAAAGCACCA | Per3_R2.Br_12835 | |
| TATGTACATGAACCAAATAG | Per3_R2.Br_12836 | |
| GTGATGACTGGTACGTTCCG | Phb_R2.Br_12926 | Phb |
| TTACCAGGGACACGTCATCC | Phb_R2.Br_12927 | |
| CCAATGCTGGTGTAGATACG | Phb_R2.Br_12928 | |
| TCTATGACCGCCTTCCACTG | Phb2_R2.Br_1445 | Phb2 |
| CATAGTCCAGCCCTAGACGC | Phb2_R2.Br_1447 | |
| AGCGCCGTGCCCATGCCCCG | Phb2_R2.Br_1448 | |
| GGGTACCGACAACTTGCACG | Phc1_R1.Br_4677 | Phc1 |
| GCTCATCCTGATGCCTAATG | Phc1_R1.Br_4678 | |
| CTCAACCTTAGTCAAGCCGG | Phc1_R1.Br_4679 | |
| CCAAGCACAGATGTACCTCA | Phc2_R1.Br_23518 | Phc2 |
| AGGAGATGGTAACTGCAACA | Phc2_R1.Br_23519 | |
| CCACTGTACACAGAAATCTG | Phc2_R1.Br_23521 | |
| GTACCTGCAGCAGATGTACG | Phc3_B_27377 | Phc3 |
| GCAGCACCTGATGCTGCACA | Phc3_B_27378 | |
| GGAGACACAGTTAAAGGCGG | Phc3_B_27379 | |
| GCCTCACCTGGACAGCCGCA | Phc3_B_27380 | |
| CCAGGACTGTCACGTTCCCA | Phf1_B_27369 | Phf1 |
| GCACCCAAACGGCTCAGCCG | Phf1_B_27370 | |
| ATGAAGCTCTCCCTGCCATA | Phf1_B_27371 | |
| TCTTGGCCTTCCCAAAGCCG | Phf1_B_27372 | |
| GCTGCTTTCTTAATGTACTC | Phf10_As_27365 | Phf10 |
| GGACCCATCCATGTAAAAGG | Phf10_As_27366 | |
| GTCGCTGTCGCACCGCCCCG | Phf10_As_27367 | |

| | | |
|---|---|---|
| GGACTGTAAATCATATCCCA | Phf11a_B_27363 | Phf11a |
| TGAGCAACTATATTCGCTGA | Phf11a_B_27364 | |
| GCCATGGACCAAGAGAAGCC | Phf11b_B_27359 | Phf11b |
| TGAGCAACTATATTTCCTGA | Phf11b_B_27360 | |
| CCATGGACCAAGAGAAGCCA | Phf11b_B_27361 | |
| TCCGAATCACATCGAACATG | Phf12_R1.Br_67888 | Phf12 |
| GGGGGACTTCAGAACCACGT | Phf12_R1.Br_67890 | |
| TGAGCATAATGACGTCGATG | Phf12_R1.Br_67891 | |
| GCCGGGACCATTGACAGCGA | Phf13_As_27343 | Phf13 |
| GGGCTTAGCTCTCTGCAAGA | Phf13_As_27344 | |
| AGTACTCCCCCACTTGTAAG | Phf13_As_27345 | |
| AAGAGCTTCCAGGAGAGAAG | Phf14_R1.Br_27339 | Phf14 |
| ATACAAAGTCCAACAGGGGA | Phf14_R1.Br_27340 | |
| CCTACTGTAGTAAAGAGAAA | Phf14_R1.Br_27341 | |
| GTGTAGGCAGTGGTTCCATG | Phf19_R1.Br_39946 | Phf19 |
| TTTCTTACGAAGCAGCTCGG | Phf19_R1.Br_39947 | |
| AGTTGGCATCAACTGCGATG | Phf19_R1.Br_39948 | |
| CATTGTGAAGAAACTATCGT | Phf2_R1.Br_12934 | Phf2 |
| CGGCGAAGAACATCTCGCTG | Phf2_R1.Br_12935 | |
| AGTGATGTGGAGAACTATGT | Phf2_R1.Br_12936 | |
| AAGACGCATGTACAGACTCG | Phf20_R1.Br_57788 | Phf20 |
| GAACGAGCGGGAGTACTCCG | Phf20_R1.Br_57789 | |
| TGTAGAATGTCCCTGCAGTG | Phf20_R1.Br_57790 | |
| TACAACTTATCAGTACCCGA | Phf20l1_R1.Br_61508 | Phf20l1 |
| GGAGTCATCGATATGATGAG | Phf20l1_R1.Br_61510 | |
| CACAACTTAAATCAAGAGGG | Phf20l1_R1.Br_61511 | |
| GCTTTGAGTGAGAAACAGGT | Phf21a_B_27319 | Phf21a |
| GCATTGTCATAGCTACTCCA | Phf21a_B_27320 | |
| CTAGGAGGAGGGATAAACTG | Phf21a_B_27321 | |
| ACAGGCCTCTGGATAAGCTG | Phf21a_B_27322 | |
| GATAGTTCGAAGGTCCGACG | Phf23_R1.Br_44028 | Phf23 |

| | | |
|---|---|---|
| TTACATCCCCCCTAGCAAAG | Phf23_R1.Br_44029 | |
| AAGTTGAAAAAGGCAGATCG | Phf23_R1.Br_44031 | |
| ACTCAGGGGATCCAGACACG | Phf3_R1.Br_53688 | Phf3 |
| TTACCTTGTGCCATTAAACG | Phf3_R1.Br_53689 | |
| TTGGCTCAACTTGCGAAGAG | Phf3_R1.Br_53690 | |
| CTCATCACATATGCGGACCA | Phf5a_B_27303 | Phf5a |
| TGTAACTATGGATCTTACCA | Phf5a_B_27304 | |
| TCTCCTCTCTCTACAGGTGA | Phf5a_B_27305 | |
| CTACGTGCGTCCCTGCACCC | Phf5a_B_27306 | |
| TCCGAGAGAAACCTTCGCAA | Phf6_B_27299 | Phf6 |
| CTTTATCATGCAATGCACAG | Phf6_B_27300 | |
| CATTGTCCTGGAGCAACCAT | Phf6_B_27301 | |
| GGAGCAGTCCTAATGATACC | Phf6_B_27302 | |
| GACCCACCTGGATACACTTG | Phf7_R1.Br_37398 | Phf7 |
| CCAACACAGAACATCCACCA | Phf7_R1.Br_37399 | |
| AAAAACTATCAGGACTAAGA | Phf7_R1.Br_37401 | |
| GGTTTAGTGAAAAAACGTCG | Phf8_R1.Br_70184 | Phf8 |
| CCTGCCATTAGCCATCAACA | Phf8_R1.Br_70185 | |
| TCATTCCTACCTTAAGCACA | Phf8_R1.Br_70187 | |
| TCTTGTTGCTTGGAGTCGGA | Phip_As_27279 | Phip |
| ATATGGCTACTCGTCCAGCA | Phip_As_27280 | |
| ATACCTGTCTTCTACAGGGG | Phip_As_27281 | |
| TAGGACTTGAATGTACGTTG | Pias1_R2.Br_24890 | Pias1 |
| TCTGCAGCTTTATGTCCGGG | Pias1_R2.Br_24891 | |
| GACACAGCAGCGAAACCCGT | Pias1_R2.Br_24893 | |
| TAATACTCACCTAAACTCGT | Pias2_R2.Br_10697 | Pias2 |
| CTAAAGACGTAATGTTCAGG | Pias2_R2.Br_10698 | |
| TCGTACAAGATACACAGACA | Pias2_R2.Br_10699 | |
| AGATGACCAATTAACTACGA | Pias3_R2.Br_58076 | Pias3 |
| ATAGTCAGCCATACCAAGGG | Pias3_R2.Br_58078 | |
| TACAAAAACTCAGAGCCAAG | Pias3_R2.Br_58079 | |

| | | |
|---|---|---|
| TGGCCGAGGACAGATACATG | Pias4_R2.Br_26354 | Pias4 |
| GCATCTTCGCGCTAACACCT | Pias4_R2.Br_26355 | |
| CTGCACGGCAGGGCACCGAG | Pias4_R2.Br_26356 | |
| GCTGAGGTGGAGCTGAAGAA | Pkm_As_26716 | Pkm |
| GGGCAGAGTCAATGTCCAGG | Pkm_As_26717 | |
| GTTTGCATCTTTCATCCGCA | Pkm_As_26718 | |
| AGGGTGCCATGACACTACAC | Pmf1_R2.Br_29790 | Pmf1 |
| TCCTTCCTGACAGCTACGAG | Pmf1_R2.Br_29791 | |
| TGAACCCTGAGGTGACGCAG | Pmf1_R2.Br_29793 | |
| TCACTAGCAAGCCATACAGG | Pml_R2.Br_13318 | Pml |
| CCAGTGGTACCTCAAGCATG | Pml_R2.Br_13319 | |
| GCTGTCTAAGACCCAACCTG | Pml_R2.Br_13320 | |
| CAGACAGCCCCCACTCGCGA | Pnrc1_R2.Br_49136 | Pnrc1 |
| GGCACGGCCCGTCCCCCCCG | Pnrc1_R2.Br_49137 | |
| GGAACCCGAGCGGCGCTAAG | Pnrc1_R2.Br_49138 | |
| TGATAAACCAGCATTCCAGT | Pnrc2_R2.Br_22746 | Pnrc2 |
| GCAAAATAGACAGAAGAGCA | Pnrc2_R2.Br_22747 | |
| AGAAAAAGGAACGAGGACAT | Pnrc2_R2.Br_22748 | |
| GGCATACAAAACGAAGACAC | Pole3_R2.Br_26342 | Pole3 |
| CGGTGTCAACATCTCCAAGG | Pole3_R2.Br_26343 | |
| GCTTCCTTACAGCGTACAGG | Pole3_R2.Br_26344 | |
| CCTCCGGCATCCGTCCAAAG | Ppard_1 | Ppard |
| GCTTCATGCGGATTGTCCGG | Ppard_2 | |
| GGCCTCGGGCTTCCACTACG | Ppard_3 | |
| AGGGCTTGCTAACATCACAG | Ppargc1b_R2.Br_51224 | Ppargc1b |
| GGCCTTGACTACTGTCTGTG | Ppargc1b_R2.Br_51226 | |
| TCCTGTGCTAGGGAGTCTTG | Ppargc1b_R2.Br_51227 | |
| GTACACCTGAGGATTTGACC | Ppie_R2.Br_24042 | Ppie |
| GATGCTCCTACGGTCTGATG | Ppie_R2.Br_24043 | |
| CCACCCTGACACATGAACTG | Ppie_R2.Br_24045 | |
| GTAGGGACTGAAGACAACGG | Pprc1_R2.Br_56892 | Pprc1 |

| | | |
|---|---|---|
| TGACCTCGAGATCCCAGTCG | Pprc1_R2.Br_56893 | |
| ACACCCTCAGTAGACAAAGT | Pprc1_R2.Br_56895 | |
| AACACCTTGTACCAGCTCGG | Pqbp1_R2.Br_23754 | Pqbp1 |
| AGAGCGCAACTACGACAAAG | Pqbp1_R2.Br_23755 | |
| GTGGTGAGAGCCACGACACA | Pqbp1_R2.Br_23756 | |
| CTACCAGGTTCTGTTCTAGG | Prdm10_As_24889 | Prdm10 |
| ACTGGTCTACATCCACCCGG | Prdm10_As_24890 | |
| GATCTTTCAACAAATCAAGG | Prdm10_As_24891 | |
| GTACGTGGTCATTTCCCGGG | Prdm11_As_24885 | Prdm11 |
| GCGTGTCAGACACAAACACG | Prdm11_As_24886 | |
| GCTGCGGGTCTGGTACAGCG | Prdm11_As_24887 | |
| GCTGCTTTAGACTTTCCTTA | Prdm15_As_24870 | Prdm15 |
| CCTACTATCTGGGAAAAGGG | Prdm15_As_24871 | |
| AGTCCAGGAGAGTCGCCAAG | Prdm15_TF2.Br_50312 | |
| GGGTAGCGAGAAGTTCTGCG | Prdm16_B_24865 | Prdm16 |
| GCAACACATGATCGTCCACA | Prdm16_B_24866 | |
| GCTCATCACACTCATCACAG | Prdm16_B_24867 | |
| GCAGATGCTGACGGATACAG | Prdm16_B_24868 | |
| TCAGAACCCAGGCTGCAGCG | Prdm2_As_24861 | Prdm2 |
| ACACCGCTGGATAAATCCAA | Prdm2_As_24862 | |
| GATCCAATACAGATTCCCAT | Prdm2_As_24863 | |
| GATACGGATCTAGTGCTGAA | Prdm4_As_24857 | Prdm4 |
| GCTAAGGCCAACTTCATGGA | Prdm4_As_24858 | |
| GCAGAGGAGCTTACAATGGA | Prdm4_As_24859 | |
| ATTCGCTTCTCCCCAGCGAA | Prdm5_As_24853 | Prdm5 |
| GCTTGCACCAGAAGCCCACA | Prdm5_As_24855 | |
| GGAAAGCTCGATTCACACTG | Prdm5_TF2.Br_36145 | |
| TCTGACACTCAGATTACCAA | Prdm9_As_24841 | Prdm9 |
| ATAGTCCTGAAGAAGATACA | Prdm9_As_24842 | |
| GAAGCATCTGATCTACCAGT | Prdm9_As_24843 | |
| AGATCCTTACCTTCTCGTGA | Prkab1_As_24717 | Prkab1 |

| | | |
|---|---|---|
| GCATCAGTACAAGTTCTTCG | Prkab1_As_24718 | |
| GCACCTTGATCTCTTCGGAG | Prkab1_As_24719 | |
| GTACAAACTCTTTGTCCCCG | Prkab2_As_24713 | Prkab2 |
| CCTTGCCTCCTTCAGACCAG | Prkab2_As_24714 | |
| GCCATCCTGGATCTTCCAGA | Prkab2_As_24715 | |
| ACGAGTATACAGCATCGAAG | Prkag2_As_24697 | Prkag2 |
| TTAGCAGAAGACTCAGAAAG | Prkag2_As_24698 | |
| TGAACATCTCTCCAGATGCG | Prkag2_As_24699 | |
| ACTGAGGATTCAGTGTGGAG | Prkca_As_24673 | Prkca |
| GTTGCTTTCTGTCTTCTGAA | Prkca_As_24674 | |
| TTACCCGGCCAACGACTCCA | Prkca_As_24675 | |
| GAGTCTGTCGGAATATACCA | Prkcd_As_24665 | Prkcd |
| CCTGTTTAATGGCTCCACGA | Prkcd_As_24666 | |
| GTGTGTGCAGTATTTCCTGG | Prkcd_As_24667 | |
| GGGCCCTCCCAGCCTCCACG | Prkdc_As_24617 | Prkdc |
| ACAGGCTTCATCTCTCTGAA | Prkdc_As_24618 | |
| TGACTTCCTCGGAGGCCGAA | Prkdc_As_24619 | |
| GCAGTGGTGACCATCATCAA | Prmt1_B_24467 | Prmt1 |
| TGCCCACTTTGGCATCCACG | Prmt1_B_24468 | |
| GCATCTCTTCAAAGACAAGG | Prmt1_B_24469 | |
| CCTTGGCAGCAAACATGCAG | Prmt1_B_24470 | |
| TCAAAGGCAGGTATCCTCCG | Prmt10_As_24463 | Prmt10 |
| GCACGGGGACTGGAATACTG | Prmt10_As_24464 | |
| CCACCAGCCAGTTTGCTACG | Prmt10_As_24465 | |
| CCGTTCTGGATATCGGCACG | Prmt10_R2.Br_47304 | |
| AAGATGAGTGGCATCCCCGG | Prmt10_R2.Br_47305 | |
| TGGTGACCGAGTGACAATGG | Prmt10_R2.Br_47306 | |
| TCTCACAGCCATAGTCCACG | Prmt2_B_24459 | Prmt2 |
| TATCCTGAGACAAACCACGG | Prmt2_B_24460 | |
| TCTTATCCTGAGACAAACCA | Prmt2_B_24461 | |
| ACTGGAGGAATACGACCCGG | Prmt2_B_24462 | |

| | | |
|---|---|---|
| TCTTCATGCCCAGAATCCTA | Prmt3_As_24455 | Prmt3 |
| AGTTGAAGCACATGGAAGCG | Prmt3_As_24456 | |
| GCGTCTACTTCAGCTCCTAT | Prmt3_As_24457 | |
| GGTCCCTCCCGCTGGACACG | Prmt5_B_244511 | Prmt5 |
| TGATGGGCTCTCCAAGCCAG | Prmt5_B_244521 | |
| ATGAACTCCCTCTTGAAACG | Prmt5_B_24453 | |
| ACACACACAGAGGAGTACAG | Prmt5_B_24454 | |
| TGAGCTCCGTGCTCCACGCG | Prmt6_B_24447 | Prmt6 |
| GCAGCTCCACGGTCTCCACC | Prmt6_B_24448 | |
| CTAAGCGGTAGGCTTCGGTG | Prmt6_B_24449 | |
| CGCTCGCTCTTGGTCCTCCG | Prmt6_B_244501 | |
| TGGTGGGCTACCAGGCACCG | Prmt7_B_24443 | Prmt7 |
| TGAACACTATGATTACCACC | Prmt7_B_24444 | |
| GTAAACCTAGGAAGACTGCG | Prmt7_B_24445 | |
| TAGAATATAAAATACTACCA | Prmt7_B_24446 | |
| GTACTGGAAGACATTGCCAG | Prr14_As_24232 | Prr14 |
| CCAGGAGCCCGAAAAGACCG | Prr14_As_24233 | |
| TGACGATGTTGGAATCACCG | Prr14_As_24234 | |
| AGAGAGGACGACCTGCAGGT | Psip1_R2.Br_47200 | Psip1 |
| GAAAAATTTAGCTAAACCAG | Psip1_R2.Br_47201 | |
| ACTCCAAAAGCTGCCAGGCG | Psip1_R2.Br_47202 | |
| ACCCGAGAGTCATATTCAGT | Psmc3_R2.Br_13882 | Psmc3 |
| ACAGCAAGATGGCATTGGGG | Psmc3_R2.Br_13884 | |
| CGTTGACCCCAATGACCAGG | Psmc3_R2.Br_13885 | |
| AGGAACGTGAGCGCACGGCG | Ptov1_R2.Br_45513 | Ptov1 |
| CCTCGCAAGGAGAAATGTGG | Ptov1_R2.Br_45514 | |
| TCGGAACAGGGGACCCAGTG | Ptov1_R2.Br_45515 | |
| GCCAACGGATGACAGAGAAA | Ptpn2_As_23339 | Ptpn2 |
| CTAAGACTCACCAAGTATAA | Ptpn2_As_23340 | |
| CCATGACTATCCTCATAGAG | Ptpn2_As_23341 | |
| CATCCTTAGATGCCAAACGG | Pttg1_R2.Br_21750 | Pttg1 |

| | | |
|---|---|---|
| TTGAACACTTTGCCGACTCG | Pttg1_R2.Br_21751 | |
| CAAAACAGCCGACCTTGACT | Pttg1_R2.Br_21753 | |
| GCTCATCGACGACTATGGAG | Pura_R2.Br_14194 | Pura |
| ATCCGCCAGACAGTCAACCG | Pura_R2.Br_14195 | |
| GTGGAGTTCCGCGACTACCT | Pura_R2.Br_14196 | |
| AAGCTCATCGACGACTACGG | Purb_R2.Br_14198 | Purb |
| CGCCGCGGTTCACCGTCTGG | Purb_R2.Br_14200 | |
| GGAACTCGCTCTTGAGCGCG | Purb_R2.Br_14201 | |
| ACCCAACAGAGACTCGCCAA | Pwp1_R2.Br_47464 | Pwp1 |
| ATGCAATGTGTGCAGATCGG | Pwp1_R2.Br_47465 | |
| TCCTCCAGGGACTCCCTAGG | Pwp1_R2.Br_47466 | |
| GGAGGGCCCAATATACAACT | Pygo1_As_23022 | Pygo1 |
| TATGGATTAGATGAAGGCAG | Pygo1_As_23023 | |
| ATTGGCAGCCACTAGATGGT | Pygo1_As_23024 | |
| GATGTGAATATGCAGGACCC | Pygo2_R2.Br_33602 | Pygo2 |
| GGAAAGTTCATGTTGCCAGG | Pygo2_R2.Br_33603 | |
| TTTCGGAGCCCCTAAAGTGG | Pygo2_R2.Br_33604 | |
| GACAACTCTCGTCCCAAACG | Rad21_R1.Br_14354 | Rad21 |
| ATGCTTCATTACAGTCTGCG | Rad21_R1.Br_14355 | |
| TAATGCCATTACTTTACCTG | Rad21_R1.Br_14357 | |
| GGTTCCAATGGGTTTCACCA | Rad51_As_22534 | Rad51 |
| TTATCTGTATGATCTCTGAC | Rad51_As_22535 | |
| GCAGCAAAATTGGTTCCAAT | Rad51_As_22536 | |
| GCACTCATCGGCCTATCATG | Rai1_B_22458 | Rai1 |
| GGACAGCAAGAGCTGCTGCA | Rai1_B_22459 | |
| GTTGGGGACGGGTTAAGCAG | Rai1_B_22460 | |
| GCTGCAGACAAGTACCACCG | Rai1_B_22461 | |
| AAGCATGGCTTATAGACCCG | Rarg_1 | Rarg |
| CCTTAGTGCTGATGCCCCGG | Rarg_2 | |
| TGGGCAAGTACACCACGGTG | Rarg_3 | |
| GCTCTGCGCGGGGCTTTGGG | Rb1_As_22110 | Rb1 |

| | | |
|---|---|---|
| CAAAGCCCCGCGCAGAGCCG | Rb1_As_22111 | |
| AGAAATCGATACCAGTACCA | Rb1_As_22112 | |
| GGAAGAACGGGTGATCAACG | Rbbp4_As_22098 | Rbbp4 |
| GGCCCAGCTTAACTGCCCAG | Rbbp4_As_22099 | |
| GGAAGGTACCTGGTTTAGAA | Rbbp4_As_22100 | |
| GGATGGGGAGTACATAGTGG | Rbbp5_As_22094 | Rbbp5 |
| CTTTGGACAGAACTACCCAG | Rbbp5_As_22095 | |
| CGATGACTCGGATTTGAACG | Rbbp5_As_22096 | |
| TGATGAGCAGAACCATCTGG | Rbbp7_As_22086 | Rbbp7 |
| ACTGGGCCACTGAAGAGCAT | Rbbp7_As_22087 | |
| TTTCTCCTAAACAGACCAGA | Rbbp7_As_22088 | |
| TCCTGCACCAGTTAACTCTG | Rbbp8_R2.Br_56532 | Rbbp8 |
| AGGCCGTAAAGCCTTGAGTG | Rbbp8_R2.Br_56533 | |
| CAGCAACAGTAGTATTCATG | Rbbp8_R2.Br_56535 | |
| GCTCACCCCGTACTTTCCCG | Rbl1_R2.Br_14499 | Rbl1 |
| CTACCCCACTTACTGGACGG | Rbl1_R2.Br_14500 | |
| GTGGGAGCCTTCGACAGGGT | Rbl1_R2.Br_14501 | |
| GCTGTACCTGTCATACAATG | Rbl2_R2.Br_14502 | Rbl2 |
| TGAGCGCGTATTCCTTGGTG | Rbl2_R2.Br_14503 | |
| GTACGTTCTCGGAAATGTGG | Rbl2_R2.Br_14504 | |
| GGCCTGACATTCGATACCTG | Rbx1_R2.Br_24790 | Rbx1 |
| ATGGATGTGGATACCCCCAG | Rbx1_R2.Br_24791 | |
| CCACAATGTCCCAGGCCCAG | Rbx1_R2.Br_24792 | |
| CCTGAATTATTTGCCAACCG | Rcc1_R2.Br_46800 | Rcc1 |
| AAGTGGTGCAAGTGTCAGCG | Rcc1_R2.Br_46802 | |
| ACTGTGTGTCTGAGCCAAAG | Rcc1_R2.Br_46803 | |
| GCTATCGCCAAAGAGAAGCA | Rcor1_As_21750 | Rcor1 |
| GGTGGGACCCCAGTACCAAG | Rcor1_As_21751 | |
| GGGACCCCAGTACCAAGCGG | Rcor1_As_21752 | |
| GTTTGGCATCTGATACACAG | Rcor2_As_21746 | Rcor2 |
| GATGAACTAGAAGAGGGACG | Rcor2_As_21747 | |

| | | |
|---|---|---|
| GCACTTGCCATGGAACCCAA | Rcor2_As_21748 | |
| AGCAGCGACGACGAGCACGG | Rcor3_As_21742 | Rcor3 |
| GCAGATGTTGGGATGAGAGT | Rcor3_As_21743 | |
| AGAAAGGGCCCGAGTTACTG | Rcor3_As_21744 | |
| GATTCCGCTATAAATGCGAG | Rela_1 | Rela |
| TAAATGCGAGGGGCGCTCAG | Rela_2 | |
| TATCAAAAATCGGATGTGAG | Rela_3 | |
| GCAGAGGTGAAACACACAGA | Rest_As_21510 | Rest |
| GCTGTGGCTACAATACCAAC | Rest_As_21511 | |
| TCGTTCACAGGCTCTTCGGA | Rest_As_21512 | |
| CTCCTGTTACCCGCTCCGCA | Ring1_As_20916 | Ring1 |
| ACAGAATGCCAGCAAAACGT | Ring1_As_20917 | |
| CCTGCTCCCAAGCGACCCCG | Ring1_As_20918 | |
| GTAGGGTTACAGGCTCTAGG | Rnf168_As_20568 | Rnf168 |
| ATATGAAGAGGAGATAAGCA | Rnf168_As_20569 | |
| GAGATGGAAGAACAGCTGAG | Rnf168_As_20570 | |
| CAGCAGATAGAAAATGGTAG | Rnf2_As_20520 | Rnf2 |
| GTATGAAGCGCATCAGGAAA | Rnf2_As_20521 | |
| ACAAAGAGTGTCCTACCTGT | Rnf2_As_20522 | |
| TGAAGATTCAGGGACCACCG | Rnf20_As_20516 | Rnf20 |
| GCTGGTGGTTATGATTCTGG | Rnf20_As_20517 | |
| TCACAATCTGGGACACAGCA | Rnf20_As_20518 | |
| GGAATTGGCCAATAGCCGAA | Rnf40_As_20440 | Rnf40 |
| CCAGTGCTACGAGAACCAGA | Rnf40_As_20441 | |
| TGATCCCTCGACCAGACCCA | Rnf40_As_20442 | |
| GTTCTGAAGCAGAATCCTGA | Rnf8_As_20412 | Rnf8 |
| CGAGCTGCCCCGAGACCAAG | Rnf8_As_20413 | |
| TCATTCTTCGGGGCAAGGCA | Rnf8_As_20414 | |
| GGAACCGGCAGAAAAATAAG | Rrp8_As_19671 | Rrp8 |
| GTTTGGTCAACACTGTCCAG | Rrp8_As_19672 | |
| GCAGAGACCCCACCAAGAGG | Rrp8_As_19673 | |

| | | |
|---|---|---|
| AGAAGCAAAGGAATGCAGAG | Rsf1_As_19635 | Rsf1 |
| CCTCATGTATTGGTACCAGT | Rsf1_As_19636 | |
| ATACCAGAAGAAGTTGTAAG | Rsf1_As_19637 | |
| AGAACTGAGAAATGCTACCG | Runx1_1 | Runx1 |
| TAGAGCCATCAAAATCACAG | Runx1_2 | |
| TGCGCACTAGCTCGCCAGGG | Runx1_3 | |
| CAGTTCCCAATGGTACCCGG | Runx2_1 | Runx2 |
| CTGTCACTTTAATAGCTCTG | Runx2_2 | |
| GAATGCGCCCTAAATCACTG | Runx2_3 | |
| TGTGAGACAGAGAACCCCAT | Ruvbl1_As_19411 | Ruvbl1 |
| CCAGCCCCAGACCCTTCACG | Ruvbl1_As_19412 | |
| GCTGATGGAGAACTTCCGCA | Ruvbl1_As_19413 | |
| GTAGATGGTCTCCATCTCCG | Ruvbl2_As_19407 | Ruvbl2 |
| GCGCTCACTCCCACATCCGG | Ruvbl2_As_19408 | |
| GAGAAGGGAAGATTGCAGGG | Ruvbl2_As_19409 | |
| ACGCCATTGAGGCCTAGTGG | Rxra_1 | Rxra |
| AGGACGCCATTGAGGCCTAG | Rxra_2 | |
| CAAGACTGAGACATACGTGG | Rxra_3 | |
| ATTGTGGGGTCGCGTCCCGG | Rxrb_1 | Rxrb |
| GTTCAGGAGGAGCGTCAACG | Rxrb_2 | |
| TGATAACAAAGACTGTACAG | Rxrb_3 | |
| AGAAAAGCCAGAGAAAGACA | Rybp_1 | Rybp |
| AGTTACTGCCAACTGCTGTG | Rybp_2 | |
| TGTATGCTGTTAGCTTCACT | Rybp_3 | |
| GATGCTGCATTTAAAGGCTT | Rybp_As_19355 | |
| AATGCAGCATCTGCGATGTG | Rybp_As_19356 | |
| CGTCTTCCAGTACCTGGTGG | Rybp_As_19357 | |
| GGATGCTGGATAGATAAAGT | Sap130_R1.Br_68136 | Sap130 |
| GTAATCAGGCTGGATCCGAG | Sap130_R1.Br_68138 | |
| GGAAGCTCCAATGTGAAGTG | Sap130_R1.Br_68139 | |
| GAAGCCGATCGACCGCGAGA | Sap18_R1.Br_15222 | Sap18 |

| | | |
|---|---|---|
| GCGAGCTGCAGATCTACACC | Sap18_R1.Br_15223 | |
| GAATGGACGAGTTCTCCCGC | Sap18_R1.Br_15224 | |
| TGGCCGATGTATGATAACTG | Sap25_R1.Br_77071 | Sap25 |
| ACCAGGCTAATGCAGGACCT | Sap25_R1.Br_77072 | |
| GCAGTGGAGACATCCTTGAG | Sap25_R1.Br_77073 | |
| GGAGGAGATGAGCCGAGGCG | Sap30_B_19094 | Sap30 |
| ACTCCGGAGGAGATGAGCCG | Sap30_B_19095 | |
| GCAAACAACAGAGTTGCCCA | Sap30_B_19096 | |
| GCAACTCTGTTGTTTGCGGG | Sap30_B_19097 | |
| CTACGAGCGGAAGATAAAGG | Sap30bp_R1.Br_25510 | Sap30bp |
| AGATGACTTCTCTCGCCCAG | Sap30bp_R1.Br_25511 | |
| GGAGTCTTCAGACCAGCCGT | Sap30bp_R1.Br_25512 | |
| AAGAGGAAGGCAAGTGACGA | Sap30l_R1.Br_21942 | Sap30l |
| CCAGAGCTGCTGCCTCATCG | Sap30l_R1.Br_21943 | |
| CAGCTGGAACAGATCAACCT | Sap30l_R1.Br_21944 | |
| TCTTTACCTTTGGTGCCAGG | Scmh1_As_18731 | Scmh1 |
| GGAAACTGTGAGAAGAATGG | Scmh1_As_18732 | |
| GGTGACCCCATGATGGAGCA | Scmh1_As_18733 | |
| ACGAGGGAGAAGACCAGCCG | Scml4_As_18723 | Scml4 |
| GGCCTCCACTTCACTCCACA | Scml4_As_18724 | |
| AGCAGAATAGAGGTTATGGA | Scml4_As_18725 | |
| GCATCAAATACACAGTCCGA | Senp1_As_18171 | Senp1 |
| GAGGTGACCTTAGTGAACCA | Senp1_As_18172 | |
| GCCACAGTCAGGCTTTCCAG | Senp1_As_18173 | |
| GCTAGGTGCTCTCATGGCAG | Senp3_As_18163 | Senp3 |
| GCACCTAGCAGACCCAGCCG | Senp3_As_18164 | |
| CCACCACCGGACTTGAGCCG | Senp3_As_18165 | |
| ACATTTGTCAACCATCCACA | Set_R2.Br_24130 | Set |
| GAATTTCATCTGAACGAGAG | Set_R2.Br_24131 | |
| TCTCCCGGCCTGCCGAAGGG | Set_R2.Br_24132 | |
| TCTACTCTCAGGCATACGAG | Setbp1_B_17792 | Setbp1 |

| | | |
|---|---|---|
| ACACCGGTCAAAAAGAAGCG | Setbp1_B_17793 | |
| GCAACCATAAGAGGAAAAAG | Setbp1_B_17794 | |
| CCTCCATCTTGATCTCCCAA | Setbp1_B_17795 | |
| CCTTAAACTTAGGAACCGGG | Setd1a_As_17788 | Setd1a |
| GCCTTGAGTGAGAAGTTCCA | Setd1a_As_17789 | |
| CCTATGCTCTGTACACACAA | Setd1a_As_17790 | |
| TTATGTCCACAACTCAGCGG | Setd1b_As_17784 | Setd1b |
| CATGGGCAACATTATCCACG | Setd1b_As_17785 | |
| ACAGATGTCCAGCAACCGCC | Setd1b_As_17786 | |
| GCATTCGCTTAATATCCCGG | Setd2_R2.Br_60616 | Setd2 |
| TTGCTTATGATCGAATCCAA | Setd2_R2.Br_60617 | |
| TGCTCATGCTCAGAGTGACG | Setd2_R2.Br_60618 | |
| GCAGAGCCACACACTCACAG | Setd3_As_17776 | Setd3 |
| CTCTTCTTCGAAGTAGAGAG | Setd3_As_17777 | |
| GCAAAACCAAATTCCTACAG | Setd3_As_17778 | |
| CTGTGATTCGAAGCTCCCTA | Setd4_B_17772 | Setd4 |
| TTAGTCTCAGAAAAGCATGC | Setd4_B_17773 | |
| TGAGAGCTGTCTGCTCACCA | Setd4_B_17774 | |
| GTCCAGGTAAGACTTCCAGA | Setd4_B_17775 | |
| TGAGACACATGATTGCAGAT | Setd5_R2.Br_38958 | Setd5 |
| AAAGCGAATGTAGTGTTTGG | Setd5_R2.Br_38959 | |
| ATGATTGCAGTTAAGCAGGG | Setd5_R2.Br_38960 | |
| CTCGGCTCTCACCAAAACAT | Setd6_B_17764 | Setd6 |
| GGAGCTGAGTCCTAAGGTGA | Setd6_B_17765 | |
| GAGCTGCTGATAGAGTTCCA | Setd6_B_17766 | |
| TGACCTATAAGCCATCACGA | Setd6_B_17767 | |
| CCTCTGGTCAGGGTACACAT | Setd7_B_17760 | Setd7 |
| GACGATGACGGATTACCACA | Setd7_B_17761 | |
| TGAGGTGGTGGAGGAAGCCG | Setd7_B_17762 | |
| GTCCACGTAATATCCCTCCA | Setd7_B_17763 | |
| GAACTCGGTTGCCCATCATG | Setd8_1 | Setd8 |

| | | |
|---|---|---|
| TCACAGATTTCTACCCTGTG | Setd8_2 | |
| CCGCCGCCGCCTCCACCGCG | Setd8_B_17756 | |
| CTAACAAATACCTCTAGCCA | Setd8_B_17757 | |
| GAAGATGTGCAAGCCCCGCG | Setd8_B_17758 | |
| TCAAGGGCAAACAGGCCCCT | Setd8_B_17759 | |
| GCAAGAAGAGGACTAAGACA | Setdb1_B_17752 | Setdb1 |
| ACAGAAGTGGAACTAATAGA | Setdb1_B_17753 | |
| GCTGTTTGTGGGCAGTCGAG | Setdb1_B_17754 | |
| GTATCATGACAGTAGCTCTG | Setdb1_B_17755 | |
| TTTCGTAGATTCCGTCCACA | Setdb2_B_17748 | Setdb2 |
| GTATTTCTATACTTAAACCG | Setdb2_B_17749 | |
| GTTGACAGCAAAGAATGCCA | Setdb2_B_17750 | |
| TTAAGTATAGAAATACAGTA | Setdb2_B_17751 | |
| ATAGTCCAATTACGTGACCA | Setx_R2.Br_68228 | Setx |
| TCTGTATCACTACTAGACGC | Setx_R2.Br_68229 | |
| CTCACAACTCCGGACAAATG | Setx_R2.Br_68230 | |
| TTTGTATTCATCTTCCCGAT | Sf3b1_As_17708 | Sf3b1 |
| GCATACCCCATCTTTAAGAT | Sf3b1_As_17709 | |
| GCAACTCCTACACCTTTGGG | Sf3b1_As_17710 | |
| GGAGGGTCAGATGGTCCAAG | Sf3b3_As_17700 | Sf3b3 |
| GTACAACCTGACCTTACAGC | Sf3b3_As_17701 | |
| AAGACTCTCAGGGAGGACCG | Sf3b3_As_17702 | |
| GCTTCGTTATGAAGGCTATG | Sfmbt1_As_17680 | Sfmbt1 |
| AAGACCCTCGAAGCTCCAGA | Sfmbt1_As_17681 | |
| GTTGCAGGAGAAATCCCAAA | Sfmbt1_As_17682 | |
| TCAGGACCGGAATCAACGCA | Shprh_B_17230 | Shprh |
| ACCATTCCACCGCTACCAAG | Shprh_B_17231 | |
| TTGTAGCAAGGGAATCCGAG | Shprh_B_17232 | |
| TGTGAAGTGCACTCTCAGGA | Shprh_B_17233 | |
| TATGCCTCTTGTAAACGACT | Sin3a_R1.Br_15842 | Sin3a |
| TAAAGCCCATGATCAGATCA | Sin3a_R1.Br_15843 | |

| | | |
|---|---|---|
| AACGTGGGGATGCATATGGT | Sin3a_R1.Br_15844 | |
| TGAGGACCGACAGATCCTAG | Sin3b_R1.Br_15846 | Sin3b |
| CTCCAAGAAGACGCCCTACG | Sin3b_R1.Br_15847 | |
| ATGTCTATACGGTATCCAAG | Sin3b_R1.Br_15848 | |
| GCCATTAGTGAGGAGTCCAT | Sirt1_R2.Br_45800 | Sirt1 |
| CGGTATCTATGCTCGCCTTG | Sirt1_R2.Br_45801 | |
| AGCTGGATGATATGACGCTG | Sirt1_R2.Br_45802 | |
| CCTCCAGAACATAGACACGC | Sirt2_R2.Br_26958 | Sirt2 |
| GTAGCGTGTCACTCCTTCGA | Sirt2_R2.Br_26959 | |
| ACCGAAAAACACGATATCTG | Sirt2_R2.Br_26960 | |
| TAGCTGTTACAAAGGTCCCG | Sirt3_B_17070 | Sirt3 |
| TCACCCATATGTCTTCCCCT | Sirt3_B_17071 | |
| CTTCTCTCTGCAGATCCCCA | Sirt3_B_17072 | |
| GCCGGCATCAGCACACCCAG | Sirt3_B_17073 | |
| GGGCGGCACAAATAACCCCG | Sirt4_R2.Br_41578 | Sirt4 |
| CTGGTTGGTGACTCAGAACG | Sirt4_R2.Br_41579 | |
| ACTCCTCGTGATGACAGGCG | Sirt4_R2.Br_41580 | |
| CCTCACCGTGGATTTCCAGA | Sirt5_B_17062 | Sirt5 |
| ACTTGAACTTGGACGAGCCA | Sirt5_B_17063 | |
| GCACATAGCCATCATCTCGG | Sirt5_B_17064 | |
| GTTTCGAGCAAAGGCCTGAG | Sirt5_B_17065 | |
| TGTTCACCGACCTGAAGTCG | Sirt6_R2.Br_21930 | Sirt6 |
| ACAGATCTTCGACCCACCAG | Sirt6_R2.Br_21932 | |
| GCACGGAAACATGTTTGTAG | Sirt6_R2.Br_21933 | |
| CAGCAGCTTTGCGCTCCGAG | Sirt7_B_17054 | Sirt7 |
| GATCTGGTGACCGAGCTGCA | Sirt7_B_17055 | |
| GCTTCTATCCCAGATTATCG | Sirt7_B_17056 | |
| GTATGGACACTGCTTCAGAA | Sirt7_B_17057 | |
| GAGTGGGTGTAGTCATCCAG | Smad2_1 | Smad2 |
| GCCGTCTTCAGGTTTCACAC | Smad2_2 | |
| TCTTCACTGATATATCCAGG | Smad2_3 | |

| | | |
|---|---|---|
| CCATGGCCCGTAATTCATGG | Smad3_1 | Smad3 |
| CTACCTGGAATATTGCTCTG | Smad3_2 | |
| GGTTGACAGACTGAGCTAGG | Smad3_3 | |
| CAGGTGCCTTAGTGACCACG | Smad4_1 | Smad4 |
| GGTGGCGTTAGACTCTGCCG | Smad4_2 | |
| TGTCACCATACAGAGAACAT | Smad4_3 | |
| CCGACGCTGGATGCGCCCGG | Smad6_1 | Smad6 |
| CTACCGAGCGGACTTCGGAG | Smad6_2 | |
| GTAGGGGTTGCAACACACCG | Smad6_3 | |
| GATGAAGTACTCACAGGAAG | Smarca1_As_15201 | Smarca1 |
| CTTCTTCTCGCCCTTCTCAG | Smarca1_As_15202 | |
| TCTGGCATCCGAAACTGCCA | Smarca1_As_15203 | |
| GGTGACAACTTCTCAGCCGG | Smarca2_As_15197 | Smarca2 |
| GTGACAACTTCTCAGCCGGA | Smarca2_As_15198 | |
| TTTCCACATCTTCCTTCACG | Smarca2_As_15199 | |
| ACCCCCATCCAGAAGCCCCG | Smarca4_As_15193 | Smarca4 |
| GCTCTTGTAGGATCTCCACA | Smarca4_As_15194 | |
| GCATGTTCAGAGCCGCCGAG | Smarca4_As_15195 | |
| CCTCCATCTCGGTGTCCGCG | Smarca5_As_15189 | Smarca5 |
| GAGAATTCAAGACTACAAAT | Smarca5_As_15190 | |
| CAGTTACCGACACCGTAGAA | Smarca5_As_15191 | |
| TCACTTACACGGTCCCCTCG | Smarcad1_As_15185 | Smarcad1 |
| TTTGATATGGAATTGCAAAA | Smarcad1_As_15186 | |
| CAGTCTGTAAAACAGCCGCG | Smarcad1_As_15187 | |
| GTCCGCTTCGAGGTTGACAT | Smarcal1_As_15181 | Smarcal1 |
| CTCTCTCCTGAATCTCCTAG | Smarcal1_As_15182 | |
| GGTTGGTTAGAGGCTCCTGG | Smarcal1_As_15183 | |
| GACACTCCTGAAAGCCTCAG | Smarcb1_As_15177 | Smarcb1 |
| GTTCTACATGATCGGCTCCG | Smarcb1_As_15178 | |
| AACTACCTGCGTATGTTCCG | Smarcb1_As_15179 | |
| TATCCTTCCTCACAAGAGGA | Smarcc1_As_15173 | Smarcc1 |

| | | |
|---|---|---|
| CCTCTTCCTAGAGTTGGCAG | Smarcc1_As_15174 | |
| TCAGTATTCATGTCATTGAA | Smarcc1_As_15175 | |
| GCTCGGCAAGAACTACAAGA | Smarcc2_As_15169 | Smarcc2 |
| CTCCCACTCCTGAGAAACCG | Smarcc2_As_15170 | |
| CCCCAACGTGAAGTACTACG | Smarcc2_As_15171 | |
| TTCCAGTCTGTGGCTCCGAG | Smarcd1_As_15165 | Smarcd1 |
| GGTCCCATGGAAGGTCCCTG | Smarcd1_As_15166 | |
| GGTAGCCGAATGACACCTCA | Smarcd1_As_15167 | |
| CCATTCTTACCTCCACCAGA | Smarcd2_As_15161 | Smarcd2 |
| GAGAGAGAGAGAGAGATGGA | Smarcd2_As_15162 | |
| GAGATAATGCGGGAACTGCG | Smarcd2_As_15163 | |
| TCTGTCTTACCTCAACAAGG | Smarcd3_As_15157 | Smarcd3 |
| GTTTCCTCTCAAATGCTAGG | Smarcd3_As_15158 | |
| CCCGTACATGGGCAGCCCCG | Smarcd3_As_15159 | |
| AGCAAATGCCCAGCACACCA | Smarce1_As_15153 | Smarce1 |
| AATATAGGCAAGGTACGCAG | Smarce1_As_15154 | |
| TATTAATGCAAAAAGTCGTG | Smarce1_As_15155 | |
| TGAAGAAATTAGTCGCTCTG | Smc1a_R1.Br_19902 | Smc1a |
| CCAGGTGATGGAGCAATTAG | Smc1a_R1.Br_19903 | |
| CTCTTGATATCATTAATGCG | Smc1a_R1.Br_19904 | |
| ACTAAAGAGTAAGCAAGCAG | Smc2_B_15141 | Smc2 |
| CCTGCATGATGAGAAAGTGA | Smc2_B_15142 | |
| TCATGAGCTTCAAATCCCAA | Smc2_B_15143 | |
| GATTCAAGTCCTATGCACAG | Smc2_B_15144 | |
| GATCTACAATTGTTTGGTCT | Smc3_2 | Smc3 |
| TCTGCTAAGCGAGAAACGAG | Smc3_3 | |
| GTAAAAGAAAAAGAAGAGCG | Smc3_B_15137 | |
| AAGAAGAGCGAGGAATTGCG | Smc3_B_15138 | |
| ATAGATCAACCAAATGGCAA | Smc3_B_15139 | |
| ACAAAAGCAGAAATAACACG | Smc3_B_15140 | |
| AGAAGTGCTATCTCTGTAGG | Smc4_1 | Smc4 |

| | | |
|---|---|---|
| ATTGCTATGATGAAACCCAA | Smc4_2 | |
| TTAACTAGATCAAATAACCG | Smc4_3 | |
| GGGGCTTCAGAAAGAAAAGG | Smc4_B_15133 | |
| GAAGAGATGGAGCAACCGGC | Smc4_B_15134 | |
| TTATCCAGCTCCTCACCTGG | Smc4_B_15135 | |
| GCCTTAAGCAGGAAACACAG | Smc4_B_15136 | |
| TTAACTCCTCATCATCAAAG | Smc5_3 | Smc5 |
| GGCAGATTCCTATGACACGG | Smc5_R1.Br_56812 | |
| ACGCCTCAGATCATTTGTGA | Smc5_R1.Br_56813 | |
| CCAATGATCATGTTCAGATG | Smc5_R1.Br_56815 | |
| TGTGCAGCAACACATTAGCG | Smc6_R1.Br_30310 | Smc6 |
| AACTCAGACACAATTATCTG | Smc6_R1.Br_30312 | |
| ACCAAGACCAACTATGAGTG | Smc6_R1.Br_30313 | |
| GTTTCTCTATCATATAAGAA | Smchd1_R2.Br_15121 | Smchd1 |
| TATCCGAGAAAGCAGCTCGA | Smchd1_R2.Br_15122 | |
| TTTCTCCCTCAACATGGCAG | Smchd1_R2.Br_15123 | |
| CTACGCCTGCGTGCTCACCG | Smyd2_B_14869 | Smyd2 |
| AGAAGAGGAGGTCGCCCACG | Smyd2_B_14870 | |
| AGAGCTCTCCCATTTGGGAT | Smyd2_B_14871 | |
| GTTCTCCCCCAAAACAACCA | Smyd2_B_14872 | |
| AGAAGTCTCACGGAGTCGGG | Smyd3_B_14865 | Smyd3 |
| TGCACACAGTGTAAGCCAAG | Smyd3_B_14866 | |
| TGTGGCTCCACTGCGCCCCG | Smyd3_B_14867 | |
| GGTTCCGTACGTACCCCAGT | Smyd3_B_14868 | |
| ACTTTCTAGTAGTTACTCGG | Smyd4_R1.Br_69680 | Smyd4 |
| ATGATGCTCTCGTTAGACCC | Smyd4_R1.Br_69681 | |
| ACTTTGGCCACAGTACCTTG | Smyd4_R1.Br_69682 | |
| AATGCACTTTATCAGTACAG | Smyd5_R1.Br_59192 | Smyd5 |
| GAGGATAGTGAACACTCCTA | Smyd5_R1.Br_59193 | |
| TGTACTGCAGTGCAGAATGT | Smyd5_R1.Br_59194 | |
| ACTTCATCCAAGGATCACAG | Snai2_1 | Snai2 |

| | | |
|---|---|---|
| GAAATGTTTCTTGACCAGGA | Snai2_2 | |
| GTAAGAGGAGAAAGGCCACT | Snai2_As_14849 | |
| GTAACTTTCATAGAGATATG | Snai2_As_14850 | |
| TTCTTGACCAGGAAGGAGCG | Snai2_As_14851 | |
| ATAGAAAACAAGACTCCCCA | Snd1_B_14781 | Snd1 |
| TCAGGAGGAGGACCACCCCG | Snd1_B_14782 | |
| ATTGTCCGAGGGCAGCCCCG | Snd1_B_14783 | |
| GGTGCGCCATAATTGTCCGA | Snd1_B_14784 | |
| AAGGCTGGGATCATGACGTA | Sox13_1 | Sox13 |
| ACCGCAACTTACAGGAGGGT | Sox13_2 | |
| AGGAGCAGGCCCGTCTGAGC | Sox13_3 | |
| ACAACCCCAGTGGATCACTG | Sox4_1 | Sox4 |
| CAAGATTCCGTTCATCCAGG | Sox4_2 | |
| CCACGGCCGTCTACAAGGTG | Sox4_3 | |
| GGATCTGAAGAAGGAGAGCG | Sox9_1 | Sox9 |
| GTACCCGCATCTGCACAACG | Sox9_2 | |
| GTTCACCGATGTCCACGTCG | Sox9_3 | |
| GATGTTGGTGGCAATAATGG | Sp1_As_14305 | Sp1 |
| ACAGCCGCACAACTTTCACA | Sp1_As_14306 | |
| TCAAGGCCAGACGCCCCAGA | Sp1_As_14307 | |
| TGGAAGATGGAAGGGAGCGA | Sp100_B_14301 | Sp100 |
| TCTACCAGTTGCACAAAACA | Sp100_B_14302 | |
| GTTTCCAAGGAATTGATGGA | Sp100_B_14303 | |
| GCTAGATTCCTGTAGATCGC | Sp100_B_14304 | |
| GTGACGTGTGGTAACTTGAA | Sp140_As_14293 | Sp140 |
| TAGAATGTGTGGGCACCAGG | Sp140_As_14294 | |
| TGTTGGGGAACATATGACAC | Sp140_As_14295 | |
| CAGAGCCCCGAGAAACCGCG | Spen_As_13958 | Spen |
| GCACCTTGAGAGGAAGAGCG | Spen_As_13959 | |
| GGACTGGGAGAACTAACACA | Spen_As_13960 | |
| ACAGCAGCTCTATCGCCACA | Spi1_B_139181 | Spi1 |

| | | |
|---|---|---|
| GATGTTACAGGCGTGCAAAA | Spi1_B_139191 | |
| CTTTCTCTCTCGGCAGCCAT | Spi1_B_139201 | |
| ATAGAGCTGCTGTAGCTGCG | Spi1_B_13921 | |
| GCACGTCCTCGATACTCCCA | Spi1_R1.Br_156141 | |
| TGATAAGGGAAGCACATCCG | Spi1_R1.Br_156151 | |
| GTTGTTGTGGACATGGTGTG | Spi1_R1.Br_156171 | |
| CTTGATGGAACGGTTCAATG | Srcap_As_77988 | Srcap |
| TCATCCAGTCGAGAAACCGG | Srcap_As_77989 | |
| AGGAGTCAATCCATCACGAG | Srcap_As_77990 | |
| TCCTTGGTTCGGATATCCGG | Srsf1_As_13391 | Srsf1 |
| GTAGGTTACCCACGTAGATG | Srsf1_As_13392 | |
| CGACCTGAAGAACCGCCGCG | Srsf1_As_13393 | |
| GTCTTACTGCGAGCACTCCG | Ss18_As_13339 | Ss18 |
| ATTGGCATAGTCTGTGTGGG | Ss18_As_13340 | |
| GCTCACCTTCTGGATGGCGG | Ss18_As_13341 | |
| ACTGCCTGACAAGCTCATGG | Ss18l1_As_13335 | Ss18l1 |
| CATGCAGTCCCTGCTTCCCG | Ss18l1_As_13336 | |
| GTATCGCTGAGGCTGCCCTG | Ss18l1_As_13337 | |
| GAAGAAACCTCACCTCCACA | Ssrp1_As_13259 | Ssrp1 |
| CGGATATCGTATCGACCGCG | Ssrp1_As_13260 | |
| GGGCATGTCTACAAGTACGA | Ssrp1_As_13261 | |
| CCCCCAACAGAACGGCGACG | Stag1_R1.Br_16730 | Stag1 |
| CAGTTGGTGAAAATTACTGA | Stag1_R1.Br_16731 | |
| CAGACATGACCCACAAGCAG | Stag1_R1.Br_16732 | |
| TCTGGCTCAAACCGAATGAA | Stag2_B_13069 | Stag2 |
| AATGTCTTACTGCTTTACAA | Stag2_B_13070 | |
| GCTGAATGTCATCCTCCCGT | Stag2_B_13071 | |
| GGATGGGCTGATGAAAAGAA | Stag2_B_13072 | |
| CATAATACTTGACTACGTGA | Stk4_As_12826 | Stk4 |
| CATTCGGCTACGGAACAAGA | Stk4_As_12827 | |
| GGATGGATATCAGCATATGG | Stk4_As_12828 | |

| | | |
|---|---|---|
| CCACCACTCTGAATGATAGG | Supt16_As_12422 | Supt16 |
| GCTATCAAAGAAAGCAAGAG | Supt16_As_12423 | |
| ACTGGCAGCACAGCTCAACG | Supt16_As_12424 | |
| TTAGCTGTGGAGAAAGATGG | Suv39h1_B_12350 | Suv39h1 |
| ATACTCATTGATATAGACAA | Suv39h1_B_12351 | |
| CAGAAAACACCTGGGAGCCA | Suv39h1_B_12352 | |
| ACTTGTGCAGGGATGCTCCA | Suv39h1_B_12353 | |
| GGACACCTGAGGTTTCTCAA | Suv39h2_B_12346 | Suv39h2 |
| ACACTCGTCGCCAAAGACCG | Suv39h2_B_12347 | |
| GCGCCGTCAGGGTCTCCGCG | Suv39h2_B_12348 | |
| GCACGAGGTGCCTTATTCAA | Suv39h2_B_12349 | |
| AGTGCACACCAACAATCCCC | Suv420h1_1 | Suv420h1 |
| TTCCAGAGGACGGCACATAG | Suv420h1_2 | |
| GCAGAAACGCCGTTGAGAGG | Suv420h1_B_12342 | |
| GTTCTCACAGAGCTCCTTGG | Suv420h1_B_12343 | |
| ATATGCCTCGAGCTCCTCGA | Suv420h1_B_12344 | |
| TCTGCTGGACCGCCTCTCAA | Suv420h1_B_12345 | |
| GCTCACCACAGAATGGGTCA | Suv420h2_R1.Br_59412 | Suv420h2 |
| AGTGACTTGCTTCTATGGTG | Suv420h2_R1.Br_59413 | |
| CGGAGCTCATACTTGTCTAG | Suv420h2_R1.Br_59415 | |
| AGGAGCTGTAGACTTATCGT | Suz12_1 | Suz12 |
| AGTCCCTACTGGTAAAAAGC | Suz12_2 | |
| ATGGCGCCTCAGAAGCACGG | Suz12_3 | |
| CTTCTTCACCGGCAACACCG | Suz12_As_12334 | |
| GCGATGGCGCCTCAGAAGCA | Suz12_As_12335 | |
| GCTTCGGGCGGCAAATCCGG | Suz12_As_12336 | |
| GATTCCTAAGAGTAAAACCA | Syncrip_As_12157 | Syncrip |
| ACAGGCAGAGAGAAAAACAG | Syncrip_As_12158 | |
| GTTCAATGAAGACGGCGCAT | Syncrip_As_12159 | |
| TTCGTGGCAAAGGAGCCCCA | Tada1_As_11838 | Tada1 |
| GCCAAGCCTGGAAAACCCAA | Tada1_As_11839 | |

| | | |
|---|---|---|
| AGACTCCTCACACAGGACAA | Tada1_As_11840 | |
| GGGCGGCTGGACCAGCCGCG | Tada2b_As_11830 | Tada2b |
| GCTGGTAGCCGTGGTAGCGG | Tada2b_As_11831 | |
| CTGCGTGTACTGCTTGGCCG | Tada2b_As_11832 | |
| ATGTTGGTCGGGTTCTGCGG | Taf4a_As_11778 | Taf4a |
| GCTCCCATTCAAAGTTTGCG | Taf4a_As_11779 | |
| CGCCGGCTCTGTCATCCAGG | Taf4a_As_11780 | |
| TTAGAGGTAGAGTAATTCCA | Taf7_As_11754 | Taf7 |
| GTATAGGTCACCATCTACTG | Taf7_As_11755 | |
| ATATGCCGCTACGGTGAGGA | Taf7_As_11756 | |
| CGTTTGCTCACATCTCTTGG | Taf9b_As_11738 | Taf9b |
| GGTGATGGCACAGATCCTGA | Taf9b_As_11739 | |
| GTTAAGCAGTATCTATCAGG | Taf9b_As_11740 | |
| GCTGGTTACCTGATTCCTGG | Tbl1x_As_11274 | Tbl1x |
| AGCCACAGCCACAAGCACAG | Tbl1x_As_11275 | |
| GGCGGAGATCAGCATCAACG | Tbl1x_As_11276 | |
| GATGATAGAGATGAGTGCAG | Tbl1xr1_As_11270 | Tbl1xr1 |
| GGAAATCCCTTCCAATAAAG | Tbl1xr1_As_11271 | |
| AGATCCTTGCTGGTTAGTGG | Tbl1xr1_As_11272 | |
| GTCATTGGTGAAGGGTACAA | Tbp_As_11258 | Tbp |
| CATTGGTGAAGGGTACAAGG | Tbp_As_11259 | |
| AGATGTTGATTGCTGTACTG | Tbp_As_11260 | |
| AGTCGATTAGGAACCCACGA | Tcf12_1 | Tcf12 |
| CCAATGTCCAGCTTCCACCG | Tcf12_2 | |
| TCACTTGCTGTTCTAGACTA | Tcf12_3 | |
| GTACAAAGTCAAGTTCCCTG | Tdrd7_B_10864 | Tdrd7 |
| ACACTGGCCTCACATTTGCA | Tdrd7_B_10865 | |
| ACATACCATTGACTTCCGTA | Tdrd7_B_10866 | |
| TCAGCGCTGTTAAGAGACAG | Tdrd7_B_10867 | |
| ACAGCAGAGAGACCCAGACT | Tead1_1 | Tead1 |
| AGCCAGACGTACCTTAATGG | Tead1_2 | |

| | | |
|---|---|---|
| TAAGCCGATTGACAACGACG | Tead1_3 | |
| CACTCTGGAACATTCCATGG | Tead2_1 | Tead2 |
| CTGACAGGATGATCTTGCGA | Tead2_2 | |
| CTGGCCATCTACCCTCCCTG | Tead2_3 | |
| ACAAGGAAGAAAGCGTTCGG | Tead3_1 | Tead3 |
| AGCTGAGGACTGGAAAAACC | Tead3_2 | |
| GAACTTGTTCTGTAGAACGC | Tead3_3 | |
| TGTTGGGTACCTTCTCAGGA | Tet1_As_10684 | Tet1 |
| GCCCCATTAGATCTTACCCA | Tet1_As_10685 | |
| ATAATATCCAGGACGAGCCG | Tet1_As_10686 | |
| CGAAGCTTGCAAATTCCGGT | Tet2_As_10680 | Tet2 |
| GGCAATGTCAACATGCCAGG | Tet2_As_10681 | |
| AATACTATCCTAGTTCCGAC | Tet2_As_10682 | |
| GCTGCTCCAGTTCTGCCATA | Tet3_As_10676 | Tet3 |
| TTAGCTGCCTTGAATCTCCA | Tet3_As_10677 | |
| GGTACAGGCCAGGAGTTCCG | Tet3_As_10678 | |
| GCAGGATGAAACCCATACGT | Tex10_As_10672 | Tex10 |
| TTTGCCTGAACAACTTAAAG | Tex10_As_10673 | |
| CCTTGGCAGATGGATCCAGT | Tex10_As_10674 | |
| GAAGAGGAAGAGGCAGCCCG | Tfpt_As_10464 | Tfpt |
| TGAGAACTCTTCAAAGCCCA | Tfpt_As_10465 | |
| GATAACTCGGAGACTCCAGC | Tfpt_As_10466 | |
| AGAGCGTTCATGGTTCCGAG | Tgfbr1_1 | Tgfbr1 |
| ATTGTGTTACAAGAAAGCAT | Tgfbr1_2 | |
| CACGGTGGTGAATGACAGTG | Tgfbr1_3 | |
| CGGTTTGGAGAAGTTTGGCG | Tgfbr1_As_10420 | |
| CTTGATGGAACGGTTCAATG | Tgfbr1_As_10420 | |
| TCATCCAGTCGAGAAACCGG | Tgfbr1_As_10421 | |
| TGCACCATCTTCAAAAACAG | Tgfbr1_As_10421 | |
| AGGAGTCAATCCATCACGAG | Tgfbr1_As_10422 | |
| GATCGCCCTTTCATTTCAGA | Tgfbr1_As_10422 | |

| | | |
|---|---|---|
| TCCGGGATAGTGAACTTAAA | Tle1_As_9938 | Tle1 |
| GTCTCTGGACCGGATTAAAG | Tle1_As_9939 | |
| TTACTTCCTGAGACAGAAAT | Tle1_As_9940 | |
| GTACCCGCAGACGCGCCACC | Tle4_As_9926 | Tle4 |
| TGGAAGATGCAACAGAAGCG | Tle4_As_9927 | |
| AGTCCTGCACTGCTACCGAT | Tle4_As_9928 | |
| ACTGCTGCAGTGTCTCCAGT | Tonsl_As_8154 | Tonsl |
| TTTGAAGTAGTTGTTGCACA | Tonsl_As_8155 | |
| GTTGAGGGCGTCATGCAGGG | Tonsl_As_8156 | |
| ATTGTGGGTTTACCTCACCA | Top2a_As_8142 | Top2a |
| TGAACAGGTCAATCCCCGGT | Top2a_As_8143 | |
| GAAAAAGAAGAACAAGGGCG | Top2a_As_8144 | |
| TATGGAGTCTTCCCACTCAG | Top2b_As_8138 | Top2b |
| GCGCCCCGTCAGAGCCCGCG | Top2b_As_8139 | |
| TGTAGGGATGAACTGCAGGG | Top2b_As_8140 | |
| TCTTGGGGCCAGCAGAGTAA | Trim16_As_7638 | Trim16 |
| AATTGGATCTGATTGCCCCG | Trim16_As_7639 | |
| GGGCCTTCCGTACAGCTGCG | Trim16_As_7640 | |
| CCAGCGGGTGAAATACACCA | Trim28_B_7602 | Trim28 |
| TGAATTGCAGAAGGTGACCG | Trim28_B_7603 | |
| GCTCTCACAGAACAGCACGA | Trim28_B_7604 | |
| CGCCGCAGCGAATAATTCGG | Trim28_B_7605 | |
| AACAGATCGTCCATGCAGTG | Trp53_R1.Br_180821 | Trp53 |
| TCCACCCGGATAAGATGCTG | Trp53_R1.Br_180831 | |
| GAAGTCACAGCACATGACGG | Trp53_R1.Br_180841 | |
| GCTGCTTTCCAATTGCCCAG | Trrap_As_7122 | Trrap |
| GCAATGGTATTCATGATCAG | Trrap_As_7123 | |
| GATGGTGGCAAACAGAGCAG | Trrap_As_7124 | |
| AGCTCCTTATGAAACTGCGG | Tsfm_B_7057 | Tsfm |
| TCCTTACTCTCCCCAGGCGA | Tsfm_B_7058 | |
| GGGCGCGCGCGCTCACCGAG | Tsfm_B_7059 | |

| | | |
|---|---|---|
| CCATGGCTCACGTTTCACGC | Tsfm_B_7060 | |
| TCGTTCCTCCAACTTGACTC | Tyw5_As_6243 | Tyw5 |
| AGCAGCGTCTTCCGGTACCC | Tyw5_As_6244 | |
| CTTCCGGTACCCCGGCTGCG | Tyw5_As_6245 | |
| GTCACCCTAAGCCTACCCAG | Uba2_As_6203 | Uba2 |
| GTTGGGTTTGAAAGACCAGC | Uba2_As_6204 | |
| CCAGACACATCCTATTCACG | Uba2_As_6205 | |
| TCTCTTACCATCCTCAAACG | Ube2a_As_6127 | Ube2a |
| TTACAGGAAGATCCTCCGGC | Ube2a_As_6128 | |
| TCTTGAAGTCCCTCATGAGG | Ube2a_As_6129 | |
| CATGAGCCTCCTACGGGCCG | Ube2b_As_6123 | Ube2b |
| AATTACTTACCATCTTCAAA | Ube2b_As_6124 | |
| GGCTCATGCGGGATTTCAAG | Ube2b_As_6125 | |
| GGATCCGCTTCAGCGCCATG | Ube2d1_As_6111 | Ube2d1 |
| TCCACCTGCCCACTGCTCAG | Ube2d1_As_6112 | |
| CCACTGCTCAGCGGGACCCG | Ube2d1_As_6113 | |
| ACAATAATACTTGCCTTAGG | Ube2d3_As_6099 | Ube2d3 |
| CTTGCCTTAGGTGGTTTGAA | Ube2d3_As_6100 | |
| AATGACAGCCCATATCAAGG | Ube2d3_As_6101 | |
| GCTTACCTGCAGTTTGGCGG | Ube2e1_As_6087 | Ube2e1 |
| TTTCACAGTGCTGGTCCCAA | Ube2e1_As_6088 | |
| GTTGGAGGACGAAGATGACG | Ube2e1_As_6089 | |
| TTGCTCAGTGAGTCTCCAGA | Ubn1_As_5927 | Ubn1 |
| ACTAGAGCATCCTTGCTGCA | Ubn1_As_5928 | |
| GAGTTGGTAAAGAATATCCG | Ubn1_As_5929 | |
| TCTTGTTTCAGGTTGCCGAG | Uchl5_As_5803 | Uchl5 |
| TCTTGTTTCAGGTTGCCGAG | Uchl5_As_5803 | |
| GACCATGTCGAGCAATGCCG | Uchl5_As_5804 | |
| GACCATGTCGAGCAATGCCG | Uchl5_As_5804 | |
| TTTCTTCTACTTGGGCCCCT | Uchl5_As_5805 | |
| TTTCTTCTACTTGGGCCCCT | Uchl5_As_5805 | |

| | | |
|---|---|---|
| GGTCATGGCCAACTATAACG | Uhrf1_B_5625 | Uhrf1 |
| GATACCATGGGAGAACCTGG | Uhrf1_B_5626 | |
| GTACATGATGTCATCCTCCG | Uhrf1_B_5627 | |
| GATTGAGGAAGTGTTTCACG | Uhrf1_B_5628 | |
| TTAGGCGGATTATGAGAAGA | Uhrf2_As_5613 | Uhrf2 |
| GCTTCTGATGGACATTCACG | Uhrf2_As_5614 | |
| TTTCATCTACAGACATCACT | Uhrf2_As_5615 | |
| TATACCTTGCGCTCAATAGG | Usp11_As_5298 | Usp11 |
| GGACCGAGACACACAGCCCG | Usp11_As_5299 | |
| ACACTGGTATAAACAGTGGG | Usp11_As_5300 | |
| ATTCGCCTCCATCTGTACCA | Usp12_As_5294 | Usp12 |
| AATCTCTTTCTCTAATGCCG | Usp12_As_5295 | |
| ACAAGGTTGCGGAAAGAAAA | Usp12_As_5296 | |
| GCTAAAGTAAGAGGACCTGG | Usp16_As_5278 | Usp16 |
| TAGGTGCTACAAGTGTGACG | Usp16_As_5279 | |
| TAAGACAGAGCCAAACCGAA | Usp16_As_5280 | |
| ACAGGAGTCAGGGTGCCAGA | Usp21_As_5245 | Usp21 |
| GCTGCTCAAACACTGTAGCA | Usp21_As_5246 | |
| GCTCAGGCCGGAGTGCCAAG | Usp21_As_5247 | |
| CTCTGCCTCAGAATTCAAAA | Uty_As_5025 | Uty |
| CTAGAGGCGAGGGCGAAGAG | Uty_As_5026 | |
| TCTGCCTCAGAATTCAAAAG | Uty_As_5027 | |
| CATTCCGGTCAAAGTCACCA | Vdr_As_4861 | Vdr |
| GTACAGATCAGAGTTTGAGG | Vdr_As_4862 | |
| ATGACCTGTGAAGGCTGCAA | Vdr_As_4863 | |
| GGAATTAAAGTTCTACCAGA | Vrk1_As_3559 | Vrk1 |
| CTTACCACTTTCACAACACA | Vrk1_As_3560 | |
| TGAGCATGAGTACGTGCACG | Vrk1_As_3561 | |
| AGAGTTAGAGAAAGGGACGG | Wac_As_3415 | Wac |
| TTGAACTATGAAGTGCACTG | Wac_As_3416 | |
| AGACAGGGATTACAGAAGAG | Wac_As_3417 | |

| | | |
|---|---|---|
| TTACACACTGACAAGCAAGA | Wapal_R1.Br_3407 | Wapal |
| GTAAGATGAACCCTTACCCT | Wapal_R1.Br_3408 | |
| TCATCACTATCAAAACCAAA | Wapal_R1.Br_3409 | |
| AGAAATGCTGAAGCAGGCGC | Wbscr27_B_3331 | Wbscr27 |
| GCTTTCACCTGGTCATACTC | Wbscr27_B_3332 | |
| GCCCGGGTCGGAACCTCCCA | Wbscr27_B_3333 | |
| GAGCCCAGTCGTCATAGAAG | Wbscr27_B_3334 | |
| GGACAGACAGATATTCCTTT | Wdr48_As_3187 | Wdr48 |
| TCAGAGTGTTCACATCCCAA | Wdr48_As_3188 | |
| ACCGGCAGAACACAGCCGGG | Wdr48_As_3189 | |
| ATTGGGGCTGAACTTCACAG | Wdr5_As_3183 | Wdr5 |
| GGAGAAGAAGCCAGAGACAG | Wdr5_As_3184 | |
| CTATGCCCTGAAGTTCACCC | Wdr5_As_3185 | |
| GTACAGTAAGAAGTACGGTG | Wdr82_As_3095 | Wdr82 |
| TAGGGCCTCATGCATCTACA | Wdr82_As_3096 | |
| GCTGTCGGTCAGCTTCATGG | Wdr82_As_3097 | |
| ACTGTGGCCAGATTGTGTGG | Wsb2_As_2775 | Wsb2 |
| ATTTGGGACCTGAATAAGCA | Wsb2_As_2776 | |
| GCAAGAATGACCCAAAAGGA | Wsb2_As_2777 | |
| CAGCACGGTCACTTTCGACG | Wt1_1 | Wt1 |
| CCCGCACGTCGGAACCCATG | Wt1_2 | |
| GGTCATGCATTCAAGCTGGG | Wt1_3 | |
| GCGACAAGAAGAGCCCCACC | Yaf2_As_2526 | Yaf2 |
| ATGATGTGCGATGTGCGGAA | Yaf2_As_2527 | |
| CACCTTCCGGAACAGCGCCG | Yaf2_As_2528 | |
| ATAAACGATTGGCTTAACGA | Yeats4_As_2486 | Yeats4 |
| AATTTGGACCTGACTCCGGC | Yeats4_As_2487 | |
| GCAACATTGCCATAAACGAT | Yeats4_As_2488 | |
| AGAGAGATCAGAAGGAAGAA | Ythdc1_As_2402 | Ythdc1 |
| GCCAAGTCTCCTACACCAGA | Ythdc1_As_2403 | |
| CATGGCGGCCGACAGCCGGG | Ythdc1_As_2404 | |

| | | |
|---|---|---|
| ATAGGTAGTGAGATACGGGA | Ythdf1_As_2394 | Ythdf1 |
| GGTAAGGATCACTCATCGAG | Ythdf1_As_2395 | |
| GTAAGTGTAACTGCTCCCAT | Ythdf1_As_2396 | |
| CAGCCGAAACTTAAACCCAA | Ythdf3_As_2386 | Ythdf3 |
| GGTGCTGCACTGCTAACAGG | Ythdf3_As_2387 | |
| GCTCTCCCAAGAGAACTAGG | Ythdf3_As_2388 | |
| AATCAATGAAGAAAGTAGCA | Ywhae_As_2378 | Ywhae |
| GAGCAGGCCGAGCGATACGA | Ywhae_As_2379 | |
| TTAACAATCTCACCATTTGC | Ywhae_As_2380 | |
| GCTCCCACTTCTTATTCCCC | Yy1_As_2358 | Yy1 |
| CGAGGTGGAGACCATCCCGG | Yy1_As_2359 | |
| CGACACCCTCTACATCGCCA | Yy1_As_2360 | |
| GCAACTCTAAGCCAAACACA | Zbtb33_As_2234 | Zbtb33 |
| ATATCAGGTACAGAACAGGA | Zbtb33_As_2235 | |
| AACTCCTTGAATGAACAGCG | Zbtb33_As_2236 | |
| TGTGCAGCCTCAACGAGCAG | Zbtb7c_As_2158 | Zbtb7c |
| GGACTATTCGGACACACCCA | Zbtb7c_As_2159 | |
| TTTCCCTAACCACAGCAGCG | Zbtb7c_As_2160 | |
| GCTCGGTTGTGACGTGAGAG | Zfp185_1 | Zfp185 |
| TGGACAAAGTGACCCAACTG | Zfp185_2 | |
| TGTCAGGTCTCCAAACGCAA | Zfp185_3 | |
| TCAGGAGAAGAGCATGGCCG | Zfp217_As_1662 | Zfp217 |
| CATCACAGCTCAACTCCTCG | Zfp217_As_1663 | |
| AGAGGAACCCACGTCCCCGA | Zfp217_As_1664 | |
| AGAGTGACCGAGTGCCTGCG | Zfp36l1_1 | Zfp36l1 |
| CCCGTACGGGCAAAAGCCGA | Zfp36l1_2 | |
| TAGGGGAGTCTGAGCCACTG | Zfp36l1_3 | |
| AGAAGGTCCTTACCTCGATG | Zfp503_1 | Zfp503 |
| CGGTTTGTAGGGAGATACCG | Zfp503_2 | |
| GGGTCGAGTGCCACACCGTG | Zfp503_3 | |
| AGATCTGCAAGATAAGAACG | Zfp57_As_1139 | Zfp57 |

| | | |
|---|---|---|
| AGTTTCCTGTGAGGTCCCGG | Zfp57_As_1140 | |
| TATAACCTAAATGAGCACGT | Zfp57_As_1141 | |
| ATACATTCAGGGAGCCCCCA | Zfp592_As_1107 | Zfp592 |
| ATCGGGGGAAATCAAACGGA | Zfp592_As_1108 | |
| ACAAAGATCTGGATTCCAGT | Zfp592_As_1109 | |
| ATACCAAAGCAAGACACGTG | Zfp68_1 | Zfp68 |
| GTTAGACTGTTTGCTCTCAG | Zfp68_2 | |
| TGTATAGACTAACTTGCTCG | Zfp68_3 | |
| TCACCTGGTGACAAACCACA | Zfpm1_B_4681 | Zfpm1 |
| GTGCACACAGACACGCTCAG | Zfpm1_B_4691 | |
| GCACACTTACCTTCTCTGGG | Zfpm1_B_4701 | |
| AGTAAGGCCGAAGTCACCAA | Zfpm1_B_471 | |
| GTTAGCACAGGTGATAACTG | Zhx1_As_392 | Zhx1 |
| TTCCTACACACATTTCCACA | Zhx1_As_393 | |
| CATTGCCCTGACAGTGGCAG | Zhx1_As_394 | |
| AAAGGTGCAGAAGATGGAGG | Zmym2_As_272 | Zmym2 |
| TGTGATCCCGAACTTCCTTC | Zmym2_As_273 | |
| ATATTGCGAATACTGCCAAG | Zmym2_As_274 | |
| GCTAGTCCCCCTTCACCTGA | Zmym3_As_268 | Zmym3 |
| CTCCATCCAGAACTCCTCCA | Zmym3_As_269 | |
| ATGCTCCTCCTCCTGCCCAG | Zmym3_As_270 | |
| ACAAAATCCCACATACGCAA | Zmynd11_As_248 | Zmynd11 |
| GCTTCTATCTGTCAAACGCA | Zmynd11_As_249 | |
| GTAAGCATCCTATGTACCGG | Zmynd11_As_250 | |
| CACTTAGCGTGATAAACCCG | Zmynd8_As_232 | Zmynd8 |
| AGAGGATGTTTATACAGCCG | Zmynd8_As_233 | |
| AGACTGACATCGGAGCCAGA | Zmynd8_As_234 | |
| TCCGGAAGCGAAGTTTAAAG | Znhit1_As_220 | Znhit1 |
| CCGAAAACCCGAGATGCAGA | Znhit1_As_221 | |
| TCTGCATCATCATCAAACTG | Znhit1_As_222 | |
| ACACCCTGAGAACTTCACCT | Zzz3_As_1 | Zzz3 |

| | | |
|---|---|---|
| CCTCCTCTCACAACTCTCTA | Zzz3_As_2 | |
| GCAGTAGTAGAAGAAATCAC | Zzz3_As_3 | |
| GCGAGGTATTCGGCTCCGCG | NonTargetingControlGuideForMouse_0001_MGLibA_66406 | NTC |
| GCTTTCACGGAGGTTCGACG | NonTargetingControlGuideForMouse_0002_MGLibA_66407 | |
| ATGTTGCAGTTCGGCTCGAT | NonTargetingControlGuideForMouse_0003_MGLibA_66408 | |
| ACGTGTAAGGCGAACGCCTT | NonTargetingControlGuideForMouse_0004_MGLibA_66409 | |
| GACTCCGGGTACTAAATGTC | NonTargetingControlGuideForMouse_0005_MGLibA_66410 | |
| CCGCGCCGTTAGGGAACGAG | NonTargetingControlGuideForMouse_0006_MGLibA_66411 | |
| ATTGTTCGACCGTCTACGGG | NonTargetingControlGuideForMouse_0007_MGLibA_66412 | |
| ACCCATCGGGTGCGATATGG | NonTargetingControlGuideForMouse_0008_MGLibA_66413 | |
| CGGGCGTCACCTGCTAGTAA | NonTargetingControlGuideForMouse_0009_MGLibA_66414 | |
| GCTTCTACTCGCAACGTATT | NonTargetingControlGuideForMouse_0010_MGLibA_66415 | |
| TACAGTTATACGTCGCGGTG | NonTargetingControlGuideForMouse_0011_MGLibA_66416 | |
| AAGCACAAGAACGGTCCGCC | NonTargetingControlGuideForMouse_0012_MGLibA_66417 | |
| CAGCCACCGCACCGGCGTAA | NonTargetingControlGuideForMouse_0013_MGLibA_66418 | |
| GTCAAGCCGAACGCTGCCGG | NonTargetingControlGuideForMouse_0014_MGLibA_66419 | |
| CCTTAGACCGGGTGTACCTC | NonTargetingControlGuideForMouse_0015_MGLibA_66420 | |
| AAGTCTATGCGGGGCTCGTA | NonTargetingControlGuideForMouse_0016_MGLibA_66421 | |
| TTGTCAACTTCGGCCAACGC | NonTargetingControlGuideForMouse_0017_MGLibA_66422 | |
| ATAGATGTCTACGCGCCGTT | NonTargetingControlGuideForMouse_0018_MGLibA_66423 | |
| CTCGGGCTATTCAGCGATAG | NonTargetingControlGuideForMouse_0019_MGLibA_66424 | |
| GCGGTTACCGCGAAAACCAT | NonTargetingControlGuideForMouse_0020_MGLibA_66425 | |
| ACCAACGCTACGATCCCGGA | NonTargetingControlGuideForMouse_0021_MGLibA_66426 | |
| CCCTATATGCGAGATCCATA | NonTargetingControlGuideForMouse_0022_MGLibA_66427 | |
| AGAAAGGCACGTGCGACGTC | NonTargetingControlGuideForMouse_0023_MGLibA_66428 | |
| GACCAACCTTACGGTAACTC | NonTargetingControlGuideForMouse_0024_MGLibA_66429 | |
| ATTATTCCTCCGGATGACGA | NonTargetingControlGuideForMouse_0025_MGLibA_66430 | |
| TCTCAGTTCGTAGCGAACGA | NonTargetingControlGuideForMouse_0026_MGLibA_66431 | |
| TCCCGGGAGGTACGGTGTAC | NonTargetingControlGuideForMouse_0027_MGLibA_66432 | |
| GGGGATGGCCTTACGTCGCG | NonTargetingControlGuideForMouse_0028_MGLibA_66433 | |
| GTATCCTCCTTACGGCCCGT | NonTargetingControlGuideForMouse_0029_MGLibA_66434 | |

| | |
|---|---|
| GCTCGGACCTTTTAGACGTC | NonTargetingControlGuideForMouse_0030_MGLibA_66435 |
| GTGGTTACGTTAACGACTAC | NonTargetingControlGuideForMouse_0031_R1.Br_66436 |
| TGCAACGATGGTTACGGTAC | NonTargetingControlGuideForMouse_0032_R1.Br_66437 |
| AACGGGCGCAATACCCTTTT | NonTargetingControlGuideForMouse_0033_R1.Br_66438 |
| TCGTGTCTAGCTATCGAGTG | NonTargetingControlGuideForMouse_0034_R1.Br_66439 |
| GACGTCTAATTTCTGGCCGT | NonTargetingControlGuideForMouse_0035_R1.Br_66440 |
| TAGTCCTAGTTAGATTCGCG | NonTargetingControlGuideForMouse_0036_R1.Br_66441 |
| AAGGCCTTAACACGTCGACC | NonTargetingControlGuideForMouse_0037_R1.Br_66442 |
| ATTCGTGCATCGCGGGGTTT | NonTargetingControlGuideForMouse_0038_R1.Br_66443 |
| CACCTCGCGTCATATCACTA | NonTargetingControlGuideForMouse_0039_R1.Br_66444 |
| GACCTCGCAATTGAGCGCTC | NonTargetingControlGuideForMouse_0040_R1.Br_66445 |
| TGTATCCACCGTGACCCGGT | NonTargetingControlGuideForMouse_0041_R1.Br_66446 |
| GATCTTACCACTCGTCGTAG | NonTargetingControlGuideForMouse_0042_R1.Br_66447 |
| TTCGCCGGCGACGAAGTGCA | NonTargetingControlGuideForMouse_0043_R1.Br_66448 |
| AGCACTAGGATCGCGGCCTT | NonTargetingControlGuideForMouse_0044_R1.Br_66449 |
| GCGTATCTACCCTACCGCCG | NonTargetingControlGuideForMouse_0045_R1.Br_66450 |
| GCGCGAGGGCACCGACAAGT | NonTargetingControlGuideForMouse_0046_R2.Br_66451 |
| CGCGTATATGTCACACGGCA | NonTargetingControlGuideForMouse_0047_R2.Br_66452 |
| TAAAACCGATCACGATACGA | NonTargetingControlGuideForMouse_0048_R2.Br_66453 |
| TCCTGCGCGATGACCGTCGG | NonTargetingControlGuideForMouse_0049_R2.Br_66454 |
| TGGCCCACAAGGTGCGATAT | NonTargetingControlGuideForMouse_0050_R2.Br_66455 |
| GGGGTAGGCCTAATTACGGA | NonTargetingControlGuideForMouse_0051_R2.Br_66456 |
| CGCTTCGTCTCTCGCAAACA | NonTargetingControlGuideForMouse_0052_R2.Br_66457 |
| TTATGTGCCTCTCGGCGCAT | NonTargetingControlGuideForMouse_0053_R2.Br_66458 |
| CCGTTCTGACGACGCTAAAG | NonTargetingControlGuideForMouse_0054_R2.Br_66459 |
| AAACTCCCGTGTCAACCGAT | NonTargetingControlGuideForMouse_0055_R2.Br_66460 |
| GTTTGCGAGTCAAAGTACGC | NonTargetingControlGuideForMouse_0056_R2.Br_66461 |
| GGGTGCACACGCCGGCCTAT | NonTargetingControlGuideForMouse_0057_R2.Br_66462 |
| CTTTATACCGCGCGTCGGCA | NonTargetingControlGuideForMouse_0058_R2.Br_66463 |
| TCAAACGCCCGGGCGCCCCA | NonTargetingControlGuideForMouse_0059_R2.Br_66464 |
| GACCGGTGTGTTTACGCGTG | NonTargetingControlGuideForMouse_0060_R2.Br_66465 |
