## Supplementary Tables 2-7 for "Identification of epigenetic regulators of fibrotic transformation in cardiac fibroblasts through bulk and single-cell CRISPR screens"

**Supplementary Table 2. List of antibodies used for flow cytometry and FACS analyses.**

| NAME | CLONE | SUPPLIER | DILUTION |
|---|---|---|---|
| CD31-FITC | 390 | BioLegend | 1:200 |
| CD45-PerCP/Cy5.5 | 30-F11 | BD Pharmingen | 1:200 |
| Anti-feeder cells-APC | MEFSK4 | Miltenyi | 1:50 |
| CD146-PE/Cy7 | ME-9F1 | BioLegend | 1:100 |
| CD44-APC | KM201 | Invitrogen | 1:200 |
| CD90-PerCP/Cy5.5 | 30-H12 | BioLegend | 1:200 |
| PDGFRa-PE | APA5 | BioLegend | 1:200 |
| aSMA-PE | 1A4 | Sigma | 1:100 |
| CD29-PE/Cy7 | HMb1-1 | BioLegend | 1:200 |
| SCA1-FITC | E13-161.7 | BD Bioscience | 1:300 |
| CD34-FITC | RAM34 | BD Bioscience | 1:100 |
| CD73-PE | Ty/11.8 | BioLegend | 1:100 |
| NG2-PE | 1E6.4 | Miltenyi | 1:50 |

**Supplementary Table 3. List of primers used for qPCR analyses in mouse samples.**

| GENE | SEQUENCE Fw (5’ to 3’) | SEQUENCE Rv (5’to 3’) |
|---|---|---|
| *Acta2* | ATGGAGTCAGCGGGCATC | CGTTCTGGAGGGGCAATGAT |
| *Ltbp2* | ACCATCACCACCTCCACTC | CTCGAGTCCTGAAGGCAAAG |
| *Postn* | TATGCTCTGCTGCTGCTGTT | CTTTTTGGTGCCCAGAATTT |
| *Col1a1* | GCATCGCCAAGAAGACATCC | CAGATCAAGCATACCTCGGG |
| *Col1a2* | AGGCAGAGATGGTGTTGATG | AGGGCCAGATGAAACTCCTT |
| *Lox* | TGCCAACACACAGAGGAGAG | CTGCCGCATAGGTGTCATAA |
| *Gapdh* | ACTTTGTCAAGCTCATTTCC | TGCAGCGAACTTTATTGATG |
| *Rpl4* | GCCGCTGGTGGTTGAAGATAA | CGTCGGTTTCTCATTTTGCCC |

**Supplementary Table 4. List of primers used for qPCR analyses in human samples.**

| GENE | SEQUENCE Fw (5' to 3') | SEQUENCE Rv (5'to 3') | PROBE (5'to 3') |
|---|---|---|---|
| *Col1a1* | TTCTGTACGCAGGTGATTG G | GACATGTTCAGCTTTGTG GAC | /56-FAM/TCGAGGGCC/ZEN/AAG ACGAAGACATC/3IABkFQ/ |
| *Lox* | AGTGGCTAAACTCATCCAT ACTG | GCTCAGATTTCCCAAAG AGT | /56-FAM/TGACAACTG/ZEN/TGC CATTCCCAGGA/3IABkFQ/ |
| *Acta2* | AGAGTTACGAGTTGCCTG ATG | CTGTTGTAGGTGGTTTCA TGGA | /56-FAM/AGACCCTGT/ZEN/TCCA GCCATCCTTC/3IABkFQ/ |
| *Cthrc1* | CAAGCAGTGTTCATGGAG TTC | CTTGACAGCATGCATTTC TGC | /56-FAM/TTGTTCAGT/ZEN/GGCT CACTTCGGCTAA/3IABkFQ/ |
| *Gapdh* | ACATCGCTCAGACACCAT G | TGTAGTTGAGGTCAATG AAGGG | /56-FAM/AAGGTCGGA/ZEN/GTC AACGGATTTGGTC/3IABkFQ/ |

**Supplementary Table 5. Primer sequences used in this study.**

| NAME | SEQUENCE (5' to 3') | USE |
|---|---|---|
| U6 Forward | TAATTTCTTGGGTAGTTTGCAG | Sanger sequencing of sgRNAs |
| Read1-U6 | CTACACGACGCTCTTCCGATCTGTGGAAAGGACGAAACACCG | CRISPR screen |
| Read2-scaffold | AGACGTGTGCTCTTCCGATCTGCTGTCCCTGTAATAAACCCG | CRISPR screen |
| P7-index | CAAGCAGAAGACGGCATACGAGAT[i7]GTGACTGGAGTTCAGA CGTGTGCTCTTCCGATCT | CRISPR screen, INDEL-seq |
| P5-index | AATGATACGGCGACCACCGAGATCTACAC[i5]ACACTCTTTCCC TACACGACGCTCTTCCGATCT | CRISPR screen, INDEL-seq |
| P5-i5-Read1 | AATGATACGGCGACCACCGAGATCTACAC[i5]TCGTCGGCAGC GTCAGATGTGTAT | ATAC-seq |
| P7-i7-Read2 | CAAGCAGAAGACGGCATACGAGAT[i7]GTCTCGTGGGCTCGGA GATGTG | ATAC-seq |
| Read1-gDNA_*Wdr82* | ACACTCTTTCCCTACACGACGCTCTTCCGATCTTGCCTTCAGCA GCAGCAG | INDEL-seq |
| Read2-gDNA_*Wdr82* | GTGACTGGAGTTCAGACGTGTGCTCTTCCGATCTGCACGATGG AGTCATCATCG | INDEL-seq |
| Read1-gDNA_*Tgfbr1* | ACACTCTTTCCCTACACGACGCTCTTCCGATCTGATGGTCTATA TCTGCCATA | INDEL-seq |
| Read2-gDNA_*Tgfbr1* | GTGACTGGAGTTCAGACGTGTGCTCTTCCGATCTGCCCCAACC AACTAATTA | INDEL-seq |
| Read1-gDNA_*Smad4* | ACACTCTTTCCCTACACGACGCTCTTCCGATCTATGTAAAGGT GAAGGTGACG | INDEL-seq |
| Read2-gDNA_*Smad4* | GTGACTGGAGTTCAGACGTGTGCTCTTCCGATCTCAGGCAACA GAGGGTTTTAA | INDEL-seq |
| Read1-gDNA_*Srcap* | ACACTCTTTCCCTACACGACGCTCTTCCGATCTCCATCTTTTTT CCCTTTCCT | INDEL-seq |
| Read2-gDNA_*Srcap* | GTGACTGGAGTTCAGACGTGTGCTCTTCCGATCTTTAGGCACC ACCTATAGATG | INDEL-seq |
| Read1-gDNA_*Kat8* | ACACTCTTTCCCTACACGACGCTCTTCCGATCTGCGCGTGGCG AACCAGAAGT | INDEL-seq |
| Read2-gDNA_*Kat8* | GTGACTGGAGTTCAGACGTGTGCTCTTCCGATCTAGTGGGCTT CTCAGCTCCCG | INDEL-seq |
| Read1-gDNA_*Hcfc1* | ACACTCTTTCCCTACACGACGCTCTTCCGATCTTTTGGGGGTCT GGCCAATGA | INDEL-seq |
| Read2-gDNA_*Hcfc1* | GTGACTGGAGTTCAGACGTGTGCTCTTCCGATCTAGACCTACC CAGGGGTTCAG | INDEL-seq |
| Read1-gDNA_*Kat5* | ACACTCTTTCCCTACACGACGCTCTTCCGATCTATGACTGGCA GTCTGGTGTC | INDEL-seq |
| Read2-gDNA_*Kat5* | GTGACTGGAGTTCAGACGTGTGCTCTTCCGATCTTCTGGATAG GCAGCCTCCCA | INDEL-seq |
| Read1-gDNA_*KAT5* | ACACTCTTTCCCTACACGACGCTCTTCCGATCTTGCTATTCTCA TAGCCCTGG | INDEL-seq |
| Read2-gDNA_*KAT5* | GTGACTGGAGTTCAGACGTGTGCTCTTCCGATCTTCTTGGGGA ACTGGATCTTC | INDEL-seq |

**Supplementary Table 6. List of primary and secondary antibodies used for Western Blot.**

| PRIMARY ANTIBODIES | | | |
|---|---|---|---|
| **NAME** | **CLONE** | **SUPPLIER** | **DILUTION** |
| SMAD4 | D3R4N | Cell Signaling, 46535 | 1:1000 |
| TGFBR1 | pAb | Invitrogen, PA5-32631 | 1:500 |
| KAT5 | pAb | Abcam, ab23886 | 1:500 |
| **SECONDARY ANTIBODIES** | | | |
| **NAME** | **CLONE** | **SUPPLIER** | **DILUTION** |
| Donkey HRP-conjugated anti-rabbit | pAb | GE Healthcare, NA934 | 1:5000 |

**Supplementary Table 7. sgRNAs sequences used for human CRISPR/Cas9.**

| sgRNA sequence | sgRNA ID | Gene ID |
|---|---|---|
| CTGAGCGTGAAGGACATCAG | KAT5_H2 | *KAT5* |
| AGTACCCCTAGGTATGGGGA | NTC_H2 | Non-targeting control (NTC) |
